# Deep Lipidomics and Molecular Imaging of Unsaturated Lipid Isomers: A Universal Strategy Initiated by mCPBA Epoxidation

**DOI:** 10.1101/649103

**Authors:** Ting-Hao Kuo, Hsin-Hsiang Chung, Hsin-Yuan Chang, Chiao-Wei Lin, Ming-Yang Wang, Tang-Long Shen, Cheng-Chih Hsu

**Affiliations:** Department of Chemistry, National Taiwan University, Taipei 10617, Taiwan; Department of Animal Science and Technology, National Taiwan University, Taipei 10617, Taiwan; Department of Plant Pathology and Microbiology, National Taiwan University, Taipei 10617, Taiwan; National Taiwan University Hospital and College of Medicine, Taipei 100, Taiwan

## Abstract

Cellular lipidome is highly regulated through lipogenesis, rendering diverse double-bond positional isomers (C=C isomer) of a given unsaturated lipid species. In recent years, there are increasing reports indicating the physiological roles of C=C isomer compositions associated with diseases, while the biochemistry has not been fully understood due to the challenge in characterizing lipid isomers inherent to conventional mass spectrometry-based lipidomics. To address this challenge, we reported a universal, user-friendly, derivatization-based strategy, **MELDI** (**m**CPBA **E**poxidation for **L**ipid **D**ouble-bond **I**dentification), which enables both large-scale identification and spatial mapping of biological C=C isomers using commercial mass spectrometers without any instrument modification. With the developed liquid-chromatography mass spectrometry (LC-MS) lipidomics workflow, we elucidated more than 100 isomers among mono- and poly-unsaturated fatty acids and glycerophospholipids in both human serum, where novel isomers of low abundance were unambiguously quantified for the first time. The capability of MELDI-LC-MS in lipidome analysis was further demonstrated using the differentiated 3T3-L1 adipocytes, providing an insight into the cellular lipid reprogramming upon stearoyl-coenzyme A desaturase 1 (SCD1) inhibition. Finally, we highlighted the versatility of MELDI coupled with mass spectrometry imaging to spatially resolve cancer-associated alteration of lipid isomers in a metastatic mouse tissue section. Our results suggested that MELDI will contribute to current lipidomics pipelines with a deeper level of structural information, allowing us to investigate underlying lipid biochemistry.

## INTRODUCTION

Lipids are essential biomolecules that serve as players of crucial cellular functions, including energy homeostasis, cell signaling, and organ protection. These functions are highly regulated through coordinated lipid anabolism and catabolism.^1^ The imbalance in lipid metabolism thus leads to alterations in cellular environment, and further associates with progression of diseases, such as cancer,^2^ Alzheimer’s disease,^3^ and cardiovascular disease.^4^ To fully elucidate the underlying networks involved in lipid metabolism, lipidomics has emerged as a distinct discipline that aims for comprehensive analysis of lipids in biological entities.^5^ In the past two decades, the field of lipidomics has been driven by the development of mass spectrometry (MS) technology, as it provides rich information of structures and quantities of lipid molecules in a complex mixture at high sensitivity and throughput.^5–6^ Nowadays, biological lipid profiles can be routinely obtained via versatile MS-based approaches, which paves an innovative way in tissue imaging^7^ and biomarker discovery.^8^

Lipid structures are extremely diverse, in principle rendering over 100,000 unique species by theoretical estimation.^9^ The exact number of lipid molecules is even larger due to the presence of structural isomers. One of the significant challenges in lipidomics analysis is to obtain complete structural characterization of lipid molecules.^10^ Taking glycerophospholipid (GPL) for example, the level of structural-defined molecule requires annotations of the lipid head group, the alkyl chain length, the desaturation degree, the position of carbon-carbon double bonds (C=C), the substitutional positions of alkyl chains to the glycerol backbone (*i.e. sn-*1 and *sn-2* linkages), and the chirality. Current tandem mass spectrometry (MS/MS) techniques provide the structural annotation at the level of alkyl/alkyl chain composition in routine chromatography-based lipidome analysis as well as molecular imaging.^11^ Nonetheless, annotation of C=C positions is still obscure in a vast majority of lipidomic studies,^12–13^ where each lipid species is reported as the summed combinations of multiple C=C positional isomers (C=C isomers). This problem is resulted from that C=C isomers yield near-identical MS/MS fragments upon collision-induced dissociation (CID), the most popular MS/MS technique equipped in commercial mass spectrometers. In addition, C=C isomers are not easily discriminated using traditional separation techniques such as liquid chromatography (LC) and gas chromatography (GC).^10,14^ Ion mobility spectrometry (IMS) allows separating of lipid isomers to some degree,^15–16^ while precise identification of C=C positions still relies on the database that may take years to establish. As each unsaturated lipid isomer represents a distinct pathway in cellular lipogenesis,^17^ such ambiguity in structural annotations may restrict the mining of lipid biology.

In the most recent years, there have been a booming development of MS-based methods to address this challenge. First, using newly-developed ion activation techniques, specific ions indicative to the C=C position can be generated during fragmentation. These techniques, including ultraviolet-photodissociation (UVPD),^18–22^ electron impact excitation of ions from organics (EIEIO)^23–24^ and ozone-induced dissociation (OzID),^25–27^ are compatible with mass spectrometry imaging (MSI) platforms,^22,27^ but usually require intense modification to commercial mass spectrometers. Alternatively, conducting chemical derivatization specific to C=C bonds enables the bond cleavage prior to MS,^28^ or makes the modified structure labile upon CID^29–32^ so as to generate diagnostic ions indicative to C=C positions. Recently, one of the derivatization methods, *Paternò–Büchi* (PB) photochemical reaction, has been applied in combination with LC-MS for a large-scale analysis of lipid isomers,^33^ and with MSI for lipid isomer mapping.^34^ Nonetheless, these derivatization-based methods still require, although gentle, modification on ionization sources to carry out lipid derivatization.

Bearing these in mind, we were thus motivated to develop a method with universal accessibility as well as compatibility with commercial mass spectrometers, so as to make the study of unsaturated lipid isomers in a handy fashion. Here we describe such an innovative, derivatization-based method, **MELDI** – (**m**CPBA **E**poxidation for **L**ipid **D**ouble-bond **I**dentification) (**Fig. 1a**). More specifically, MELDI utilizes *meta*-chloroperoxybenzoic acid (mCPBA), a commonly used epoxidation reagent in organic synthesis,^35^ to derivatize a C=C bond into an epoxide structure, which is labile upon low-energy CID. Subsequently, the affirmative C=C position diagnosis is easily achieved by the diagnostic ions generated via CID in negative ion mode. Recently, the utility of mCPBA epoxidation was also applied to positive-ionization shotgun analysis.^36^ In this study, we comprehensively explore the ability of MELDI in lipidomics and molecular imaging of multiple classes of mono- and poly-unsaturated lipids. Such versatile method allowed us to achieve deep lipidomic interrogations and molecular imaging in various biological systems, including human serum, mouse adipocytes, and cancer tissue sections. These results suggested that C=C isomers are important to cellular lipid homeostasis and thus serve as potential disease biomarkers.

**Figure 1.**
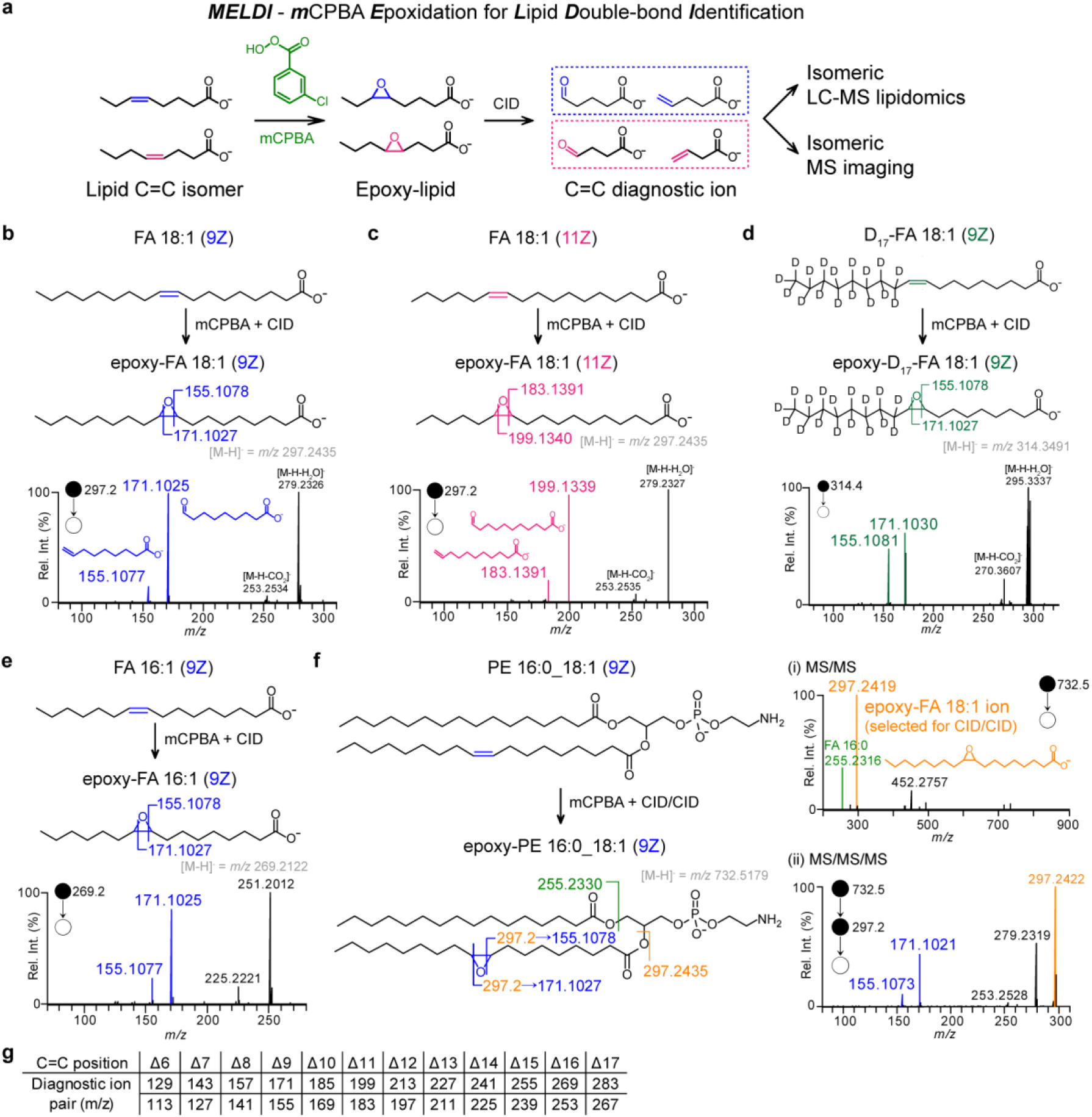
MELDI (mCPBA Epoxidation for Lipid Double-bond Identification) to pinpoint C=C positions of unsaturated lipids. **(a)** Schematic representation of MELDI. The C=C bond of an unsaturated lipid is epoxidized by mCPBA to form an epoxide, which is labile upon CID in negative ionization mode, so as to yield abundant diagnostic ions for identifying the C=C position. The FT-MS/MS spectra and CID schemes of **(b)** FA 18:1 (9Z) **(c)** FA 18:1 (11Z) **(d)** D_17_-FA 18:1 (9Z) **(e)** FA 16:1 (9Z), where the C=C diagnostic ion pairs generated upon CID of the epoxidized lipid were indicated. **(f)** For PE 16:0_18:1 (9Z), its C=C position was elucidated by MS^3^ of the epoxy-FA 18:1 acyl chain ion (*m/z* 297.24) from the MS/MS spectrum of the epoxidized lipid parent ion ([epoxyM-H]^−^ = *m/z* 732.52). **(g)** List of C=C diagnostic ions for epoxidized mono-unsaturated lipids.

## Results and Discussion

### Pinpoint C=C positions in unsaturated lipids by MELDI

The utility of MELDI to pinpoint lipid C=C positions was systematically investigated using a series of commercial standard mono-unsaturated lipids by high-resolution Fourier transform (FT) MS/MS. First, we showed that for two monounsaturated fatty acid (MOFA) C=C isomers, FA 18:1 (9Z) and (11Z), after in-solution epoxidation with mCPBA and subjected to electrospray ionization (ESI) MS/MS analysis in negative ionization mode, distinguishable CID spectra of their dehydrated parent ions ([M-H]^−^ = *m/z* 297.2435) were obtained (**Fig. 1b and 1c**), where their C=C positions were identified by the abundant C=C diagnostic ion pairs (*m/z* 155.1077 and 171.1025 for Δ9; *m/z* 183.1391 and 199.1339 for Δ11). The epoxide ring breaking upon CID was also confirmed by a deuterium labeled standard, D_17_-FA 18:1 (9Z) (**Fig 1d**). For another MOFA, FA 16:1 (9Z), with a shorter carbon chain and a C=C bond at Δ9, the CID spectrum of its epoxide showed an identical diagnostic ion pair as in FA 18:1 (9Z) (**Fig. 1e**). In addition to FAs, the C=C position of MOFA in a GPL molecule can be similarly identified through sequential CID (CID/CID) of the epoxy-FA daughter ion in the MS/MS spectra. Such strategy was further demonstrated by various subclasses of GPL, including phosphatidylethanolamine (PE) (**Fig. 1f**), phosphatidylserine (PS) (**Fig. S1a**), phosphatidylcholine (PC) (**Fig. S1b**), phosphatidic acid (PA) (**Fig. S1c**). In summary, the C=C diagnostic ions generated from MOFA-derived lipid epoxides are predictable and merely follow Δ positions regardless of fatty acyl chain lengths (**Fig. 1g**).

Notably, in addition to allowing the CID-based diagnosis of C=C positions, mCPBA epoxidation also enhanced chromatographic separation of unsaturated lipids in reverse-phase (RP) LC. This was demonstrated by an isomeric mixture of FA 18:1 (9Z) and (11Z). As shown in **Fig 2a**, the isomers were not separated in RPLC-MS before derivatization, as shown by a co-eluted peak. The same isomeric mixture was then derivatized with excess mCPBA and analyzed with LC-MS/MS. Remarkably, most of the non-derivatized FA isomers were converted into the epoxides, as shown by two distinguishable peaks of epoxy-FA 18:1 (**Fig. 2b**), allowing subsequent MS/MS analysis to determine their C=C positions. The results implied (i) the epoxide yield was regardless of C=C positions, referred by their similar peak area of the epoxides, (ii) epoxidation of an unsaturated lipid increased its hydrophilicity, resulting in the earlier elution in RPLC (iii) difference in the affinity with the column between C=C isomers was enhanced after epoxidation, as indicated by two separated peaks. On the other hand, for epoxidized geometric isomers in either E- or Z- conformation (*e.g.* epoxy-FA 18:1 (9Z) and epoxy-FA 18:1 (9E)), though could be separated in RPLC, their MS/MS spectra were indistinguishable (**Fig. S2**). Therefore, C=C positions are annotated in Δ-nomenclature without referring the C=C geometry throughout this study.

**Figure 2.**
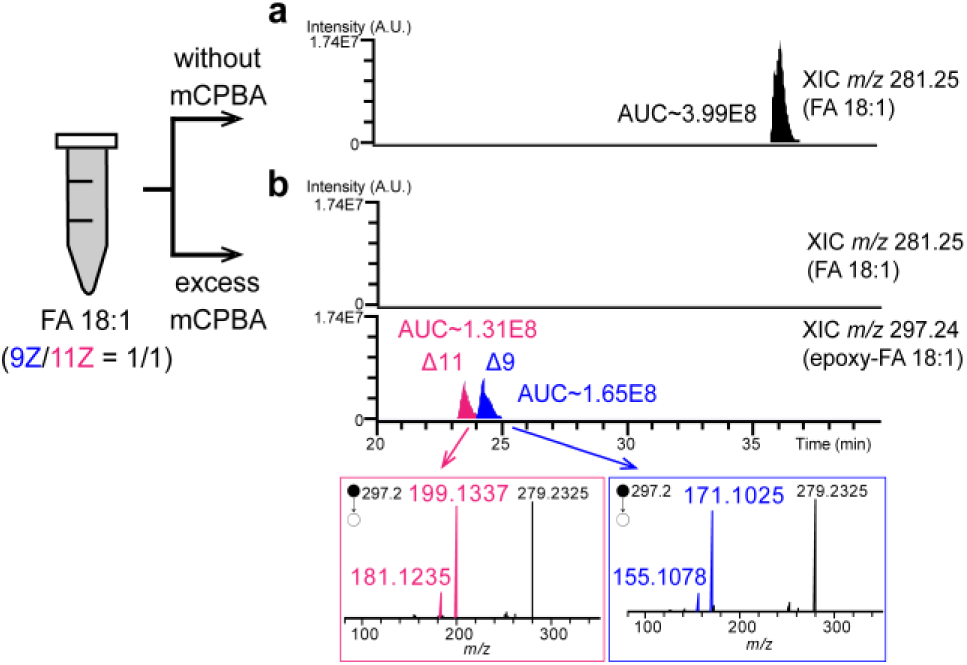
MELDI-LC-MS/MS analysis of a mixture of FA 18:1 C=C isomers. The FA 18:1 C=C isomer mixture (9Z/11Z = 1/1, 100 µM for each) was analyzed by LC-MS/MS **(a)** prior to mCPBA epoxidation and **(b)** after derivatization with excess mCPBA (200 mM) in organic solution, and incubated at 50 °C for an hour to reach to the optimal reaction yield before analysis. The extracted ion chromatograms (XIC) of FA 18:1 and epoxy-FA 18:1, and the area under curve (AUC) were shown.

For polyunsaturated fatty acid (PUFA), we found that when two or more epoxides were tagged, the MS/MS spectrum became complicated, making it difficult to probe the C=C diagnostic ions (**Fig S3**). To solve this issue, we thus targeted MS/MS analysis to the singly tagged products and reconstruct multiple C=C positions of a PUFA. Herein a series of standard PUFAs were thoroughly investigated by MELDI-LC-MS/MS. First, taking FA 18:2 (9Z, 12Z) for example, when moderate epoxidation was conducted, two mono-epoxidized products were found (**Fig. 3a**), and subsequent MS/MS provided structural details by revealing the Δ9 and Δ12 double bond nature. Applying a similar approach to ω-6 FA 18:3 (6Z, 9Z, 12Z), three mono-epoxidized products of C=C bonds at Δ6, Δ9, or Δ12, were revealed (**Fig. 3b**). Its ω-3 isomer, FA 18:3 (9Z, 12Z, 15Z), was also studied, giving a LC-MS/MS spectral set of three epoxy-FA 18:3 products (**Fig. 3c**), which was obviously distinguishable from that of the ω-6 isomer. More cases of PUFA were provided in **Fig S4**. It is noteworthy that, in comparison to MOFAs, the CID fragmentation of mono-epoxidized products of PUFAs was more sophisticated. In specific, the CID fragmentation favors the daughter ions of the cleavages at the sides adjacent to the other unsaturated sites in the epoxides. Such tendency may be due to the generation of the products with conjugated π-bond systems upon CID (**Fig. S5)**. In addition, we also noted that in some cases the fragmentation occurs at the adjacent carbon-carbon bond especially when the epoxide positions were within eight carbon atoms to the carboxyl end. Finally, for a poly-unsaturated GPL, C=C positions in its PUFA chain could be similarly resolved by LC-MS^3^ of its mono-epoxidized products, where CID/CID was applied to the epoxy-PUFA daughter ion in the MS/MS spectrum (**Fig. S6**).

**Figure 3.**
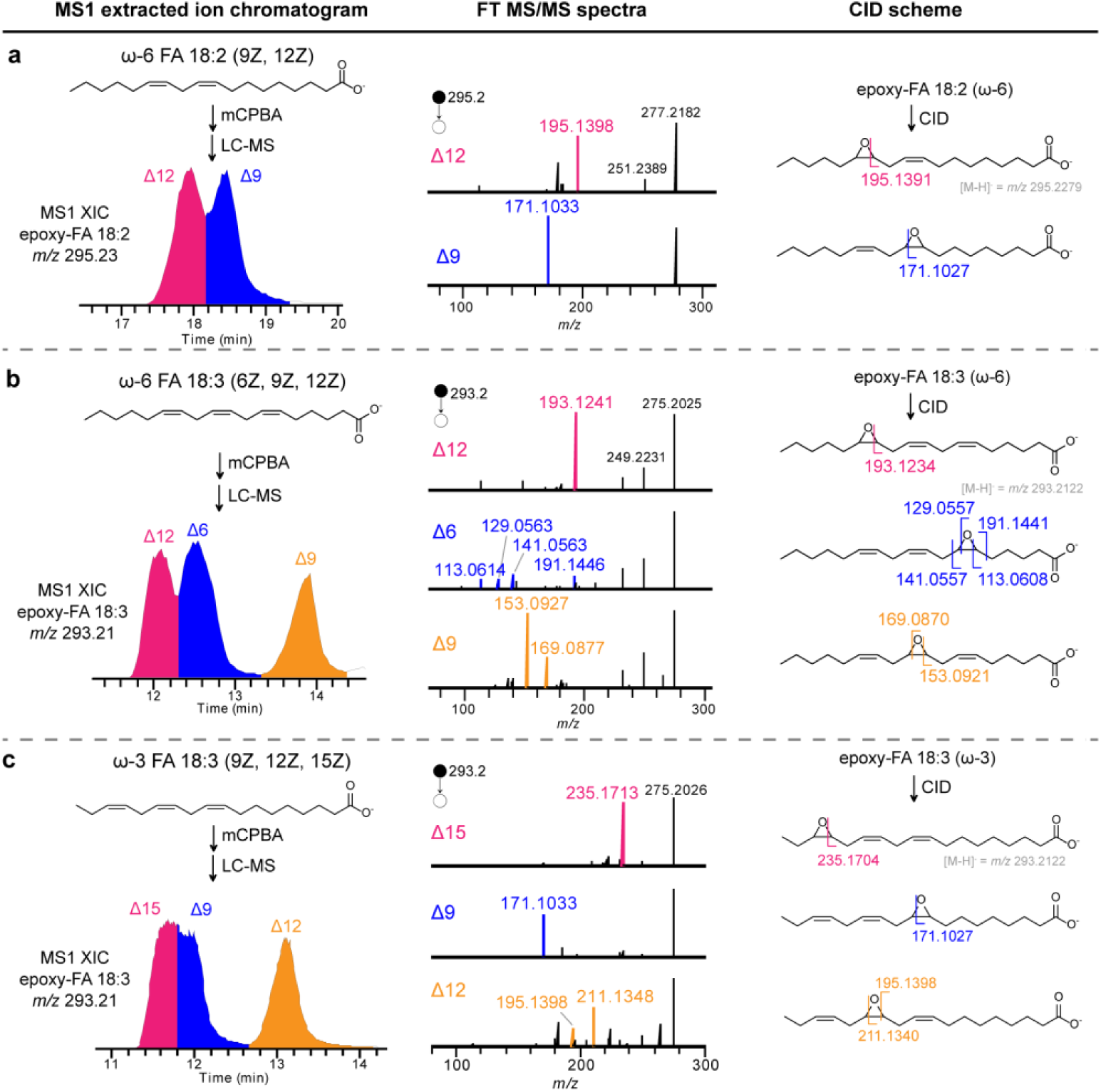
Identification of multiple C=C positions of polyunsaturated lipids by MELDI-LC-MS/MS. The MELDI-LC-MS/MS analysis of polyunsaturated lipids, including **(a)** ω-6 FA 18:2 (9Z, 12Z), **(b)** ω-6 FA 18:3 (6Z, 9Z, 12Z), and **(c)** ω-3 FA 18:3 (9Z, 12Z, 15Z), are demonstrated. Multiple C=C positions of each polyunsaturated lipid are subsequently identified by the MS/MS spectral set of its mono-epoxides.

Importantly, the epoxidation using mCPBA was highly specific to C=C bonds,^35–36^ resulting in appreciable epoxide yields for unsaturated lipids (**Fig. S7, Table S1**). This further made MELDI-LC-MS reliable in probing C=C diagnostic ions from low amount of unsaturated FAs in the nM level, and the limit of detection of C=C diagnostic ions for GPLs was assessed at sub-µM level due to the requirement of MS^3^ (**Fig. S8-S9**).

### Quantification of C=C isomer mixtures with MELDI-LC-MS

Quantification of C=C isomers via MELDI-LC-MS-MS/MS analysis was validated as described in the followings, and experimental details were provided in **Supporting Information**. We first tested a series of mixtures containing two FA 18:1 C=C isomers, 9Z and 11Z, with different molar fractions. The samples were epoxidized with excess mCPBA and analyzed by targeted LC-MS/MS of the epoxide. Then, the peak area of two C=C diagnostic ions for each isomer at different molar fractions was acquired so as to construct the calibration curve (**Fig. 4a**). As a result, a good linearity between the peak area ratios of diagnostic ions (A_11Z_/(A_9Z_+A_11Z_)) and their molar fractions were obtained (R^2^ ∼ 0.999). Interestingly, the slope of the calibration curve (0.98) is close to unity, suggesting that the monounsaturated C=C isomers possess the same levels of epoxidation yields, ionization efficiency, and similar fragmentation features. Such characters are advantageous in determining the relative proportion of each C=C positional isomer in a given unsaturated lipid species, especially when the commercial standards are not available. For absolute quantification of FA 18:1 isomers, D_17_-FA 18:1 (9Z) was spiked into the samples and used as the internal standard (IS) to quantify FA 18:1 (Δ9) and FA 18:1 (Δ11) (**Fig. 4b**). Quantification of PUFA C=C isomers was also validated using a series of ω-3 and ω-6 FA 18:3 isomer mixtures in a similar way, while moderate epoxidation condition was applied to yield high amount of the mono-epoxidized (**Fig S10, Table S2**). **Fig. 4c** showed the obtained calibration curve, which also had a good linearity (R^2^ ∼ 0.999). This enabled the determination of the endogenous ω-3/ω-6 isomeric ratio of FA 18:3 in human sera (94.3 % / 5.7 %, SD = 3.0 %; *n* = 7). Finally, quantifications of ω-6 and ω-9 FA 20:3 mixtures were also validated (**Fig. 4d**).

**Figure 4.**
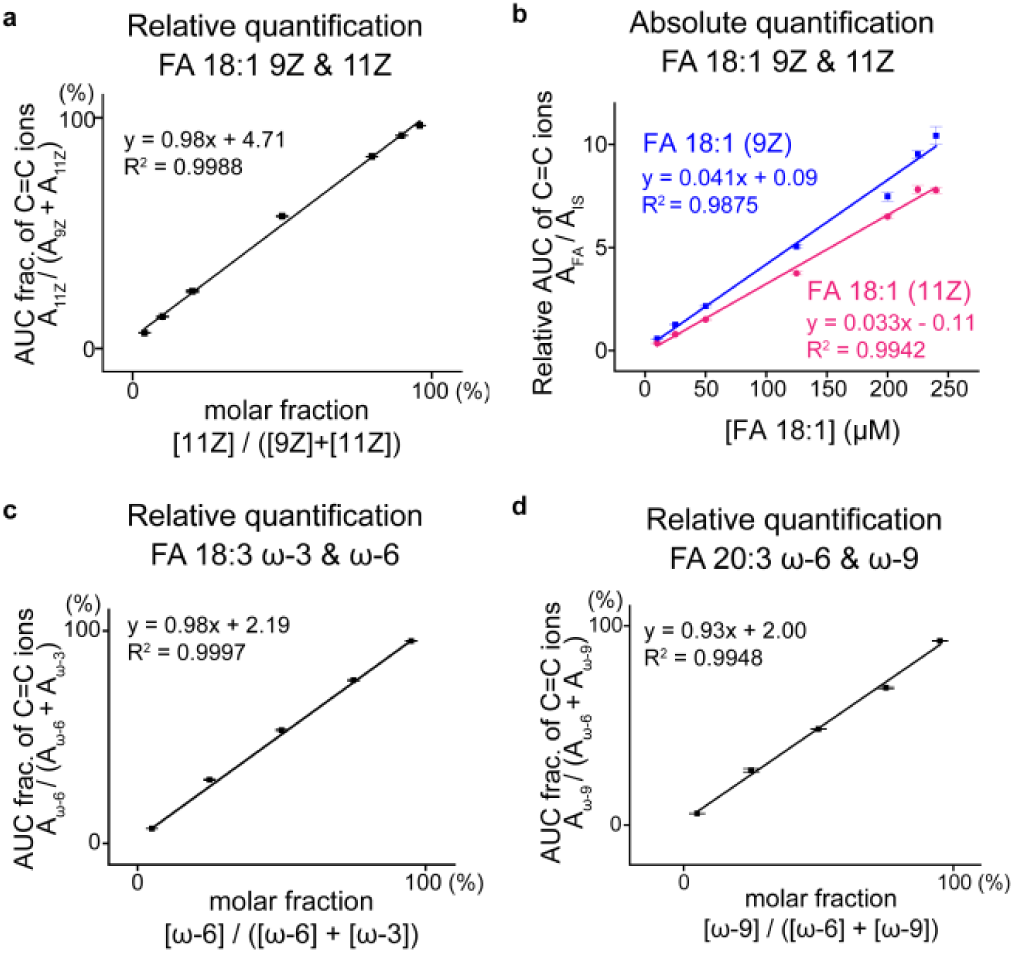
Quantification of lipid C=C isomers by MELDI-LC-MS/MS. The calibration curves for relative quantifications of **(a)** FA 18:1 (9Z) and (11Z) isomers **(c)** FA 18:3 ω-3 and ω-6 isomers **(d)** FA 20:3 ω-6 and ω-9 isomers. **(b)** The calibration curve for absolute quantification of FA 18:1 C=C isomers, where D_17_-FA 18:1 (9Z) was used as internal standard (IS). Each point represents mean ± SD calculated from a triplicate. The experimental details were described in **Supporting Information**.

### Coupling MELDI with LC-MS lipidomics for large-scale analysis of C=C isomers

In order to implement MELDI into the current pipeline of LC-MS-based lipidomics, we developed a two-stage workflow by integrating the conventional LC-MS lipidomics with parallel MELDI-LC-MS analysis for large-scale analysis of C=C isomers in biological lipid extracts (**Fig. 5**). In the first stage, as shown in **Fig. 5a**, traditional LC-MS/MS analysis of a biological lipid extract is executed, and the resulting data was processed by LipidSearch (Thermo) to give a list of identified lipid species, including putative structural information of fatty acyl chain composition (e.g. length and number of C=C bonds of each fatty acyl chain). The report is then transformed by EpoxyFinder, a lab-built graphic-user-interface (see **Supporting Information**), into a list of predicted mono-epoxide ions that would be used in subsequent MELDI-LC-MS analysis.

**Figure 5.**
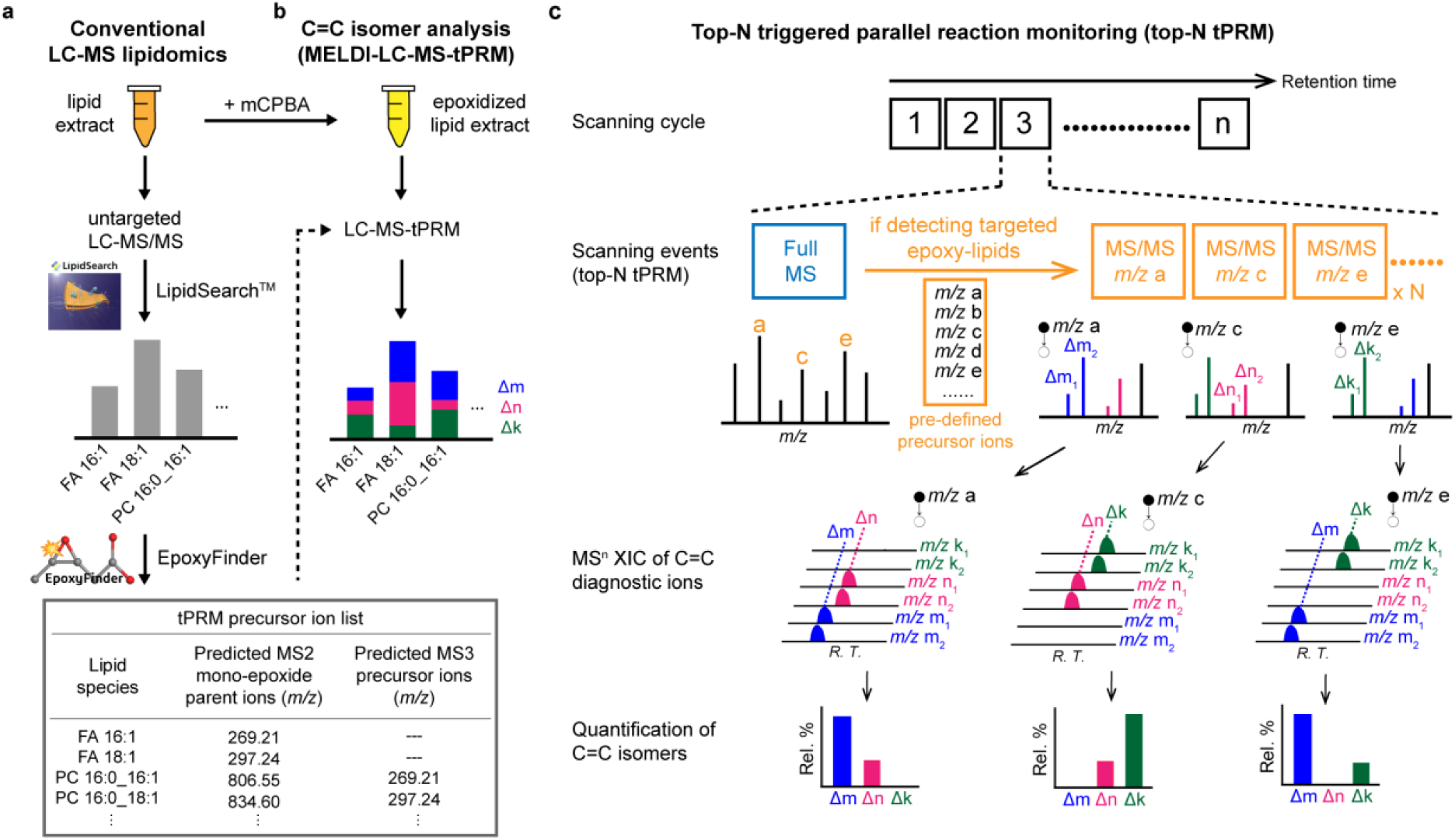
The workflow for LC-MS-based analysis of C=C isomers in biological lipid extract using MELDI. **(a)** In the first stage of the workflow, conventional untargeted LC-MS/MS analysis is applied, and the data is subsequently processed by LipidSearch™, giving structural information of unsaturated lipids in the level of fatty acyl chain compositions. Then, EpoxyFinder, a lab-built graphical-user-interface, is used for data processing to create a precursor ion list of mono-epoxidized unsaturated lipids for subsequent targeted analysis of C=C isomers. **(b)** In the second stage, the lipid extract is derivatized with mCPBA and then examined with LC-MS-triggered parallel reaction monitoring (tPRM) analysis for identification and quantification of C=C isomers. **(c)** Schematic representation of tPRM. In a top-N tPRM module, each scanning cycle contains one full-FTMS scan, followed by multiple IT-MS/MS (or MS^3^) acquisitions of top-N most intense epoxy-lipid ions (if they are present) in the full-FTMS spectrum throughout the whole analysis. Each lipid C=C isomer is subsequently quantified by summed extracted ion chromatogram (XIC) area of a set of C=C diagnostic ions under the corresponding MS/MS scanning channel of the epoxy-lipid ion.

In the second stage (**Fig. 5b**), MELDI-LC-MS analysis is applied to examine the targeted unsaturated lipids in the epoxidized sample. In this study, two individual derivatization protocols were used for analyses of monounsaturated and polyunsaturated lipids respectively, as only mono-epoxidized products were targeted for MS/MS (see **Fig. S11-S12)**. To enhance instrumental performance, we integrate MELDI with triggered parallel reaction monitoring (tPRM)^37^ as an innovative method for semi-targeted LC-MS analysis of C=C isomers. The method was demonstrated using an LTQ-Orbitrap mass spectrometer. In fact, tPRM can be regarded as a modified data-dependent acquisition (DDA), in which the targeted precursor list is pre-defined and the precursor ion exclusion is turned off. The concept of LC-MS-tPRM analysis is illustrated in **Fig. 5c**. In short, as it continuously acquires full FT-MS spectra when analytes are eluting, and once the pre-defined targeted precursor ions (*i.e.* mono-epoxidized lipids) are detected, the corresponding MS/MS events will be triggered in each scanning cycle (*e.g.* the most N intense precursor ions are selected for MS/MS acquisition in a top-N tPRM method). In comparison to tradition multiple reaction monitoring (MRM), which measures a limited number of targeted daughter ions at narrow spectral windows, tPRM simultaneously measures all daughter ions generated in a full MS/MS spectrum. More importantly, such feature of tPRM is advantageous to acquire multiple C=C diagnostic ions from each epoxy-lipid with minimal matrix effect, allowing accurate identification and quantification of C=C isomeric mixtures. To ensure the optimal performance of the analysis, a top-25 tPRM method (maximum cycling time ∼ 0.1 minute) was applied throughout the following studies, whereas unsaturated lipids of high abundance were specifically targeted by EpoxyFinder. Details of the data processing procedures were elaborated in **Supporting Information (SI section 3.3)**.

As a pilot study, the workflow was first applied to the lipid extract of the human serum. As a result, 80 of the abundant unsaturated lipid species were targeted for isomer analysis, including 44 mono-unsaturated and 36 polyunsaturated lipids species (**Table S3-S4**), from which we identified 58 monounsaturated C=C isomers with C=C positions varying from Δ6 to Δ15 (**Table S5**), and 48 polyunsaturated C=C isomers (**Table S6**). In short, we concluded that more than 100 C=C positional isomers from 59 targeted unsaturated lipid species in human serum were successfully identified (**Fig. 6a**). Representative cases of the identified C=C isomers were shown in **Fig. 6b-c**, indicating that C=C isomers were separated after derivatization and robustly identified and quantified by their diagnostic ion pairs. Similar interrogations of more identified isomers are elaborated in **Fig. S13-S21**.

**Figure 6.**
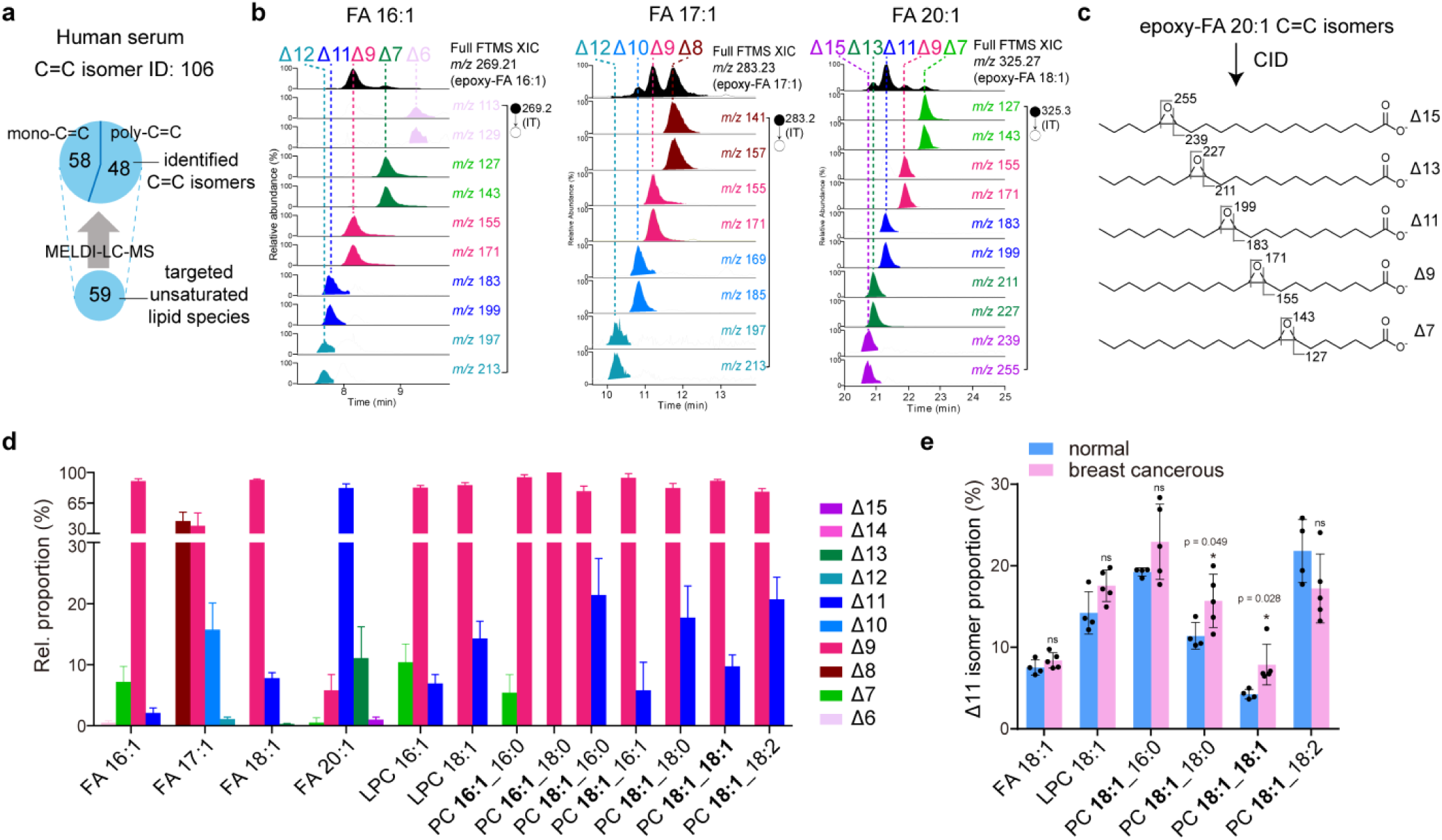
Lipid C=C isomers in human serum revealed by MELDI-LC-MS isomeric lipidomics. **(a)** The total number of the identified C=C isomers from the 59 targeted unsaturated lipid species in human serum. **(b)** LC-MS/MS identifications of novel C=C isomers from FA 16:1, FA 17:1, and FA 20:1 in human serum. The XICs of epoxidized parent ions (in full MS) and C=C diagnostic ions (in MS/MS) were shown. **(c)** The fragmentation schemes of the identified FA 20:1 C=C isomers. **(d)** Quantification of relative proportions of monounsaturated C=C isomers in the lipid extract of human serum (*n* = 10). The targeted monounsaturated FA chains in GPLs being quantified are shown in bold. (**e)** Comparison of relative proportions of Δ11 isomers in FA 18:1-containing lipids between healthy human sera (*n* = 4) and breast cancerous human sera (*n* = 5). Error bars represent SD. Statistical analyses (*t* test) are shown as: * P < 0.05; ** P < 0.01; *** P < 0.005.

For quantitative analysis, we determined the C=C isomer compositions in 18 unsaturated lipid species in 10 healthy (non-cancerous) human sera, as showed in **Fig. 6d**. Remarkably, to the best of our knowledge, the presence of many uncommon FA C=C isomers (*e.g.* FA 16:1 Δ6/ Δ7/Δ11; FA 18:1 Δ13; FA 20:1 Δ7/Δ9/Δ15) that are listed in LIPID MAPS database were unraveled in human serum for the first time, and their FT-MS/MS spectra were validated (**Fig. S13-S18**). These C=C isomers were of low endogenous quantities and rarely found in human serum via other C=C isomer resolving methods,^31,33^ which might be due to the matrix effect. However, using our platform, these low-abundant C=C isomers were separated by RPLC after derivatization, and thus their relative proportions could be estimated by the peak area of diagnostic ions without severe ion interferences. Moreover, we showed that the ω-9 isomers were the most dominant species in FA 18:1, FA 20:1, FA 22:1, FA 24:1, and all the FA 18:1 chains in GPLs, which was the reflection that ω-9 FA 18:1 serves as the original species for the downstream biosynthesis of these ω-9 monounsaturated lipids in lipogenesis mechanism in mammalian cells.^17^ On the other hand, we revealed that ω-6 family isomers dominated many of the PUFAs, including FA 18:2 and its biosynthetic products, FA 20:2, FA 20:3, FA 20:4, FA 22:4, and their constituting GPLs (**Table S6**). These quantitative observations were consistent with the previous study on PUFAs,^14^ and here we further enlarged the body of knowledge by showing that the ω-6 isomers also dominated these PUFA-constituting GPLs in human serum.

Finally, we found that although the absolute concentrations of FA 18:1 were diverse among the human sera (141.4 ± 56.7 μM for Δ9; 18.0 ± 5.4 μM for Δ11; *n* = 10), the compositions of C=C isomers in FA 18:1 and its constituting GPLs were quite similar (e.g. 91.8 ± 0.9 % for FA 18:1 (Δ9); 7.8 ± 0.9 % for FA 18:1 (Δ11); **Table S5**), indicating that C=C isomer compositions were held consistent among healthy individuals. In this regard, we hypothesized that such isomeric lipidome may serve as potential indicators to lipid homeostasis, which can be disturbed due to disease. To further prove it, we compared the healthy and breast cancerous human sera with a specific focus on isomers compositions in FA 18:1-related species. As a result, we found that in the breast cancerous human sera, the proportion of the Δ11 isomer in both PC **18:1**_18:2 and PC **18:1**_18:0, but not FA 18:1 were statistically increased (\**P* < 0.05) (**Fig. 6e and Table S7**). In conclusion, our method will allow a large-scale cohort study to evaluate if such C=C isomer ratio changes can be used as breast cancer biomarkers.

To evaluate the efficacy in studying samples of other biological systems, we applied the method to investigate isomeric lipidome in murine 3T3-L1 adipocytes, one of the most common *in vitro* models to study lipid biology. First, we concluded that the differentiated 3T3-L1 adipocytes possessed a considerable diversity of unsaturated lipid isomers, rendering more than 170 C=C isomers identified from 66 unsaturated lipid species **(Fig. 7b)**. In specific, from the targeted unsaturated lipid species (**Table S8-S9**), 77 isomers were identified and quantified in 20 monounsaturated lipid species (**Table S10**), whereas up to 96 isomers were identified in 46 polyunsaturated lipid species (**Table S11**). More LC-MS/MS annotations of the identified isomers were provided in Fig. **S22-S31**. Similarly, we also conducted quantitative analysis in isomer compositions in the targeted monounsaturated lipid species (**Fig. 7a**). Again, many novel isomers of low abundance were revealed, further supported by FT-MS/MS annotations (**Fig. S26-S32**).

**Figure 7.**
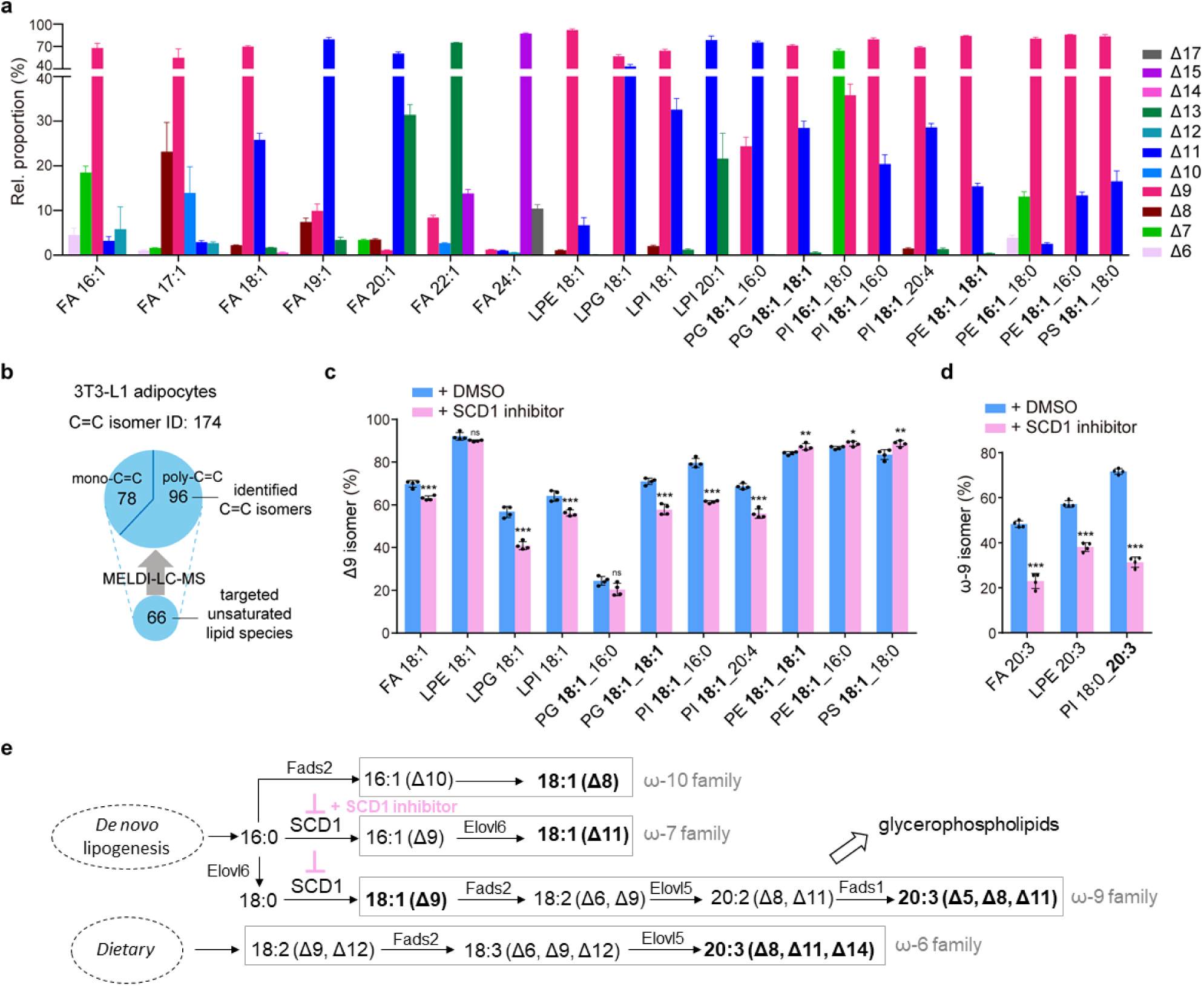
Lipid C=C isomers in the 3T3-L1 adipocytes revealed by MELDI-LC-MS isomeric lipidomics. **(a)** Quantification of mono-unsaturated C=C isomers in the lipid extract of differentiated 3T3-L1 adipocytes (*n* = 4). The targeted monounsaturated FA chains in GPLs being quantified are shown in bold. **(b)** The total number of the identified C=C isomers from 66 targeted unsaturated lipid species. **(c-d)** Alteration of C=C isomers between normal (*n* = 4) and SCD1 inhibitor-treated 3T3-L1 adipocytes (*n* = 4) is accessed in **(c)** Δ9 isomers in FA 18:1-containing lipids and **(d)** ω-9 isomers in FA 20:3-containing lipids. Error bars represent SD. Statistical analyses (*t* test) are shown as: * P < 0.05; ** P < 0.01; *** P < 0.005. **(e)** Cellular lipogenesis pathways, where the SCD1 inhibition was indicated and the targeted isomers in **(c)** and **(d)** were shown in bold. The scheme was modified from Guillou et al.^17^

As unsaturated lipid isomers are individually regulated through enzymatic pathways in cellular lipogenesis,^17^ their alterations are thus potential indicators to how cells reprogram lipogenesis in response to external perturbations. For example, aberrant SCD1 activity is associated with disrupted cellular lipidome, even in coordination with progressions of diabetes, fatty liver diseases, and cancers.^38^ A comprehensive analysis of cellular lipidome may thus shed light on such biochemistry in the molecular level. With this regard, we constructed a 3T3-L1 adipocyte model with treatment of stearoyl-coenzyme A desaturase 1 (SCD1) inhibitor,^39^ and compared its isomeric lipidome with that of the normally differentiated adipocytes. As a result, we found that in the SCD1 inhibitor-treated model, (i) relative proportions of Δ9 isomer in FA 18:1 and many of the FA 18:1-containing lipids were significantly decreased (**Fig. 7c and Table S12**), and (ii) relative proportions of ω-9 isomers in FA 20:3 and its constituting lipids were significantly decreased (**Fig. 7d and Table S13**). Importantly, our results were indicative to the inhibited function of SCD1, which is, in fact, the key enzyme that catalyzes the formation FA 18:1 (Δ9), the starting material for downstream synthesis of ω-9 unsaturated lipids (**Fig. 7e**).^17,39^ Our results revealed how the alleviated SCD1 activity mediates cellular lipogenesis at the level of isomeric lipidome.

### Mapping alterations of lipid C=C isomers on tissue sections by MELDI-DESI

MSI techniques allow visualization of compounds in biological tissue sections, providing chemical-rich information associated with tissue heterogeneity. MSI of lipids has been largely used in tissue-typing and cancer diagnosis.^40^ However, such lipid imaging can provide only the summed distribution of multiple isomeric species.^11^ Here, we reported a C=C positional isomer-resolving imaging method by coupling MELDI with desorption electrospray ionization (DESI)^41^ MSI. For ease of discussion, we will refer to this integrated strategy as MELDI-DESI-MSI.

The instrumental capability in quantifying C=C isomers was evaluated (**Fig. S32-S33)**, using the intensity ratios of diagnostic ions to indicate the molar ratios of the C=C isomers. For C=C isomer imaging, high-resolution MSI (at MS1 level) was first performed in a tissue section without MELDI to obtain the lipid profile and spatial distribution, whereas its serial section was used for C=C isomer imaging. To this end, we used a simple airbrushing method to *in situ* epoxidize unsaturated lipids with mCPBA in the tissue section (**Fig S34**), which was modified from the matrix deposition method used in MALDI MSI,^42^ and a demonstration video was also provided (see **Supporting Information, section 6-3**). Notably, the entire derivatization process took less than 10 minutes and was implemented in ambient condition, and the spatial resolving power after mCPBA pretreatment was retained as validated in **Fig. S35**. After epoxidation pretreatment, C=C isomers of an unsaturated lipid species were mapped with targeted MS/MS (or MS^3^) imaging.

Using the mouse kidney sections for our first example of MELDI-DESI-MSI, as shown in **Fig. 7b**, the ion image of *m/z* 281.25, which represented the summed distributions of all FA 18:1 C=C isomers, was observed with a slightly lower intensity in the kidney medulla. In the adjacent section pretreated by mCPBA, the epoxidation product, epoxy-FA 18:1 (*m/z* 297.24), could be clearly observed in the full-MS spectrum (**Fig. S36b**). The on-tissue CID spectrum of epoxy-FA 18:1 clearly confirmed the presence of two C=C isomers with C=C positions at either Δ9 or Δ11 (**Fig. S36c**). Targeted MS/MS imaging of epoxy-FA 18:1 enabled the semi-quantitative molecular imaging of C=C isomers by fractional distribution image (FDI) of Δ9 and Δ11 isomers (**Fig. 7c**). Here, an enriched level of the Δ11 isomer, and conversely a decreased level of Δ9 isomer, was observed in the kidney medulla (**Fig. 7d**), which implied that the decreased level of total FA 18:1 in the kidney medulla might be the result of decreased Δ9 isomer. This highlighted that FDI images showed the changes in the ratio of diagnostic ion intensities rather than the actual local concentration of C=C isomers, while alteration of C=C isomer composition could be clearly reflected by the FDI images across the tissue section. The same approach was performed for PG 16:0_18:1 in the other set of serial kidney sections. Full-MS imaging of *m/z* 747.52 showed a relatively lower PG 16:0_18:1 abundance in the kidney medulla (**Fig. 7e**). The *in situ* MS^3^ spectrum of epoxy-PG 18:1_16:0 (*m/z* 763.51) in the epoxidized section also showed a mixture of Δ9 and Δ11 isomers (**Fig. S36d**). Mapping the C=C isomers through FDI revealed the enrichment of the Δ11 isomer proportion in the kidney medulla (**Fig. 7f)**. Interestingly, the overall fraction of Δ11 isomer in PG 18:1_16:0, calculated to be about 90%, was remarkably higher than that of FA 18:1 (**Fig. 7g**), implying that FA 18:1 (Δ11) isomer was preferably selected during synthesis of PG 18:1_16:0 in the mouse kidney. In yet another perspective, we noted that the observed isomer compositions in PG might be contributed by its *sn*-substitutional isomer, bis(monoacylglycero)phosphate (BMG). Therefore, we believe that the combination of MELDI with other sn-substitutional isomer-resolving techniques will allow a more precise structural elucidation of lipids in isomeric imaging studies.

It has been shown that alteration of the ratios of C=C isomers can be used as makers for cancerous tissues,^33,43–44^ an MSI method that can easily visualize these isomers may thus be used to determine tumor margins. Such proposed idea was here demonstrated using metastatic lung tissues invaded by breast tumor cells (**Fig. 7h**). As shown in **Fig. 7i**, is was unable to distinguish the tumor region using the summed distribution of PG 18:1_16:0, later resolved as the mixture of two C=C isomers of Δ9 and Δ11 on the mCPBA-pretreated section (**Fig. S37d**). Intriguingly, with MELDI, we revealed that the fraction of PG 16:0_18:1 Δ11 isomer was significantly elevated in the tumor cell regions (**Fig. 7j and 7k**), suggesting that a single pair of C=C positional isomers could be utilized to determine the tumor margin.

**Figure 8.**
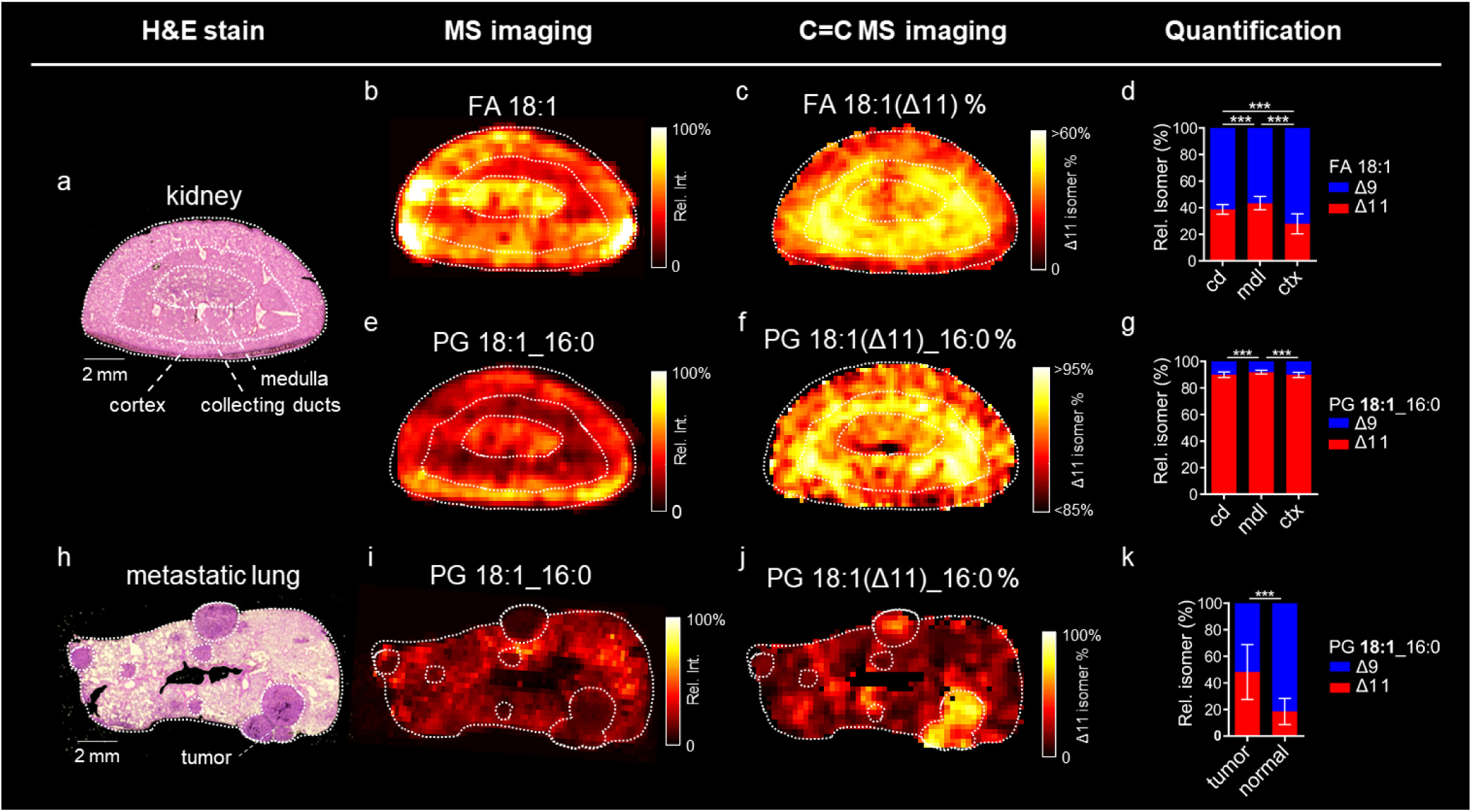
Mapping C=C isomers in mouse tissues using MELDI-DESI-MSI. **(a-f)** C=C isomer imaging of a mouse kidney section. Distinct tissue regions are outlined by white dash lines, including cortex (ctx), medulla (mdl), and collecting ducts (cd) in the H&E staining image in **(a)**. Full-MS images of **(b)** FA 18:1 (*m/z* 281.25) and **(e)** PG 18:1_16:0 (*m/z* 747.52) are shown, and their corresponding C=C images of the Δ11 isomer are shown as fractional distribution image (FDI) of Δ11/(Δ9+Δ11) % in **(c)** and **(f)**, respectively. **(h-j)** C=C isomer imaging of a metastatic lung tissue. **(i)** Full-MS image of PG 18:1_16:0 and **(j)** the corresponding FDI of the Δ11 isomer. (**d, g, k)** Quantification of relative proportions of C=C isomers in each tissue region (*t* test, *** P < 0.005). Errors bars represent SD.

## Conclusion

We concluded that MELDI is a universal, user-friendly, and cost-effective derivatization strategy to pinpoint lipid C=C positions via conventional MS/MS. To be specific, epoxidation with mCPBA gives several outstanding features, including (i) one-pot, rapid, catalyst-free, single-step derivatization in organic solvents under mild conditions, (ii) high specificity to C=C bonds and appreciable yield of epoxide (**Table S1)**, (iii) inexpensive reagents (market price ∼0.1 USD per gram of mCPBA powder, and (iv) high stability of epoxides in room temperature (**Fig. S11c**) (v) feasibility for a broad range of lipid classes (**Fig. S37 and Table S14)**, suggesting an innovative way for studying lipid C=C isomers in biomedical research.

An inevitable demand in lipidomics is to elucidate chemical structures of potential lipid biomarkers, which is challenging due to the presence of structural isomers. With the advanced MS-based techniques in the past years,^22,33,43^ structural assignments of unsaturated lipid isomers becomes increasingly possible, leading to a growing understanding of unsaturated lipid isomerism and its association with disease progression. In addition to these specialized techniques, MELDI brings us to a comprehensive analysis of biological lipids. In this regard, we have achieved (i) large-scale identification of unsaturated lipid isomers in human serum, where a variety of uncommon low-abundant isomers was quantified for the first time, (ii) bridging compositional alterations in isomeric lipidome with desaturase dysfunction of the differentiated murine adipocytes, and (iii) imaging of the tumor-associated changes in unsaturated lipid isomers on a metastatic tissue section. Importantly, these findings were hardly provided through conventional lipidome analysis, but here unraveled using MELDI and commercial mass spectrometers without any instrumental modification. Finally, to make future studies easier to follow the identification process, we provided a comprehensive summary of the diagnostic ions that have been annotated in this study (**Table S15**). We believe that MELDI can be readily implemented to global MS core facilities, which may thereby speed up mining of underlying lipid biochemistry by providing a deep level of structural information in routine lipidomics experiments.

In the end, we would like to highlight that MELDI is of great potential as being integrated into versatile MS platforms for analyses of unsaturated lipid isomers. Besides (LC-)ESI and DESI, MELDI is compatible to many frequently used ionization sources, including MALDI, paper-spray ionization, and atmospheric pressure chemical ionization (APCI) (**Fig. S38-S40**), allowing many non-instrument-oriented lipid scientists to study lipid C=C isomers in a routine manner.

## ASSOCIATED CONTENT

## SUPPORTING INFORMATION

Materials and methods, the spectral data set of lipid C=C isomer standards, features of mCPBA epoxidation, data processing strategy in LC-MS-tPRM analysis, EpoxyFinder software package, the demonstration video as well as the method for for *in situ* derivatization using mCPBA, and other related contents are provided in Supporting Information.

## AUTHOR INFORMATION

### Author Contributions

T.-H. K. and C.-C. H. designed the experiments. T.-H. K. and H.-H. C. carried out all the MS experiments and analyzed the data. H.-Y. C. constructed the cell experiments. C.-W. L., T.-L. S., and M.-Y. W. designed and carried out the animal experiments and collected the clinical samples. C.-C. H. supervised the study. T.-H. K. and C.-C. H. wrote the paper.

### Notes

The authors declare no competing financial interest.

## ACKNOWLEDGMENT

This research was supported by Ministry of Science and Technology (MOST), R.O.C. (Grant nos.: MOST 106-2113-M-002-013-MY2, 107-2321-B-001-038-, and 108-2636-M-002-008-), and Center for Emerging Materials and Advanced Devices, National Taiwan University (NTU) (Grant nos.: NTU-ERP-108L880116). T.-H. K. acknowledges the financial support from National Taiwan University, and Institute of Nuclear Energy Research, Atomic Energy Council, Executive Yuan, R.O.C. The instrument support from NTU Mass Spectrometry Platform was acknowledged. The technical assistance from Ying-Chen Huang, Li-En Lin, Kai-Hung Huang, and Dr. Qiang Lyu in C.-C. H. Lab, Ko-Chien Chen and Yu-Ju Wang in T.-L. S. Lab, and Bo-Rong Chen in M.-Y. W. Lab, was also acknowledged.

---

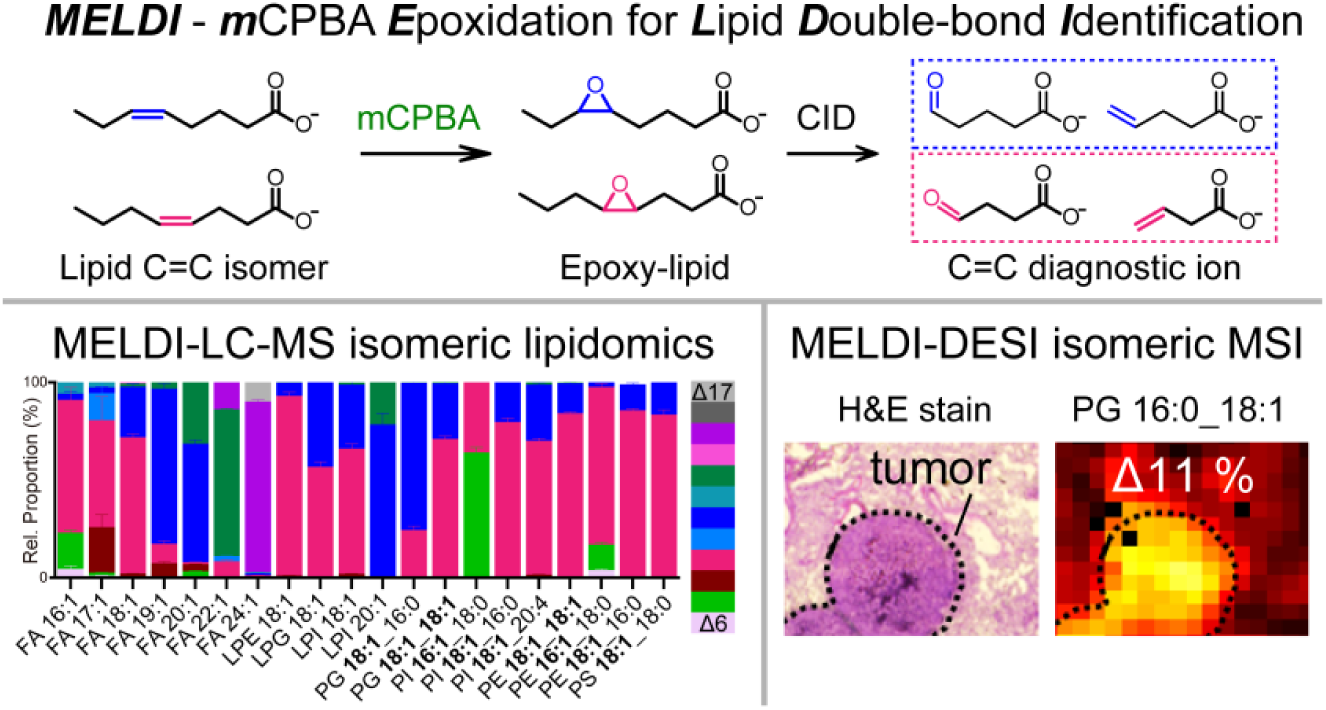

