## Supplementary Information for "Deep Lipidomics and Molecular Imaging of Unsaturated Lipid Isomers: A Universal Strategy Initiated by mCPBA Epoxidation"

for

Contents

S0. Materials and Methods

S1. Identification of C=C positions in unsaturated lipids using MELDI-MS/MS

S1-1. Monounsaturated GPLs

S1-2. C=C geometric isomers

S1-3. Polyunsaturated FAs and GPLs

S1-3-1. FT-MS/MS spectra of fully epoxidized PUFAs

S1-3-2. Identification of C=C positions in mono-epoxidized PUFAs by LC-MS/MS

S1-3-3. The CID scheme of mono-epoxidized PUFAs

S1-3-4. Identification of C=C positions in mono-epoxidized GPLs by LC-MS/MS/MS

S2. Features of mCPBA epoxidation

S2-1. Epoxide yield of unsaturated FAs and GPLs

S2-2. Limit of detection of C=C diagnostic ions

S2-3. Controlling epoxidation for studying PUFA C=C isomers

S3. MELDI-LC-MS-tPRM platform for large-scale investigation of C=C isomers

S3-1. Development of the mCPBA epoxidation protocols for biological lipid extracts

S3-1-1. Monounsaturated lipids

S3-1-2. Polyunsaturated lipids

S3-2. EpoxyFinder: creating the targeted epoxide list for LC-MS-tPRM analysis

　	　 S3-2-1. Supplementary software: EpoxyFinder

S3-3. The strategy in processing LC-MS-tPRM data

S3-3-1. MOFA

S3-3-2. PUFA

S4. MELDI-LC-MS-tPRM analysis in human serum lipid extracts

S4-1. The LC-MS-tPRM target ion list for human serum lipid extract

S4-2. Identification and quantification of C=C isomers in human serum

S4-2-1. Monounsaturated lipids

S4-2-2. Polyunsaturated lipids

S4-3. Data processing: identification of C=C isomers

S4-4. Data processing: quantification of C=C isomers

S4-5. Representative LC-MS-tPRM results of the identified C=C isomers in the human serum

S4-6. Healthy vs breast cancerous human serum

S5. MELDI-LC-MS-tPRM analysis in 3T3-L1 adipocyte lipid extracts

S5-1. The LC-MS-tPRM target ion list for 3T3-L1 adipocyte lipid extracts

S5-2. Identification and quantification of C=C isomers in 3T3-L1 adipocytes

S5-2-1. Monounsaturated lipids

S5-2-2. Polyunsaturated lipids

S5-3. Representative LC-MS-tPRM results of the identified C=C isomers in the 3T3-L1 adipocyte

S5-4. Normal vs SCD1 inhibitor-treated

S6. MELDI-DESI-MS imaging platform for mapping C=C isomers in tissue sections

S6-1. Quantification of C=C isomer standards using MELDI-DESI-MS/MS

S6-2. Limit of detection of C=C diagnostic ions with MELDI-DESI MS/MS.

S6-3. Development of an airbrushing method for *in situ* epoxidation in biological tissue sections

S6-3-1. Method development

S6-3-2. Demonstration video

S6-3-3. Protocol

S6-3-4. Validation of spatial resolution

S6-4. *In situ* DESI-MS analysis on mouse tissue sections

S6-4-1. Mouse kidney sections

S6-4-2. Mouse metastatic lung sections

S7. Identification of C=C positions of unsaturated lipids in positive ionization mode

S8. Compatibility to commercial mass spectrometers

S9. Summary of the C=C diagnostic ions of the identified mono-epoxidized unsaturated lipids

S10. References

**S0. Materials and Methods**

**Chemicals and reagents**

Glycerophospholipid (GPL) standards were purchased from Avanti Polar Lipids (Alabaster, AL, USA). Fatty acid (FA) standards were purchased from Cayman Chemical (Michigan, USA). Methanol (MeOH), methyl tert-butyl ether (MTBE) and isooctane was purchased from Honeywell (Michigan, USA). *meta*-chloroperoxybenzoic acid (mCPBA) was purchased from Sigma Aldrich (St. Louis, MO) and used without further purification. Ammonium acetate and 2-propanol (IPA) was purchased from Sigma-Aldrich (Missouri, USA). Acetonitrile (ACN) was purchased from J.T. Baker (Phillipsburg, NJ). Hydrochloric acid (HCl) was purchased from Acros Organics (New Jersey, USA) Phosphate buffered saline (PBS) was purchased from Bioman Scientific (Taipei, Taiwan). Dimethylformamide (DMF) was purchased from Tedia (OH, USA). Ultrapure water (18.2 MΩ cm) was prepared by a Milli-Q system (Millipore, Milford, MA).

**Extraction of human serum lipids**

All human sera were provided by National Taipei University Hospital under Institutional Review Board (NTUH-REC No.: 201512122RINC) protocol and approved by the Ethics Committee of National Taipei University Hospital Board. All the serum samples were stored in -80 ^o^C before extraction. To extract lipids from serum, the MTBE extraction protocol ^1^ was applied with laboratory modifications according to Giera et al ^2^. The internal standard (IS) solution, containing five GPLs 14:0/14:0 (PC, PA, PG, PS, and PE; 10 μg/mL for each), D_31_-FA 16:0 (10 μg/mL), and D_17_-FA 18:1(9Z) (250 μg/mL), was prepared beforehand. An aliquot of 100 μL serum were spiked with 20 μL of IS mixtures and subsequently extracted by addition of 600 μL MTBE and 150 μL MeOH in a 1.5-mL micro-centrifuge tube and vortexed for 30 minutes at room temperature. For phase separation, 200 μL dd-water were added and the sample was centrifuged for 3 min at 12,000 rpm. The upper organic portion was collected into another 1.5-mL microcentrifuge tube. The extraction was repeated by adding 100 μL water, 100 μL MeOH and 300 μL MTBE and vortexed for additional 10 minutes, and after centrifugation the upper portion was collected together. The combined solution was dried in a vacuum concentrator (Vacufuge plus Vacuum Concentrator, Eppendorf) for at least 3 hours, giving the yellowish extracts. The extracts were added 100 μL of reconstituted solution (ACN/IPA/water, v/v/v = 65/30/5) and ultrasonicated for 3 minutes. The reconstituted samples were stored in -80 ^o^C before further analysis.

**3T3-L1 adipocytes cultures**

***Chemical, cell line, and reagent.*** Murine 3T3-L1 adipocytes cell line (ATCC number: CL-173) was purchased from Bioresource Collection and Research Center (BCRC; Taiwan). Cell culture reagents including Dulbecco’s modified Eagle’s medium, fetal bovine serum, and 0.5% Trypsin-EDTA were purchased from Gibco by Life Technologies (Carlsbad, CA, USA). Penicillin–streptomycin solution was purchased from Hyclone Laboratories (Logan, UT, USA). Other cell culture reagents including 3-isobutyl-1-methylxanthine (IBMX), dexamethasone, human insulin, and Oil Red O dye were purchased from Sigma Aldrich (St Louis, MO, USA). Dimethyl sulfoxide (DMSO; ≧99.9% purity) were purchased from BioShop Life Science (Burlington, ON, Canada). SCD1 inhibitor, CAY10566, was purchased from Cayman Chemical (Ann Arbor, MI, USA).

***Cell culture experiment.*** The experiment of adipocyte differentiating was followed by the previous protocol (J. C. Ralston et al. ^3^). 3T3-L1 adipocyte cells were incubated in 5.0 % CO_2_ at 37 °C. Adipocytes were cultured in main medium consisted of Dulbecco’s modified Eagle’s medium (DMEM), 100 unit/mL penicillin–streptomycin solution, and 10.0 % heat-inactivated fetal bovine serum. Cells were seeded at a density of 7.0 × 10^5^ cells per 100-mm dish. The SCD1 inhibitor (CAY10566) was dissolved in DMSO to form 10 μM stock solution, and a final working concentration of 10 nM was used in all cell experiments. On day 0, 3T3-L1 adipocyte differentiation was incubated with standardized differentiation reagents consisting of IBMX (0.5 mM), dexamethasone (1 μM) and human insulin (5 μg/ml) in basic cell culture medium. The medium was changed to the medium consisted of human insulin (5 μg/ml) in cell culture medium at day 2. On day 4, the fetal bovine serum from the basic cell culture medium was removed to accomplish serum-free conditions, and adipocytes were treated with either 10 nM SCD-1 inhibitor or an equivalent volume of DMSO (as the control experiment). On day 7, cells were stained with Oil Red O to confirm lipid droplets accumulation in the cytoplasm.

**Extraction of 3T3-L1 adipocytes lipids**

The extraction followed the previous protocol (J. Li *et al.* ^4^) with laboratory modification. Cell pellets were added 1 mL MeOH and 10 uL IS solution, and acidified by HCl to a final concentration of 25 mM. Then 1 mL isooctane was added and the sample was vortexed for 30 minutes. 300 μL PBS was added to form phase separation, and then the upper layer was collected and dried under vacuum for 45 minutes. The extract was dissolved with 50 μL reconstituted solution, sonicated, and stored at -80 ^o^C.

**Quantification of lipid isomers**

***The calibration curve for relative quantification of FA 18:1 9Z and 11Z isomers.*** The total concentration of FA 18:1 was kept at 250 μM with the molar ratio varied ([11Z]/[9Z] = 1/24, 1/9, 1/4, 1/1, 4/1, 9/1 and 24/1). Each mixture was derivatized with excess mCPBA (200 mM) at 50 ^o^C for 1 hour and then analyzed by LC-MS-tPRM. The calibration curve was constructed by plotting the molar fractions of the 11Z isomer ([11Z]/([9Z]+[11Z])%) against the fractions of the summed extracted ion chromatographic (XIC) area of diagnostic ions (A_9Z_/(A_11Z_+A_9Z_) %), where A_9Z_ = [A_m/z 155_+A_m/z 171_] and A_11Z_ = [A_m/z 183_+A_m/z 199_], both obtained from the MS/MS channel at *m/z* 297.24. Each point represents a technical triplicate.

***The calibration curve for absolute quantification of FA 18:1 C=C isomers.*** A series of FA 18:1 isomer ([9Z] or [11Z] = 10, 25, 50, 125, 200, 225, 240 μM) was spiked with 25 μM of D17-FA 18:1 (9Z) as the internal standard (IS). Each mixture was epoxidized with excess mCPBA (200 mM) at 50 ^o^C for 1 hour and analyzed by LC-MS-tPRM. The calibration curve was constructed by plotting relative summed XIC area of the C=C diagnostic ions (A_FA_/A_IS_) against the concentration of each FA 18:1 isomer. A_FA_ denoted [A_m/z 155_+A_m/z 171_] for the 9Z isomer and [A_m/z 183_+A_m/z 199_] for the 11Z isomer, both obtained from the MS/MS channel at *m/z* 297.24; A_IS_ denoted [A_m/z 155_+A_m/z 171_] obtained from the *m/z* 314.35 MS/MS channel.

***The calibration curve for relative quantification of FA 18:3 ω-3 and ω-6 isomers.*** The total concentration of FA 18:3 was kept at 200 μM with the isomer molar ratios varied ([ω-3]/[ω-6] = 1/19, 1/3, 1/1, 3/1, and 19/1). Each sample was derivatized by 10 mM mCPBA at 50 ^o^C for 1 hour and then analyzed by LC-MS-tPRM, where the mono-epoxide of FA 18:3 (*m/z* 293.21) was targeted. The calibration curve was constructed by plotting the molar fractions ([ω-6]/([ω-3]+[ω-6]) %) against the fractions of the summed XIC area of C=C diagnostic ions (A_ω-6_/(A_ω-3_+A_ω-6_)%). Due to interferences of the co-eluted mono-epoxides, we chose two abundant C=C diagnostic ions of two mono-epoxides for each isomer. As a result, A_ω-3_ denoted [A_m/z 235(Δ15)_ + A_m/z 171(Δ9)_] and A_ω-6_ denoted [A_m/z 193(Δ12)_+ A_m/z 153(Δ9)_], both obtained from the MS/MS channel at *m/z* 293.21. The C=C diagnostic ions from Δ6 epoxide of *ω-6* epoxy-FA 18:3 and Δ12 epoxide of *ω-3* epoxy-FA 18:3 were neglected due to ion interference by the co-elution of these two mono-epoxides.

***Relative quantification of ω-6 and ω-9 isomer proportions of FA 20:3 containing lipids in 3T3-L1 adipocytes****.*

The total concentration of FA 20:3 was kept at 200 μM with the isomer molar ratios varied ([ω-6]/[ω-9] = 1/19, 1/3, 1/1, 3/1, and 19/1). Relative proportions the ω-9 isomer in FA 20:3 were estimated by AUC of the C=C diagnostic ions of the mono-epoxides. Specifically, the proportions of the ω-9 isomers (([ω-9]/([ω-6]+[ω-9])%) were estimated by A_ω-9_/(A_ω-6_+A_ω-9_) %, where A_ω-9_ denoted [A_m/z 221(Δ5)_+A_m/z 181(Δ8)_+A_m/z 179(Δ11)_] and A_ω-6_ denoted [A_m/z 157(Δ8)_+A_m/z 197(Δ11)_+A_m/z 221(Δ14)_], both obtained from the MS/MS channel at *m/z* 321.24.

**Epoxidation derivatization of biological lipid extracts using mCPBA**

The mCPBA reagent (200 mM) was prepared beforehand by dissolving mCPBA powders with reconstituted solution. The epoxidation derivatization of lipids was readily conducted by adding lipid extracts with excessed amount of mCPBA. For mono-unsaturated lipid analysis, the optimal condition of derivatization was conducted by mixing 10 uL of the lipid extract with 10 uL of the epoxidation reagent, and then the mixture was incubated at 50 ^o^C for at least 1 hour till the reaction completed; for poly-unsaturated lipids, 10 uL of the lipid extract with equal volume of the 10-fold diluted epoxidation reagent (20 mM mCPBA), incubated at 50 ^o^C for at least an hour. Subsequently, the derivatized sample was centrifuged and transferred to a sample vial for LC-MS analysis.

**LC-MS**

***Instrumental parameters.*** All LC-MS experiments were performed using a hybrid LTQ-Orbitrap Elite Mass Spectrometer (Thermo Scientific, Waltham, Massachusetts‎, US) couple with Shimazu LC-20A HPLC system (Shimazu, Kyoto, Japan). A heated electrospray ionization (HESI) probe was equipped as the ionization source with the following parameters: spray voltage at 4.0 kV in negative ionization mode; capillary temperature at 320 ^o^C; HESI heater temperature at 180 ^o^C; sheath gas flow at 30 (A.U.) and auxiliary gas at 10 (A.U.); the ion optics were tuned at *m/z* 283.26 ([M-H]^-^ ion of FA 18:0). Liquid chromatography was performed using a 2.6 μm SpeedCore C18 column (100 × 2.1 mm, Fortis technology, UK). A binary gradient was performed with mobile phase A of ACN/water (40/60, v/v) and mobile phase B of IPA/ACN (90/10, v/v). Both A and B solvents contain 1.0 mM ammonium acetate. The chromatographic separation was performed at room temperature (40 ^O^C) with flow rate of 0.1 mL/min. For conventional lipidomics, a 50-min gradient was established as: 0-3 min, 20% B; 3-8 min, 20-70% B; 8-30 min, 70-90% B; 30-30.5 min, 90-99% B; 40-41 min, 90-20% B; 41-50 min, 20% B. For C=C isomeric lipidomics of mono-unsaturated lipids, a 75-min gradient was set as: 0-5 min, 20% B; 5-50 min, 20-90% B; 50-51 min, 90-99% B; 51-60 min, 99% B; 60-61 min, 99-20% B; 61-75 min, 20% B; for poly-unsaturated lipids, a 90-min gradient was set as: 0-10 min, 5% B; 10-60 min, 5-90% B; 60-61 min, 90-99% B; 61-71 min, 99% B; 71-72 min, 99-5% B; 72-90 min, 5% B.

***Untargeted LC-MS/MS lipidomics.*** For conventional lipidomics analysis, the lipid extract was diluted 2 times and 3 μL of the sample was injected for LC-MS analysis. A top-10 data-dependent acquisition (DDA) method was applied. The scanning cycle of top-10 DDA consists one full FT-MS scanning with mass range of *m/z* 200-1500 and spectral resolution of 30,000, followed by 10 DDA IT-MS/MS scanning events. The minimal threshold to trigger MS/MS was 0.0. The dynamic exclusion was enabled after 2 repeat counts with 10 seconds of each repeat duration, exclusion list size of 500, and exclusion duration of 120 seconds. The exclusion mass width is set of +/- 10 ppm relative to reference mass. MS/MS spectra were acquired via ion activation type of CID with default charge state of 1, isolation width of *m/z* 2.0, normalized collision energy (NCE) of 40.0, activation Q of 0.250, and activation time of 10.00 ms. The maximum injection time for both full FT-MS and IT MS^n^ was set at 500 ms with auto-gain-control (AGC) of 3.00e+6 for full FT-MS scan and 3.00e+4 for IT MS/MS. Data was collected with Xcalibur 3.0 (Thermo).

***Creating targeted epoxy-lipid precursor ion lists for LC-MS-tPRM analysis.*** The LC-MS lipidomics data were processed via LipidSearch^TM^ (Thermo Scientific) in the search mode of “ProductSearch_Orbi”, in which the settings were optimized for lipid identification by high resolution MS1 precursors *m/z* and MS/MS fragments. Product ions with ion intensity above 1.0% threshold were searched and identified with precursor tolerance of 10.0 ppm and product tolerance of 0.5 Da. Both FA and all GPL classes were targeted for identification, and their negative-polarity ion adducts were chosen, including -H^+^ and +CH3COO^- +^ (for PC) adducts. Then the search results were processed in the alignment mode to generate the report, containing all the identified lipids, and the identified lipids with ID quality of D were removed. The alignment result was then imported into an excel file and processed through EpoxyFinder, a lab-built MATLAB-based software, to create a tPRM precursor ion list of the targeted mono-epoxidized unsaturated lipids for subsequent C=C isomer analysis. In general, this list included MS/MS and MS/MS/MS precursor ion *m/z* parameters of mono-epoxidized unsaturated lipids and the corresponding NCE.

***C=C isomeric lipidomics via LC-MS-tPRM.*** For C=C isomeric lipidomics, 3 μL of the epoxidized lipid extract was injected for LC-MS-tSRM analysis. A top-25 tPRM scanning module was applied, where each scanning cycle contained one full FT-MS scan event with mass range of *m/z* 200-1500 and spectral resolution of 30,000, followed by 25 IT-MS^n^ scanning events. To establish the tPRM environment, the IT-MS^n^ acquisitions were only triggered by the top-25 intense targeted precursors, *i.e.* the targeted epoxy-lipid ions, that were detected in the full FT-MS spectrum. The maximum injection time for both full FT-MS and IT MS^n^ was set to 200 ms with AGC of 3.00e+6 for full FT-MS scan and 3.00e+4 for IT MS^n^. The ion activation methods of CID included default charge state of 1, isolation width of *m/z* 2.0, activation Q of 0.250, and activation time of 10.00 ms; NCE of was set to 40.0 for MS/MS of epoxy-FAs; for epoxy-GPLs, NCE of 55.0 was chosen for MS/MS to enhance the signals of epoxy-FA daughter ions, and NCE of 40.0 was chosen for MS^3^ of the epoxy-FA daughter ion. Data was collected with Xcalibur 3.0 (Thermo).

***Identification and quantification of C=C isomers by LC-MS-tPRM analysis.*** The resulting LC-MS-tPRM data of the epoxidized lipid extracts were processed via Xcalibur Qual Browser (Thermo Scientific, Waltham, Massachusetts‎, US)). The presence of each targeted epoxy-lipid was confirmed by the observed precursor ion in full-FTMS spectra. For each identified epoxy-lipid, its potential C=C isomers were identified by the extracted ion chromatograms (XICs) of C=C diagnostic ion pairs in MS/MS or MS^3^ during the proximal retention time. The presence of each C=C isomer was confirmed if its C=C diagnostic ion pair was observed and well-aligned in the retention time. Relative quantification of C=C isomers was accessed by calculating XIC area of C=C diagnostic ion pairs. Quantification was not accessed for C=C isomers when their XIC area of C=C diagnostic ions were unable to be extracted due to insufficient signals or incomplete detection of chromatographic peaks. In such cases, only the identification results were shown.

**Tissue preparation for DESI MSI**

Lung tissues containing metastatic tumors from a breast cancer mice model were collected from Dr. Tang-Long Shen’s lab in National Taiwan University. These mice were handled in accordance with a protocol approved by the Institutional Animal Care and Use Committee of National Taiwan University (IACUC approval NO. NTU104-EL-00003). Mouse brain and kidney were purchased from BioLASCO (Taipei, Taiwan), whereas the gender and age of the ICR mice were not specified. All the tissues were stored in -80 ^o^C before sectioning. Frozen sectioning was done by using a frozen microtome (LEICA, CM1900). Prior to the sectioning, tissues were flash-frozen by liquid nitrogen and kept in the cold tome at -20 ^o^C. The tissues were placed on the cutting stage by optimal cutting temperature (OCT) compound and sectioned into 20-µm think sections and thaw-mounted on silane-coated glass slides (76 × 26 × 1 mm; Yeong Jyi Chemical Apparatus, Taiwan). The slides were stored in -80 ^o^C before imaging analysis.

**DESI lipid C=C isomer imaging**

***In situ epoxidation pretreatment of tissue sections.*** The frozen tissue section was dried in a vacuum desiccator for at least 1 hour before epoxidation pretreatment. The *in situ* epoxidation was achieved by spraying tiny droplets of mCPBA solution (100 mM in MeOH) onto the section. The mCPBA application used a commercial airbrush (RH-BS, Prona, Taiwan) containing a flow-adjusting knob and a spray nozzle of 300-500 μm in diameter. The airbrush was connected to a commercial air compressor (CPM-280A air compressor, Xian-Ying, Taiwan) which contained a gas-storage bottle, a water-filtering bottle, and air pressure adjusting valves. Briefly, *in situ* epoxidation was carried out by spraying a thin layer of mCPBA onto the tissue section before C=C imaging analysis. The detailed method is described in **Section S7-3**. A demonstration video is provided at:

<https://drive.google.com/drive/folders/13Plb4-aY0NnMpS6pT9UURBMOWCDgkQ-N/>

***Instrumental parameters.*** All DESI-MS experiments were performed using a hybrid LTQ-Orbitrap Elite Mass Spectrometer (Thermo Scientific, San Jose, CA) coupled with Omni Spray 2D Ion Source (Prosolia Inc., Indianapolis, U.S.A.) as the ionization source and 2D stage. The spectra were acquired in negative ion mode with spray voltage of 3.5 kV. DESI sprayer was optimized with the angle of spray head of 56^o^, head height of 1.8 mm, head to inlet distance of 3.0 mm, and nebulizing gas pressure of 170 psi. MS ion optics were tuned at *m/z* 281 by continuously injecting FA 18:1 standard solution. A biocompatible solvent, DMF/ACN (v/v = 1/1), with flow rate of 2 μL/min was chosen as the spray solvent. All images were acquired with a stage step size of 200 μm. The full-scan MS images were acquired in FT-MS mode with mass range of *m/z* 200-1000. The C=C imaging spectra were obtained by continuously acquiring IT-MS^n^ spectra (n = 2 for FA; n = 3 for GPL) throughout the tissue section. The isolation width of parent ion was *m/z* 3.0 for MS/MS and *m/z* 2.5-5.0 for MS^3^. NCE value was set at 40.0-55.0 for MS/MS and 40.0 for MS^3^. Data was collected with Xcalibur 3.0 (Thermo).

***Data visualization.*** The resulting .RAW data were converted into .mzXML format by MSconvert (ProteoWizard 3.0.5471; <http://proteowizard.sourceforge.net/>) and then processed by a lab-built MATLAB algorithm. For full-MS imaging, ion signals of each lipid were extracted by a *m/z* width of +/- 0.01 with the center at the theoretical *m/z* value. For C=C MS images, C=C diagnostic ion signals were extracted by a *m/z* width of +/- 0.5 with the center at the theoretical *m/z* value. To reduce background noise, ion signals of noise-level intensity were excluded and the ion images were filtered by a 2D Gaussian function. The images were segmented according to the tissue margin indicated by optical H&E staining image, and visualized via fraction distribution image (FDI) by plotting fractional intensity (FI) of each isomer. FI is determined in each pixel by calculating summed ion intensities of the C=C diagnostic ions of the denoted C=C isomer as a fraction of diagnostic ion signals arising from all isomeric species. For example, for an unsaturated lipid consisting two C=C isomers, Δm and Δn, the calculation could be generalized as the following equation:

$$\text{FI of ∆m =} \frac{\sum\text{Int.}_{\text{∆m C=C ions}}}{\sum\text{Int.}_{\text{∆m C=C ions}}\text{+}\sum\text{Int.}_{\text{∆n C=C ions}}}\text{ }$$

More specifically, for PG 18:1_16:0 containing two C=C isomers, Δ9 and Δ11, the calculation became:

$$\text{FI of}\text{ }\text{ PG 18:1(∆11)\_16:0 =} \frac{\text{Int.}_{\text{m/z }\text{183}}\text{+}\text{Int.}_{\text{m/z}\text{ 199}}}{\text{Int.}_{\text{m/z}\text{ 155}}\text{+}\text{Int.}_{\text{m/z}\text{ 171}}\text{+} \text{Int.}_{\text{m/z}\text{ 183}}\text{+}\text{Int.}_{\text{m/z}\text{ 199}}}$$

For FA 18:1, because the MS/MS spectrum of epoxy-FA 18:1 contained a background signal at *m/z* 183, the calculation was modified by calculating only the aldehyde-form ions as below:

$$\text{FI of FA 18:1(∆11) =} \frac{\text{Int.}_{\text{m/z}\text{ 199}}}{\text{Int.}_{\text{m/z }\text{171}}\text{+} \text{Int.}_{\text{m/z}\text{ 199}}}\text{ }$$

***Quantification of C=C isomer proportion.*** To access C=C isomer proportion in each tissue region, the FDI image was manually segmented into distinct tissue regions with respect to the region margins, which were determined by H&E optical image. The isomer ratio in each pixel in the FDI image, except for the data point right in the margin between two regions, was extracted for calculation of mean, standard deviation, and statistical analysis.

**Statistical information.** All error bars in this study represent standard deviation (SD). All statistical comparisons in this study were performed using two-tailed t-test.

**Data availability.** The raw data that support the main findings of this study are available at:

<https://drive.google.com/drive/folders/13Plb4-aY0NnMpS6pT9UURBMOWCDgkQ-N/>

**Code availability.** EpoxyFinder software package and the algorithm are available at:

<https://drive.google.com/drive/folders/13Plb4-aY0NnMpS6pT9UURBMOWCDgkQ-N/>

**S1. Identification of C=C positions in unsaturated lipids using MELDI-MS/MS**

**S1-1. Monounsaturated GPLs**

The C=C position in a fatty acyl chain of a monounsaturated glycerophospholipid (GPL) could be identified by MS^3^ analysis of its epoxidized product. Due to instrumental restriction of Orbitrap Elite Mass Spectrometer that we used in this study, detection of diagnostic ions (usually in *m/z* < 250) was not available in CID-based MS/MS due to low-mass cutoff during ion isolation in ion trap (IT). Therefore, identification of C=C positions in GPLs required sequential CID (i.e. MS^3^) in IT, and then the MS^3^ fragments could be detected in either the IT or FT detector.

The concept was first demonstrated by PE 16:0/18:1 (9Z) in **Main** **Fig. 1f**, and herein further elaborated by PS 16:0/18:1 (9Z), PC 16:0/18:1 (9Z), and PA 18:1 (9Z)/18:1 (9Z) (**Fig. S1**). The ionization of PC in negative ion was promoted by the addition of ammonium acetate (NH_4_OAc) in ESI solution or LC mobile phase, enabling PC molecules to be ionized in the form of the acetate adduct ([M+OAc]^-^). To sum up, in the MS/MS spectrum of an epoxy-GPL in negative ion mode, daughter ions from its two fatty acyl chains were readily observed. Subsequently, targeting CID to the epoxidized FA daughter ion enabled the detection of the C=C diagnostic ion pair, which was the same with that identified from the free epoxidized FA. Taking PS 16:0/18:1 for example, in the MS/MS spectrum of its epoxide product, in addition to the neutral loss fragment (*m/z* 689.47), we found two daughter ions, *m/z* 255.23 and *m/z* 297.24, representing the FA 16:0 chain and the epoxy-FA 18:1 chain, respectively (**Fig. S1a-i**). Subsequently, MS^3^ of the epoxy-FA 18:1 revealed its C=C position at Δ9 by the diagnostic ion pair (**Fig. S1a-ii**).

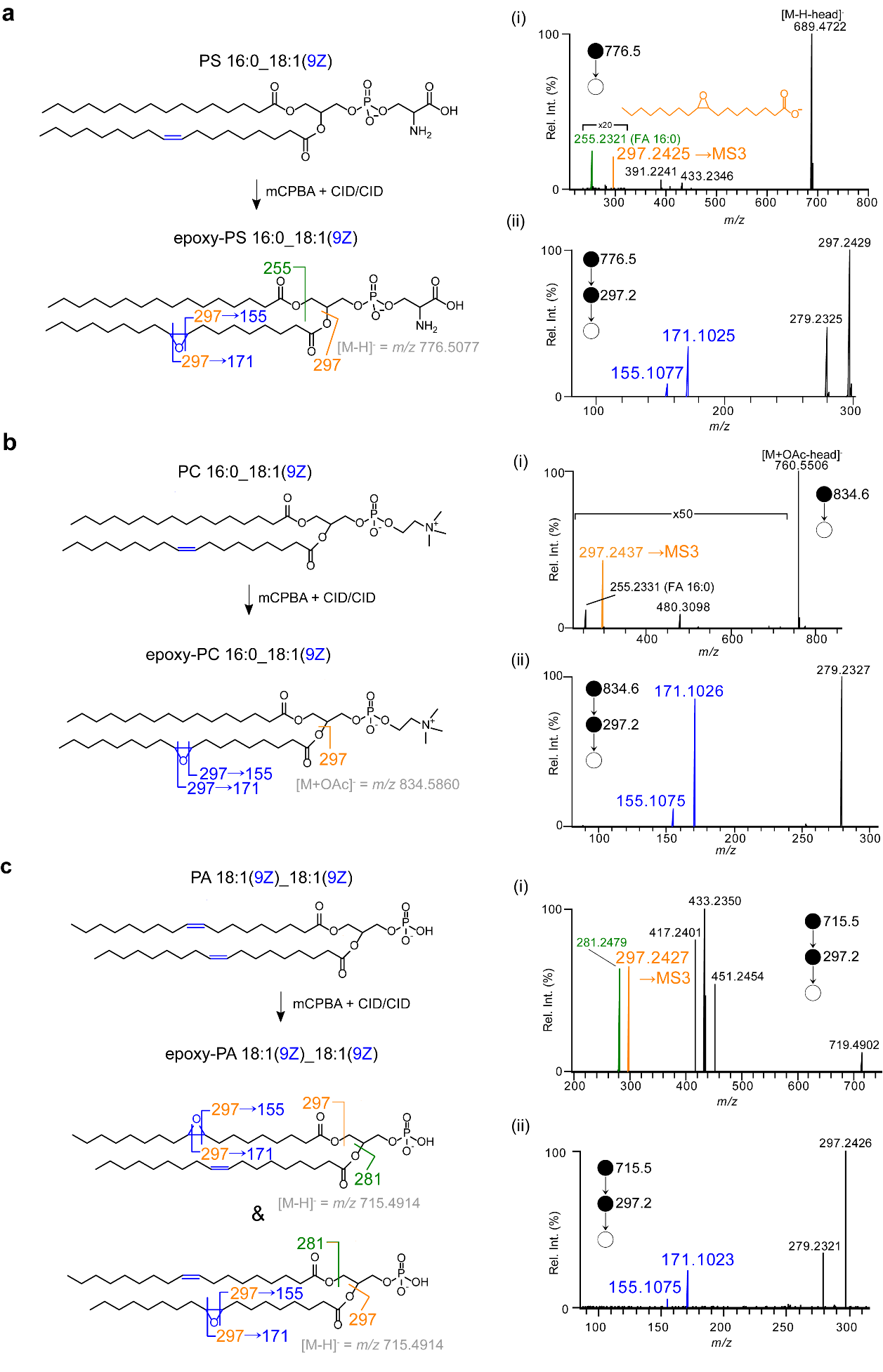

**Figure S1. Pinpointing C=C positions in unsaturated GPLs by MELDI-MS/MS/MS.**

Unsaturated GPLs, including (a) PS 16:0_18:1(9Z), **(b)** PC 16:0_18:1(9Z), and **(c)** PA 18:1(9Z)_18:1(9Z), were epoxidized by mCPBA and then subjected to FT-MS/MS and FT-MS/MS/MS analysis. The C=C position of a GPL was subsequently revealed by the C=C diagnostic ion pair in the MS^3^ spectrum of the epoxidation product.

**S1-2. C=C geometric isomers**

The epoxides of C=C geometric isomers were separable in RPLC but their MS/MS spectra were indistinguishable. A mixture of FA 18:1 (9Z) and FA 18:1 (9E) was derivatized by mCPBA and subjected to LC-MS/MS analysis. As shown in **Fig. S2a**, two peaks were found in the XIC of *m/z* 297.24, the dehydrated ion of epoxy-FA 18:1, and their annotations were confirmed by individual 9Z and 9E standards. However, the epoxides gave two identical FT-MS/MS spectra (**Fig. S2b and c**). Therefore, in our further study, C=C positions would be only annotated in Δ-nomenclature without referring the C=C geometry.

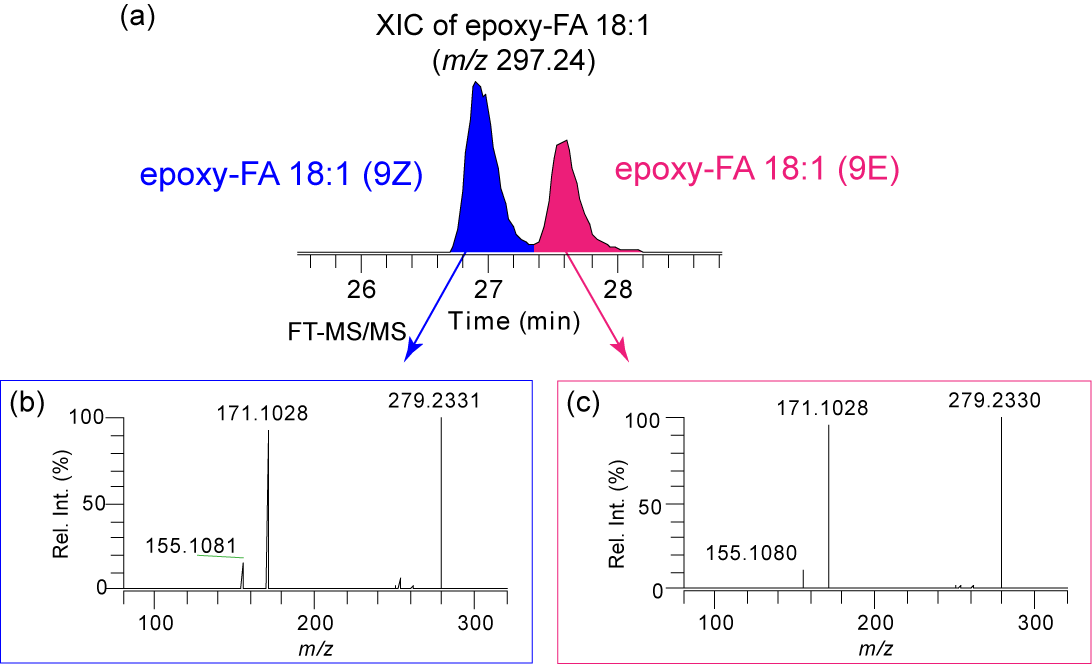

**Figure S2. LC-MS/MS analysis of FA 18:1 9Z and 9E geometric isomers.**

The extracted ion chromatogram (XIC) of epoxy-FA 18:1 showed two peaks of epoxidized 9Z and 9E isomers in **(a)**, and their FT-MS/MS spectra were found identical (**b-c**).

**S1-3. Polyunsaturated FAs and GPLs**

**S1-3-1. FT-MS/MS spectra of fully epoxidized PUFAs**

**Figure S3** showed FT-MS/MS spectra of fully epoxidized PUFAs. Generally, CID fragmentation could still occur at epoxide positions, while the spectra became more complicated than those of epoxy-MOFAs. In addition, the diagnostic ions of fully epoxidized PUFAs diminished when the number of C=C bonds increased to 4, making it difficult to annotate the C=C positions (**Fig. S3d**).

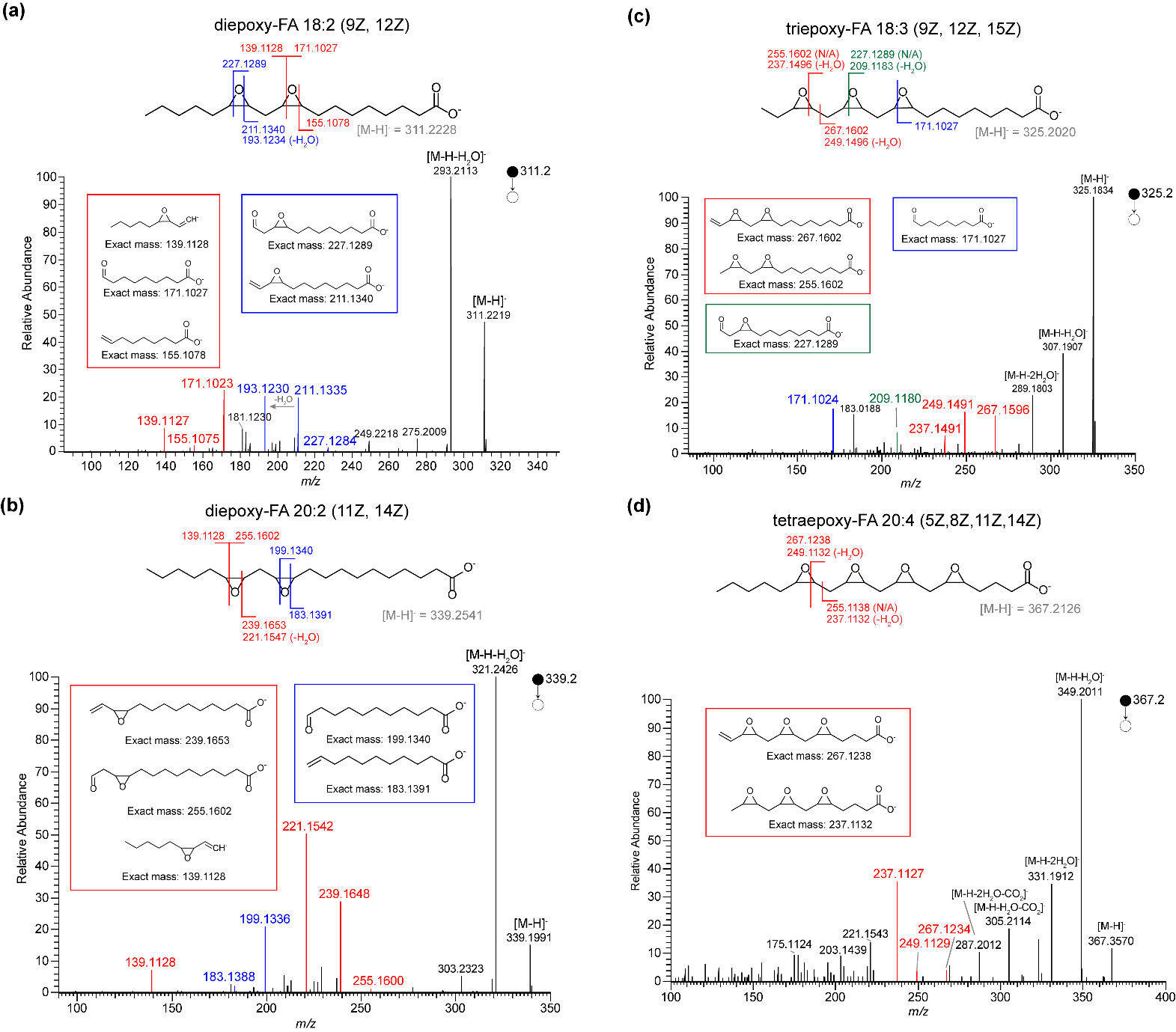

**Figure S3. FT-MS/MS spectra of fully epoxidized PUFAs**

Polyunsaturated fatty acids (PUFAs) including **(a)** FA 18:2 (9Z, 12Z) **(b)** FA 20:2 (11Z, 14Z) **(c)** FA 18:3 (9Z, 12Z, 15Z) **(d)** FA 20:4 (5Z, 8Z, 11Z, 14Z) were epoxidized in solution with excess mCPBA, and the FT-MS/MS spectra of their fully epoxidized products were shown.

**S1-3-2. Identification of C=C positions in mono-epoxidized PUFAs by LC-MS/MS**

Multiple C=C positions of a PUFA molecule can be identified by LC-MS/MS interrogation of its mono-epoxidized products. In addition to FA 18:2 and FA 18:3 that have been demonstrated in **Main Fig. 3**, herein more PUFA standards, including ω-6 FA 20:2, ω-6 FA 20:3, ω-9 FA 20:3, ω-6 FA 20:4, and ω-3 FA 20:5, were thoroughly investigated by MELDI-LC-FTMS/MS. The spectral dataset was shown in **Figure S4**, where the MS1 XICs of the mono-epoxidized products, the corresponding FT-MS/MS spectra, the CID scheme, and the proposed C=C diagnostic ions were shown. Interestingly, when the C=C number of a PUFA was less than 4 (e.g. FA 20:2 and 20:3 in **Figure S4a and b**), the yield of its multiple epoxides were almost similar, suggesting that the epoxide could tag each C=C bond with an equal tendency. When the C=C number increased up to 4 or more (e.g. FA 20:4 and 20:5 in **Figure S4c and d**), the epoxidation might prefer at C=C bonds closer to the methyl end (i.e. the higher Δ number). Furthermore, regarding the issue of co-elution of epoxides, as shown by the merged peaks in the MS1 XIC, the MS/MS spectrum of each epoxide could still be extracted in the most cases, thereby allowing us to deconvolute a single MS1 peak into multiple epoxides in the MS2 level.

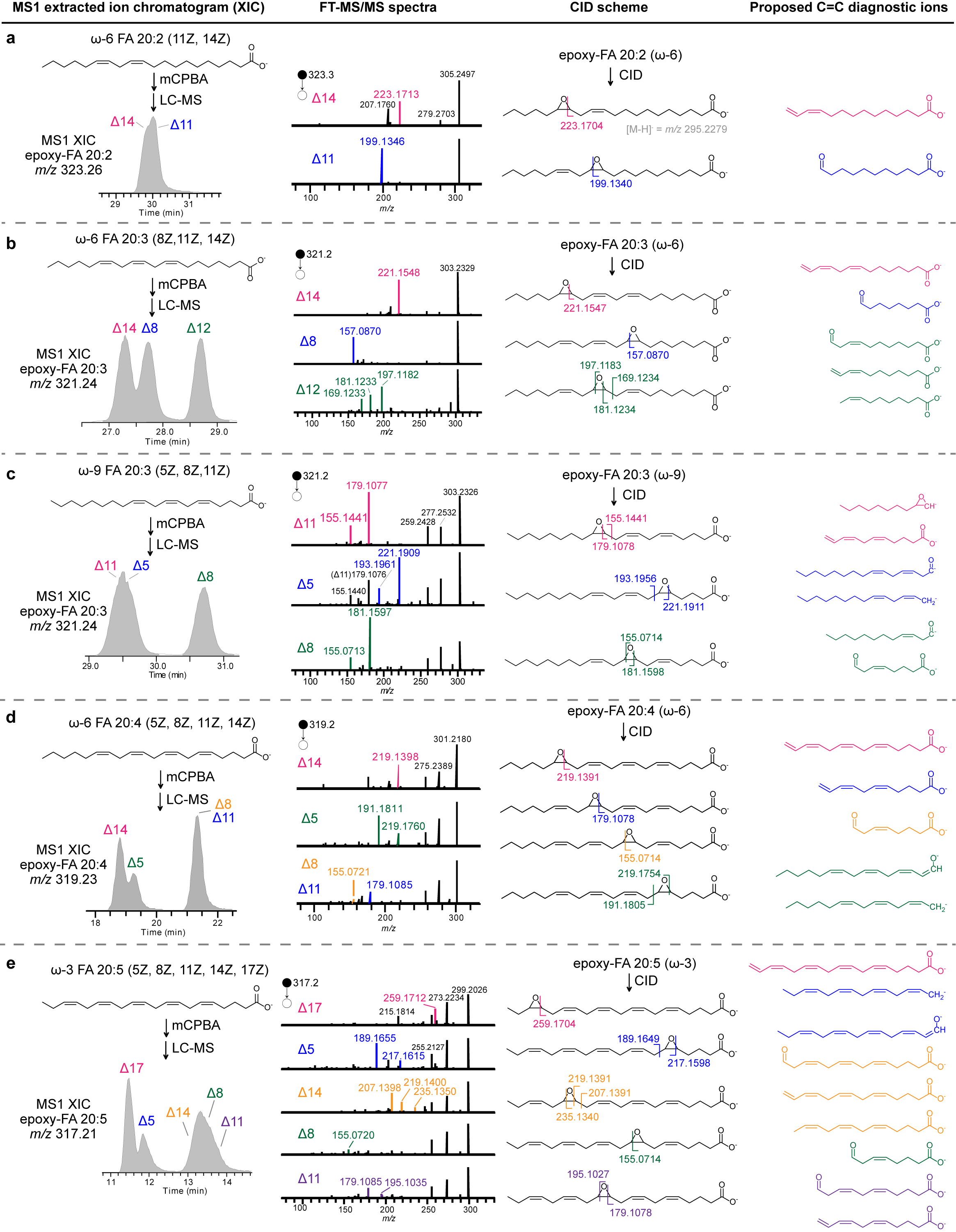

**Figure S4. Pinpoint C=C positions of polyunsaturated fatty acids by MELDI-LC-MS/MS.** Polyunsaturated fatty acids (PUFA), including **(a)** ω-6 FA 20:2, **(b)** ω-3 FA 18:3, **(c)** ω-6 FA 20:4, and **(d)** ω-3 FA 20:5, were epoxidized by mCPBA and subjected to LC-MS/MS analysis. Multiple C=C positions of a given PUFA was identified by its mono-epoxidized products and the corresponding MS/MS spectra.

**S1-3-3. The CID scheme of mono-epoxidized PUFAs**

The CID scheme of the mono-epoxidized PUFA was discussed here. We found that the epoxide structure in an epoxy-PUFA was still labile upon CID, though the resulting MS/MS spectrum was not fully favorable to form an aldehyde/alkene diagnostic ion pair as epoxy-MOFA did. Instead, the CID fragmentation of an epoxy-PUFA favors the side adjacent to the other unsaturated site(s), which might be due to the generation of products with conjugated π-bond systems upon CID. Taking ω-6 FA 20:2 (11Z, 14Z) as an example, one of its mono-epoxidized product, Δ14-epoxy-FA 20:2, might yield a C=C diagnostic ion pair, including a diene ion of *m/z* 223.1704 with a conjugated system and an aldehyde ion of *m/z* 239.1653 (**Fig. S5a**), based on the previous knowledge in diagnosing MOFA-derived epoxides. In its MS/MS spectrum, however, only the conjugated diene ion was found. For the other mono-epoxidized product, Δ11-epoxy-FA 20:2, again a diagnostic ion pair (an alkene ion of *m/z* 183.1931; an aldehyde ion of *m/z* 183.1931) was proposed, but only the aldehyde ion was observed in the MS/MS spectrum (**Fig. S5b**). This was rationalized as the bond breaking of the epoxide that resulted in the formation of the aldehyde ion might form a neutral conjugated diene yet unable to be detected**.** In this regard, the CID tendency to form a conjugated system was thus proposed. Another two case studies were also provided in **Fig. S5c and d**, which also supported the proposed CID scheme.

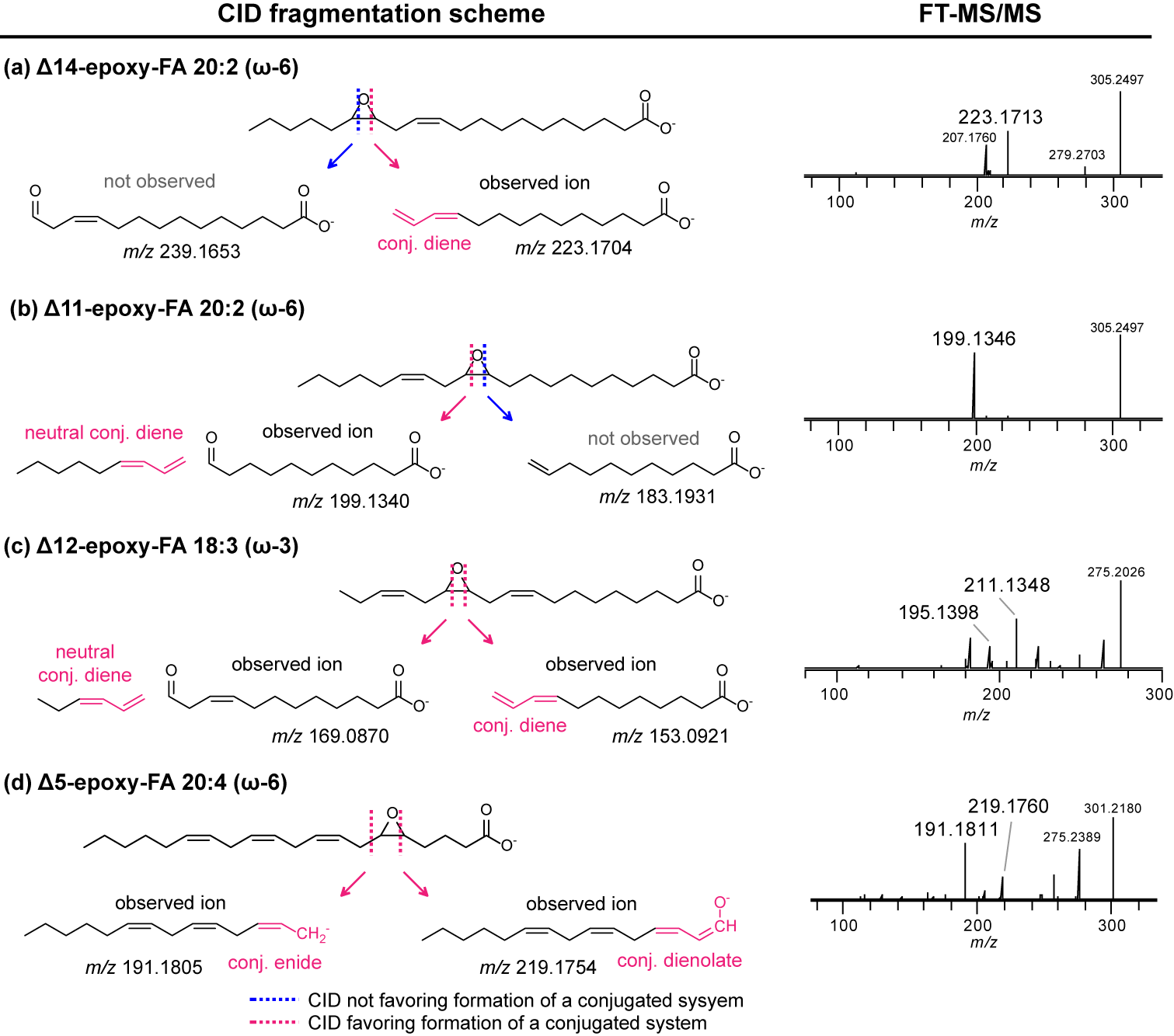

**Figure S5. The CID scheme of fragmentation of mono-epoxidized PUFAs**

The CID schemes and the FT-MS/MS spectra of mono-epoxidized PUFAs, including **(a)** the Δ14 epoxide and **(b)** the Δ11 epoxide of FA 20:2 (11Z, 14Z), **(c)** the Δ12 epoxide of ω-3 FA 18:3 **(d)** the Δ5 epoxide of ω-6 FA 20:4. The cleavage sites favor or disfavor the formation of fragments with conjugated systems were indicated.

**S1-3-4. Identification of C=C positions in mono-epoxidized GPLs by LC-MS/MS/MS**

Identification of multiple C=C positions in a PUFA chain in a GPL could be achieved by MS^3^ interrogation to the mono-epoxidized GPL, which was similar to the cases of monounsaturated GPLs. We demonstrated the concept by phosphatidylinositol (PI) 18:0/20:4 (5Z, 8Z. 11Z, 14Z), a standard GPL containing a saturated FA 16:0 chain and an ω-6 FA 20:4 chain. The compound, after epoxidation by limited mCPBA, was subjected to LC-MS, in which four mono-epoxidized products (epoxy-PI 38:4, [M-H]^-^ = *m/z* 901.5448) were separated in MS1 level (**Fig. S6a**). In their corresponding FT-MS/MS spectra, the presence of epoxidation tagging at C=C bonds was clearly indicated by the fragment ion of epoxy-FA 20:4 (*m/z* 319.2279) (**Fig. S6b**). However, the four MS/MS spectra were near-identical and C=C positions could not be resolved. We further applied FT-MS^3^ analysis to the epoxy-FA 20:4 fragments, resulting in four different MS^3^ spectra that allowed us to pinpoint where the epoxide tagged (**Fig. S6c**). In addition, such MS^3^ fragmentation was consistent with that of free ω-6 FA 20:4 (**Fig. S6e**). Finally, we showed that when IT was used to detect the MS^3^ fragments, the better spectral quality was obtained (**Fig. S6d**). With this regard, IT-MS^3^ was applied in our further study to ensure the high quality of MS^3^ annotation of polyunsaturated GPLs.

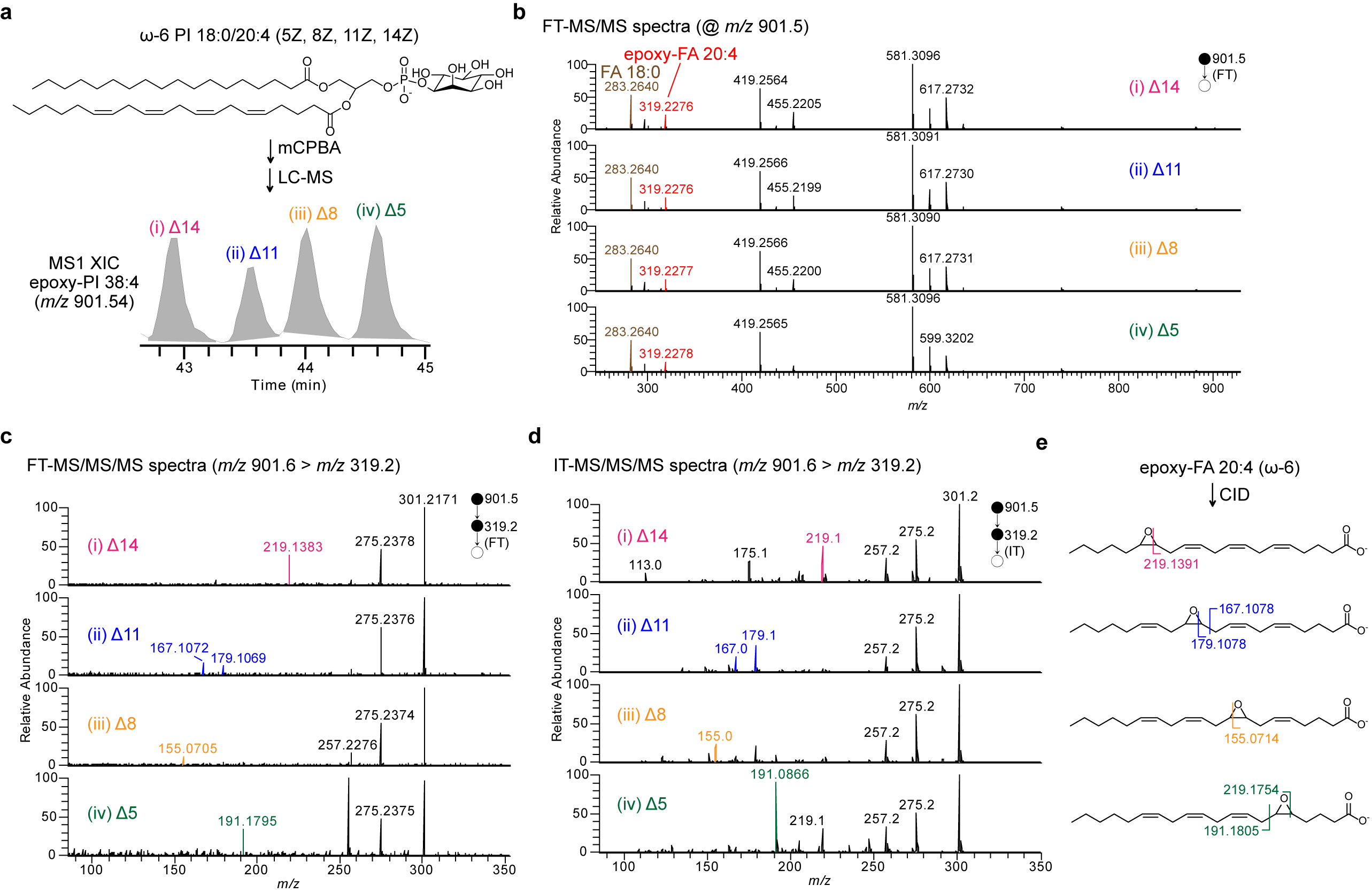

**Figure S6. MELDI-LC-MS/MS/MS enabled identification of C=C positions in ω-6** **PI 18:0/20:4.**

**(a)** The LC-MS XIC of the mono-epoxidized PI 18:0/20:4, showing four separated peaks. Their corresponding **(a)** FT-MS/MS spectra **(c)** FT-MS/MS/MS spectra **(d)** IT-MS/MS/MS spectra were provided. **(e)** The CID scheme of the mono-epoxidized FA 20:4 (ω-6).

**S2. Features of mCPBA epoxidation**

**S2-1. Epoxide yield of unsaturated FAs and GPLs**

Epoxidation using mCPBA allowed the derivatization of an unsaturated lipid to form an epoxide. Here the epoxide yield was investigated by a series of mono-unsaturated lipids reacted with excess mCPBA, and the experimental method was described. First, the lipid standard (100 µM) was diluted with equal volume of methanol (MeOH), analyzed with LC-MS, and the area under curve (AUC) of each non-epoxidized lipid in extraction ion chromatogram (XIC) was obtained, denoted by A_lipid_. Secondly, the lipid standard was added with equal volume of mCPBA solution (200 mM in MeOH), incubated at 50 ^o^C for at least 1 hour, analyzed with LC-MS, and AUC of its epoxidation product was obtained, denoted by A_epoxy-lipid_. Finally, the epoxide yield was estimated by A_epoxy-lipid_ / A_lipid_. An investigation of PC 16:0/18:1 (9Z) was shown in **Fig. S7**, and the overall results were summarized in **Table S1**, where the epoxide yield of each lipid was calculated from a single experiment. In brief, the epoxide yield of a MOFA was estimated to be above 70 %, and the epoxide yield of different subclasses of GPLs varied in 26~91 %. Note that the yield was not estimated by the purified compound and could be affected due to difference in ionization efficiencies between unsaturated lipids and their epoxides. However, difference of epoxide yield among lipid subclasses would not affect the quantification of C=C isomer composition in a given species.

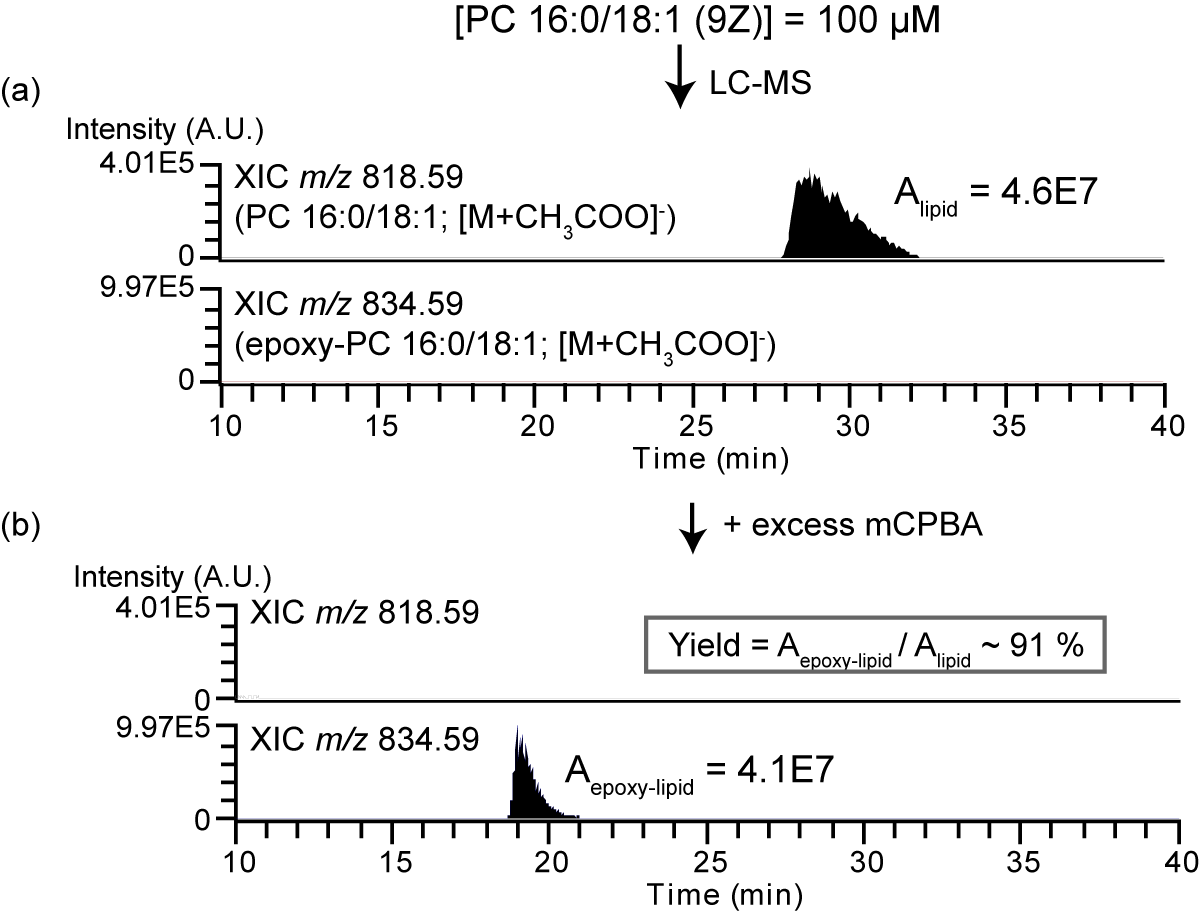

**Figure S7. Estimation of the epoxide yield of PC 16:0/18:1 (9Z) with mCPBA epoxidation.**

The XICs of PC 16:0/18:1(9Z) (100 µM) **(a)** prior to epoxidation and **(b)** after epoxidation using excess mCPBA were shown. The epoxidation yield was estimated by the area under curve ratio (A_epoxy-lipid_ / A_lipid_).

**Table S1. The estimated epoxide yields of mono-unsaturated lipids.**

| **Mono-unsaturated lipid** | **Targeted mono-epoxy-lipid** | **Epoxide yield (%)** |
| --- | --- | --- |
| FA 18:1 (9Z) | epoxy-FA 18:1 (9Z) | 73 |
| D_17_-FA 18:1 (9Z) | epoxy-D_17_-FA 18:1 (9Z) | 78 |
| PC 16:0/18:1 | epoxy-PC 16:0/18:1 | 91 |
| PE 16:0/18:1 | epoxy-PE 16:0/18:1 | 29 |
| PS 16:0/18:1 | epoxy-PS 16:0/18:1 | 26 |

**S2-2. Limit of detection of C=C diagnostic ions**

Limit of detection of MELDI-LC-MS to detect C=C diagnostic ions was evaluated here.

**MOFA**

A series of D_17_-FA 18:1 (9Z) samples (10^-3^, 10^-2^, 10^-1^, 10^0^, 10^1^, and 10^2^ µM) were derivatized by excess mCPBA (100 mM), incubated at 50 ^o^C for an hour, and analyzed by targeted LC-MS/MS, in which both FT-MS/MS and IT-MS/MS (with MS1 parent ion at *m/*z 314.35) were set. Then, the summed AUC of the diagnostic ion pair of Δ9 (*m/z* 155 and *m/z* 171) were plotted against the lipid concentrations, as shown in **Fig. S8a and S8b** for FT-MS/MS and IT-MS/MS, respectively. The good linearity was obtained in both FT and IT mode (R^2^ ~ 0.98), and the diagnostic ions were detectable when the initial FA concentration was 1 nM.

**MOFA-derived GPL**

For the GPL containing an MOFA chain, several standards, including PE 16:0/18:1 (9Z), PS 16:0/18:1 (9Z), and PC 16:0/18:1 (9Z) (10^-2^, 10^-1^, 10^0^, 10^1^, and 10^2^ µM) were examined similarly except that MS^3^ was used. In general, the LOD of diagnostic ions was estimated at sub-µM level (**Fig. S8c-h**), while the LOD was increased to µM level in FT-MS^3^ for PC (**Fig. S8g**), which might be due to the ion loss during longer transmission toward the Orbitrap detector. In comparison to free MOFAs, the LOD of GPL was increased by approximately 2 orders of magnitude. Since the epoxide yields for both free FA and GPL were estimated to be at the same order of magnitude (**Table S1**), the increased LOD of GPL could be mainly explained by (i) the requirement of MS^3^ for GPLs (ii) low abundant epoxy-FA fragments in MS/MS spectra. For the later, we found that the fragment from neutral loss of the headgroup would dominate the MS/MS spectrum (see the MS/MS spectra of the epoxidized PC and PS in **Fig. S1**). With this regard, we further showed that the LOD of GPL could be improved by MS^4^. In specific, we first applied MS/MS to the epoxidized parent ion, MS^3^ to the head-loss fragment, and finally MS^4^ to the epoxy-FA fragment ion to yield C=C diagnostic ions. As a result, the LOD could be improved by an order of magnitude (**Fig. S8g and h**). However, the MS^4^ setting was unable to be included in a LC-MS-tPRM method in Thermo Xcalibur software due to the vendor’s design, but it could still be feasible in a targeted manner (i.e. targeted LC-MS^4^). Therefore, we compromised with using MS^3^ for all subclasses of GPLs in LC-MS-tPRM analysis.

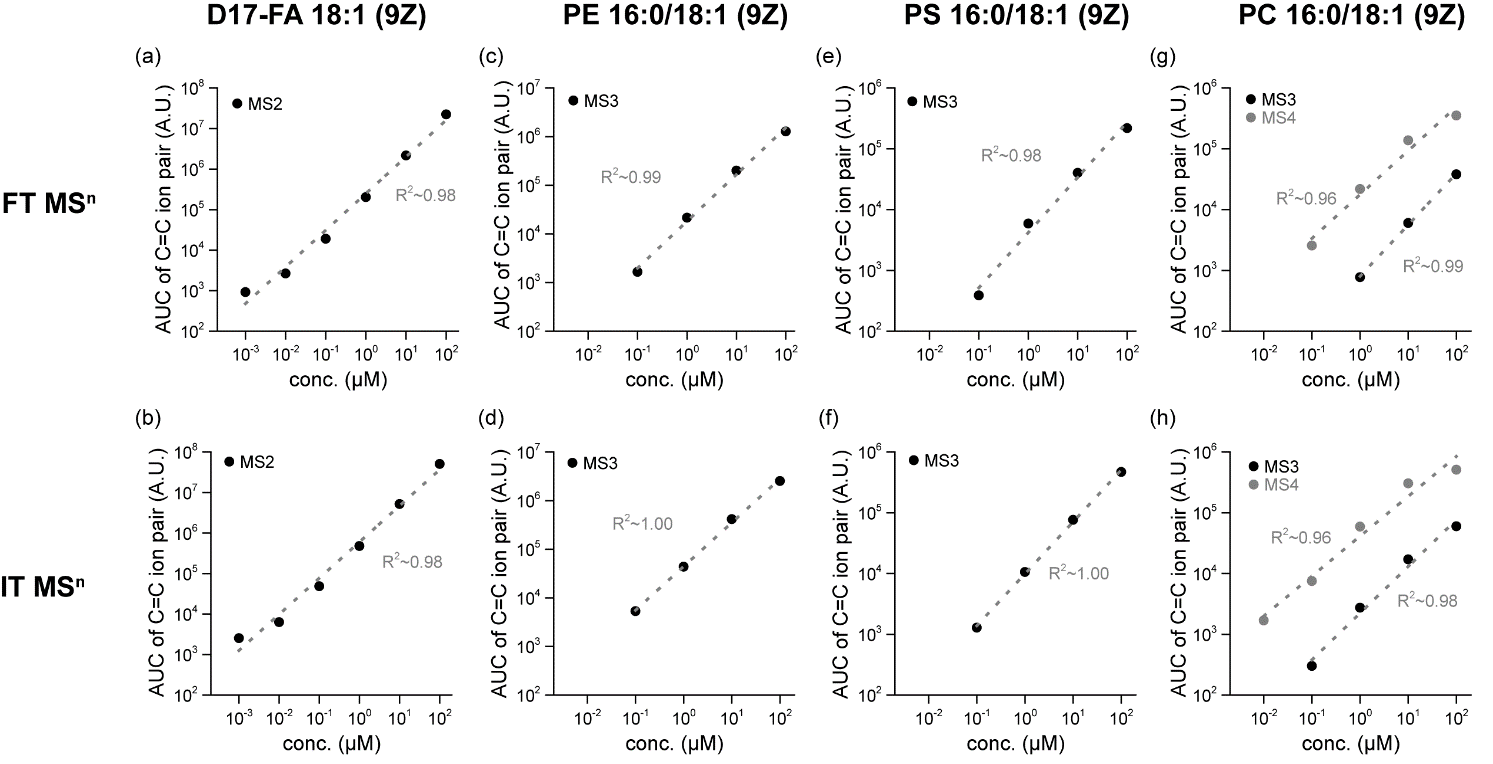

**Figure S8. LOD of C=C diagnostic ions in the mono-unsaturated FA and GPL.**

Serial dilutions of each unsaturated lipid standard (10^-3^ ~ 10^2^ µM for FA; 10^-2^ ~ 10^2^ µM for GPL), all containing a 9Z C=C bond, were subjected to the targeted MELDI-LC-MS^n^ analysis, and their corresponding AUCs of the diagnostic ion pair (*m/z* 171 and *m/z* 155) were shown in relation with the initial lipid concentration. Ion signals detected in FT mode (upper panel) and IT mode (lower panel) were both examined. The plots were shown in the log_10_ scale.

**PUFA**

LOD of C=C diagnostic ions for PUFA was evaluated by a standard PUFA, ω-6 FA 20:3 (8Z, 11Z, 14Z). The LC-MS/MS spectra of its three mono-epoxidized products were previously studied in Fig XX. A series of ω-6 FA 20:3 (10-3, 10-2, 10-1, 100, 101, and 102 µM) were derivatized using the previously developed protocol, in which the PUFA was mildly epoxidized with mCPBA (10 mM) at 50 ^o^C for an hour, and analyzed with LC-MS/MS targeting the three mono-epoxidized products (*m/z* 321.24). The LOD of C=C diagnostic ions obtained from each of the three epoxides was first examined separately (**Fig.S9a-f)** and then combined as a general description for the LOD of ω-6 FA 20:3 (**Fig. S9g and h**). As a result, while LOD for individual epoxides varied in FT-MS/MS, we concluded the LOD was about 1 nM. Also, since a lower LOD in IT-MS/MS could be expected, diagnostic ions of the three epoxides were constantly detectable from the PUFA sample of 1 nM.

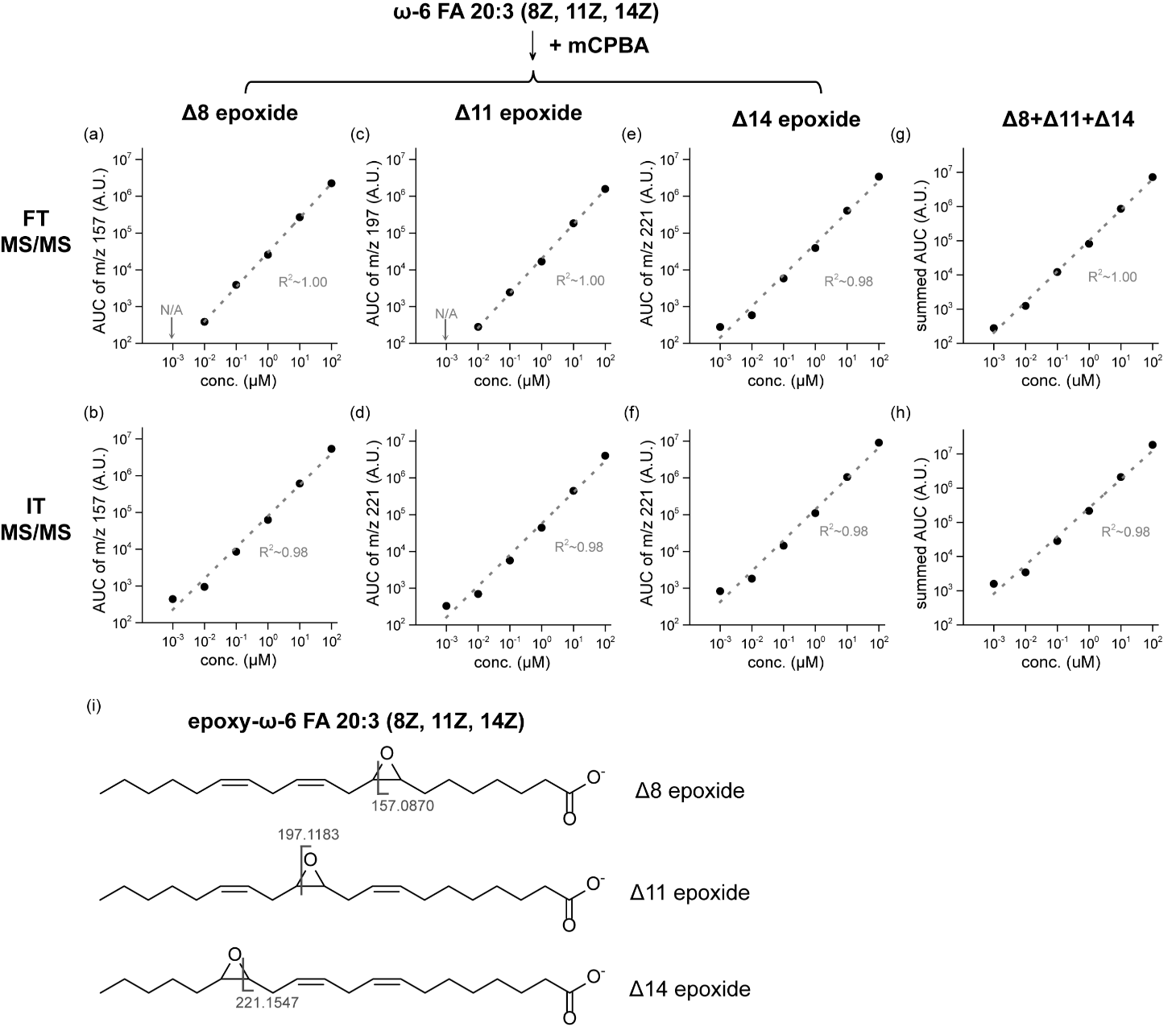

**Figure S9. LOD of C=C diagnostic ions for ω-6 FA 20:3.**

Serial dilutions of ω-6 FA 20:3 (10^-3^ ~ 10^2^ µM) were subjected to the targeted MELDI-LC-MS/MS analysis. For each epoxide, AUC of its most abundant C=C ion was extracted, and plotted against the initial FA concentration **(a-f)**. Ion signals detected in FT mode (upper panel) and IT mode (lower panel) were both examined. Results of the three epoxides were further combined and shown in **(g)** and **(h)**. The plots were shown in the log_10_ scale. **(i)** The CID scheme of three targeted mono-epoxidized ω-6 FA 20:3, where the most abundant diagnostic ion was indicated.

**S2-3. Controlling epoxidation for studying PUFA C=C isomers**

In the analysis of a PUFA, epoxidation should be carefully controlled in order to keep a high proportion of its mono-epoxidized product. The derivatization protocol was evaluated with three PUFAs: ω-6 FA 18:3, ω-3 FA 18:3, and ω-9 FA 20:3. We showed that for 200 µM of each PUFA, after epoxidation by 10 mM mCPBA at 50 ^o^C for an hour, the proportion of the mono-epoxidized product was controlled at 40 % (**Fig. S10 and** **Table S2**). The developed protocol would allow us to target mono-epoxidized products of PUFAs, and quantification of PUFA isomers was thus possible, which would be demonstrated later.

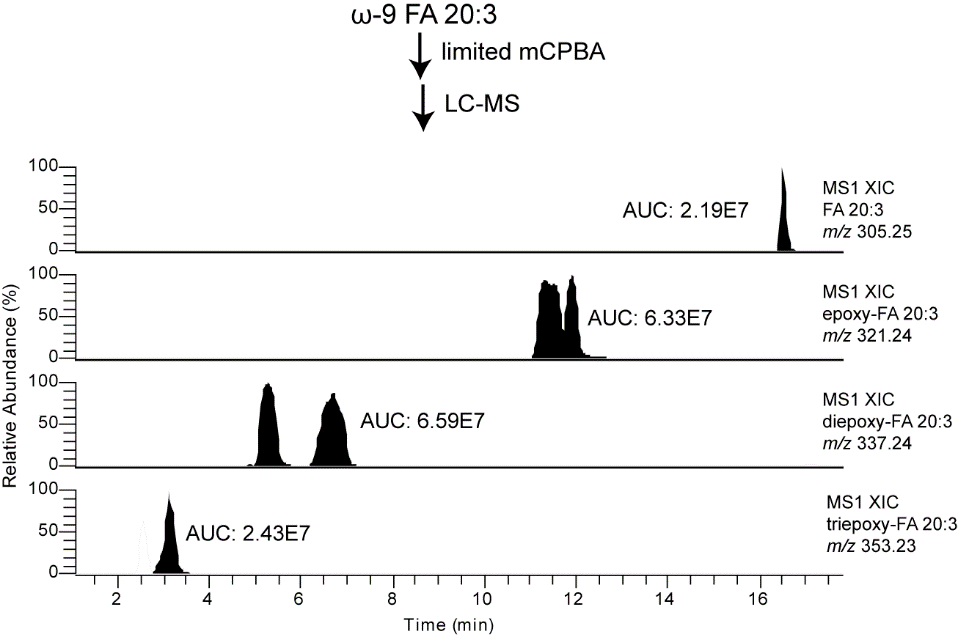

**Figure S10. Estimation of the epoxide yield of PC 16:0/18:1 (9Z) with mCPBA epoxidation.**

The XICs of PC 16:0/18:1(9Z) (100 µM) **(a)** prior to epoxidation and **(b)** after epoxidation using excess mCPBA were shown. The epoxidation yield was estimated by the area under curve ratio (A_epoxy-lipid_ / A_lipid_).

**Table S2. The composition of epoxidation products of PUFAs under the controlled mCPBA epoxidation.**

| **PUFA** | **Epoxide proportion (%)** | | | |
| --- | --- | --- | --- | --- |
|  | **Non-epoxidized lipid** | **epoxy-lipid** | **di-epoxy-lipid** | **Tri-epoxy-lipid** |
| ω-6 FA 18:3 | 6 | 42 | 38 | 13 |
| ω-3 FA 18:3 | 8 | 40 | 35 | 16 |
| ω-9 FA 20:3 | 13 | 32 | 37 | 19 |

**S3. Applications of LC-MS-tPRM in biological lipid extracts**

**S3-1. Development of the mCPBA epoxidation protocols in biological lipid extracts**

**S3-1-1. Monounsaturated lipids**

To ensure the optimal performance of the analysis, two individual derivatization protocols were used for monounsaturated and polyunsaturated lipids respectively, as only mono-epoxidized products were targeted for MS/MS (or MS^3^ for GPL ). For the analysis of monounsaturated lipids, the highest amount of epoxides was ensured by complete epoxidation with excess mCPBA. In the optimized protocol, the biological lipid extract (~5 mg/mL) was added with equal volume of 200 mM mCPBA, accelerated by incubation at 50 ^o^C for at least an hour, and subjected to LC-MS analysis. The representative LC-MS chromatogram of the epoxidized human serum lipid extract was shown in **Fig. S11a**, in which non-epoxidized unsaturated lipids diminished and abundant fully epoxidized unsaturated lipids were highlighted. Using this protocol, the epoxidation could be monitored by the spiked internal standard, D_17­_-FA 18:1 (9Z), resulting an averaged epoxide yield of 83 % (n = 10; SD = 21 %; human serum lipid extract) (**Fig. S11b)**. In addition, the stability of epoxides was validated by analyzing a derivatized serum extract after 1 and 21 hours of epoxidation (stored under room temperature), and no significant degradation of the epoxides was observed (**Figure S11c**).

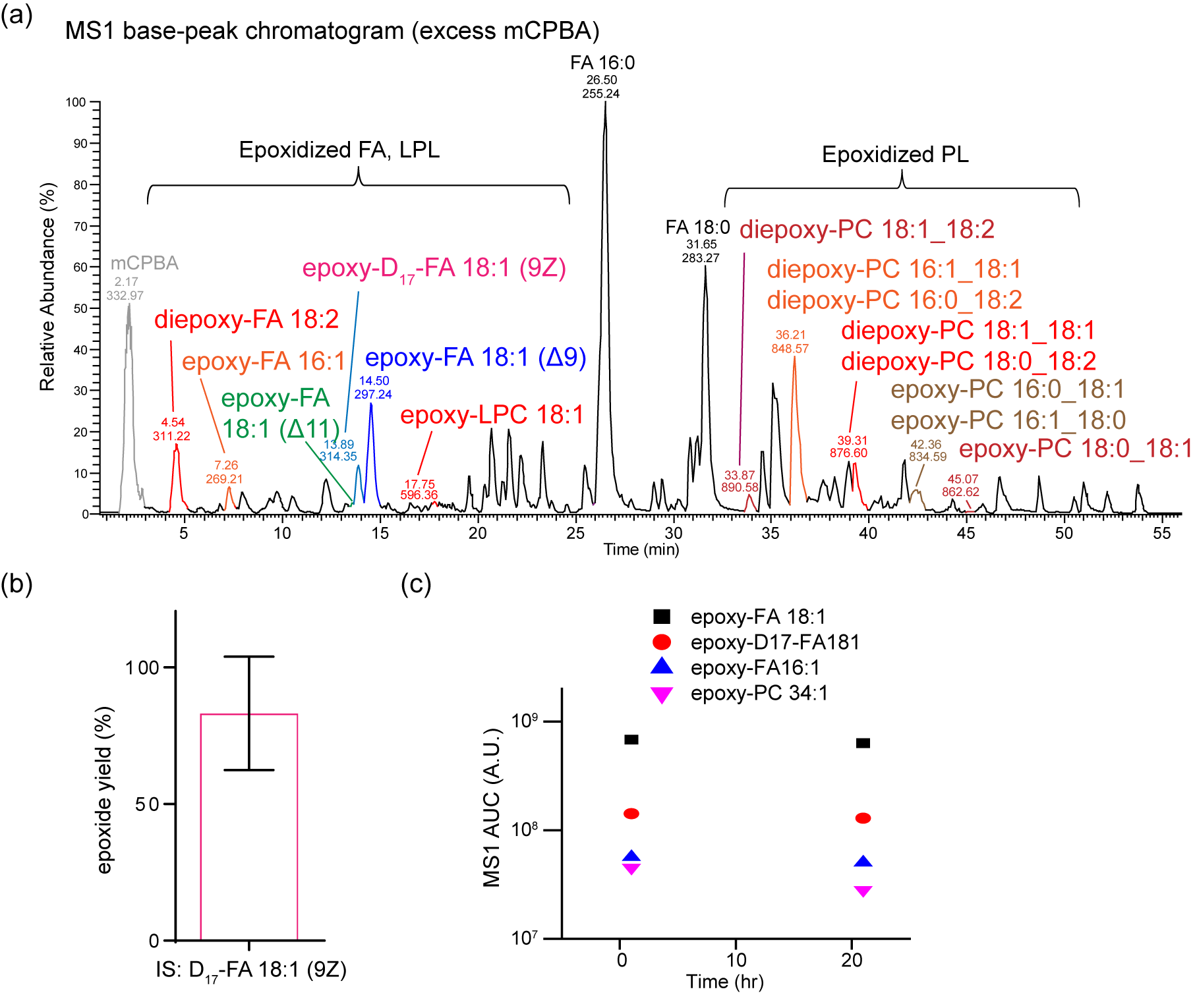

**Figure S11. Evaluation of the epoxidation protocol for the analysis of mono-unsaturated lipids.**

The epoxidation protocol for the anlaysis of mono-unsasturated lipids was evaluated by human serum lipid extracts. **(a)** The representative basepeak chromatogram of a human serum lipid extract after derivatization using excess mCPBA. The epoxidation products were indicated. Peaks were labled by retention time and basepeak *m/z*. **(b)** Estimation of the epoxide yield by the spiked internal standard, D_17­_-FA 18:1 (9Z). The error bar represents SD (n = 10). **(c)** Evaluation of the epoxide stability. The MS1 AUC of four representative epoxidized unsaturated lipids were shown at the time points of 1 hour and 21 hours after derivatization.

**S3-1-2. Polyunsaturated lipids**

The previous protocol was modified for the analysis of polyunsaturated lipids to optimize the amount of mono-epoxidized lipids. In the optimal protocol, the mCPBA solution was diluted into 20 mM; the biological extract was added with equal volume of the diluted mCPBA solution, incubated at 50 ^o^C for at least an hour, and subjected to LC-MS analysis. The representative LC-MS chromatogram was shown in **Fig. S12a**. Importantly, the epoxidation was well-controlled by the limited mCPBA, ensuring the highest proportion of mono-epoxidized species in a given polyunsaturated lipid species (**Fig. S12b-c**).

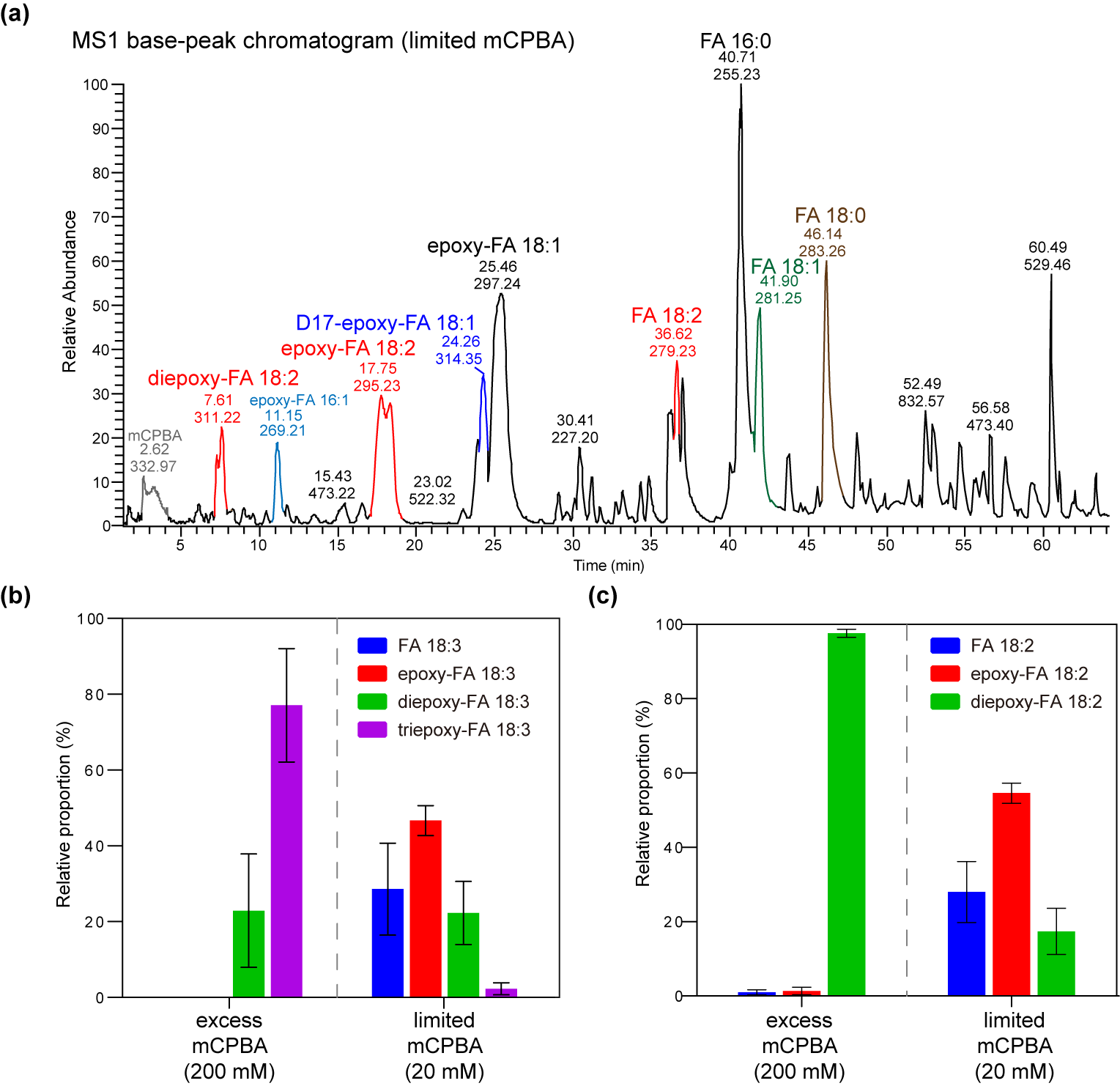

**Figure S12. Evaluation of the epoxidation protocol for poly-unsaturated lipids.**

The epoxidation protocol for the anlaysis of poly-unsasturated lipids was evaluated by human serum lipid extracts. **(a)** The representative basepeak chromatogram of a humns serum lipid extract after derivatization using limited mCPBA. Peaks were labled by retention time and basepeak *m/z*. The epoxide compositions of endogenous polyunsaturated lipids, **(b)** FA 18:3 and **(c)** FA 18:2, using the two derivatization protocols (excess mCPBA for mono-unsaturated lipids, or limited mCPBA for poly-unsaturated lipids). Error bars represent standard deviations (n = 10).

**S3-2. EpoxyFinder: creating the targeted epoxide list for LC-MS-tPRM analysis**

**S3-2-1. Supplementary software: EpoxyFinder**

We developed EpoxyFinder, a graphic-user-interface built using MATLAB algorithms, to assist the transformation of a conventional lipidomics report, which contains a list of identified lipid species including putative structural information of fatty acyl chain composition, e.g. length and number of C=C bonds of each fatty acyl chain, into a LC-MS-tPRM target ion list. The software package, user manual, and the corresponding algorithms are available at:

<https://drive.google.com/drive/folders/13Plb4-aY0NnMpS6pT9UURBMOWCDgkQ-N/>

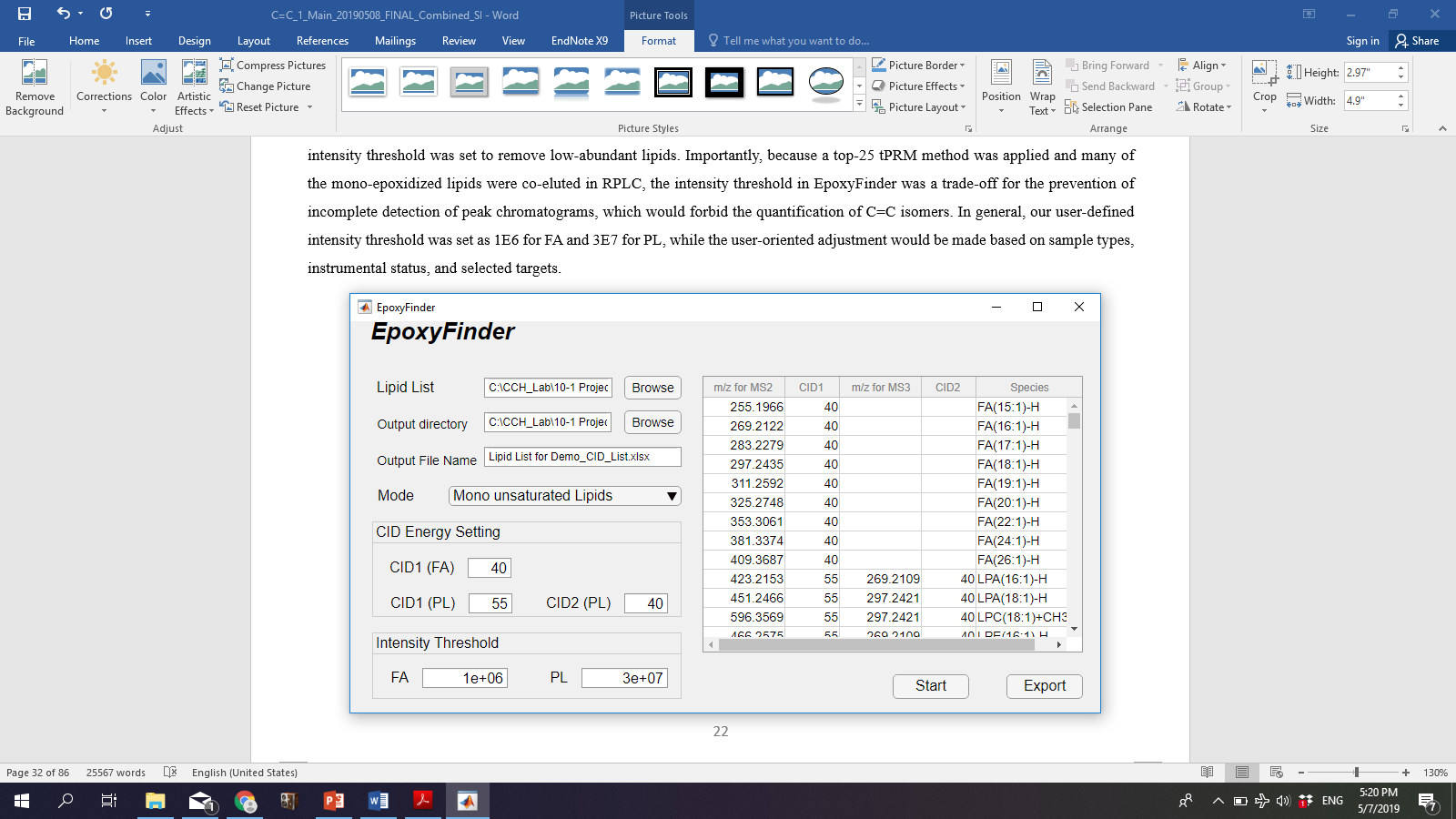

In EpoxyFinder, the report of the identified lipid species (obtained from Thermal LipidSearch software) was loaded and an intensity threshold was set to remove low-abundant lipids. Importantly, because a top-25 tPRM method was applied and many of the mono-epoxidized lipids were co-eluted in RPLC, the intensity threshold in EpoxyFinder was a trade-off for the prevention of incomplete detection of peak chromatograms, which would forbid the quantification of C=C isomers. In general, the user-defined intensity threshold was set at 1E6 for FA and 3E7 for PL in our experiments, while the user-oriented adjustment would be made based on sample types, instrumental status, and selected targets.

**S3-3. The strategy in processing LC-MS-tPRM data**

The general strategy to identify and quantify C=C isomers in biological lipid extracts via LC-MS-tPRM analysis is discussed here. The contents were divided into two parts, as MOFAs and PUFAs were examined separately, and similarly applied to their corresponding GPLs except for that MS^3^ was used to detect the diagnostic ions.

**S3-3-1. MOFA**

1. **High-resolution MS1 identification of parent ions**
   The epoxy-FA parent ion of monoisotopic mass should be detected in the full FT-MS spectrum (with mass error less than 5 ppm), where peaks representing multiple C=C isomers were shown in the XIC.
2. **LC separation of C=C isomer-derived epoxides**
   Isomeric epoxy-lipids of each lipid species were separated in order by RPLC. In specific, the epoxy-lipid was eluted earlier as its C=C position moves toward the methyl end (i.e. larger Δ number, shorter retention time).
3. **XIC confirmation of co-eluted C=C diagnostic ion pairs**
   The corresponding ion-trap (IT) MS/MS spectra for each epoxide peak were inspected carefully. The identification of each isomer was achieved by the detection of a co-eluted C=C diagnostic ion pair.
4. **Quantification by AUC of C=C diagnostic ion pairs**
   Quantification of relative proportions of C=C isomers was estimated by area under curve (AUC) of C=C diagnostic ion pairs in MS2 level. If C=C diagnostic ions were interfered with fragments from the adjacent C=C isomer that resulted in an overlapped XIC of two peaks, the quantification was not accessed.

**S3-3-2. PUFA**

1. **High-resolution MS1 identification of parent ions**
   The parent ion of monoisotopic mass of mono-epoxidized PUFA should be found in the full FT-MS spectrum with mass error less than 5 ppm, and its XIC would show peaks representing multiple isomeric epoxy-PUFAs, which could belong to single or multiple PUFA species.
2. **LC separation of PUFA-derived mono-epoxides**
   Isomeric epoxides of each PUFA species were mostly separated by RPLC, while co-elution might occur in some cases.
3. **LC-MS/MS identification**

As MS/MS of each epoxide could indicate one C=C bond and provide the positional information related to other C=C bonds, identification of a PUFA C=C isomer was achieved by reconstructing the results of multiple mono-epoxidized products. Additionally, the retention time (RT), elution order of epoxides, and MS/MS spectra were further confirmed by those obtained from commercial PUFA standards, if available, in the pre-built LC-MS/MS spectral dataset, as demonstrated in the previous contents (**Main Fig. 3 and Fig. S4-S6**). For uncommon PUFAs that were not listed in our dataset, the C=C position was assigned through carefully interrogating the MS/MS spectra, comparing the elution order with its analogues epoxides, and confirming the RT alignment of the diagnostic ions to each epoxide.

4. **Quantification**
   Due to the complexity of MS/MS spectra of epoxy-PUFA, C=C isomer quantification was only assessed for (i) ω-3 and ω-6 FA 18:3 (ii) ω-6 and ω-9 FA 20:3 (iii) FA 18:3 or FA 20:3 constituting GPLs, as their quantifications were supported by the built calibration curves in **Main Fig. 4**. For other PUFA species with multiple C=C isomers, their quantities were assigned in the order of “major, minor-1, minor-2…” based on the observed MS1 peak heights of the epoxides.

**S4. MELDI-LC-MS-tPRM analysis in human serum lipid extracts**

**S4-1. The LC-MS-tPRM target ion list for human serum lipid extract**

In our current study on C=C isomers in human serum, the target ion list of monounsaturated lipids was provided in **Table S3**, which included 1 internal standard (D_17­_-FA 18:1) and 44 endogenous mono-unsaturated lipid species (7 FA, 4 LPL, and 33 GPL). In addition, the target ion list of polyunsaturated lipids was provided in **Table S4**, which included 50 endogenous poly-unsaturated lipid species (10 FA, 5 LPL, and 35 GPL). Eventually, after removing MS^n^ spectra of low qualities, C=C isomers in 23 targeted monounsaturated lipid species were confidently identified (**Table S5**). and C=C isomers in 36 targeted polyunsaturated lipids species were identified (**Table S6**).

**Table S3. The LC-MS-tPRM target ion list for the analysis of monounsaturated lipids in human serum lipid extract**

For GPLs, the suffix “_2” indicated that the secondary FA chain is targeted, otherwise the first FA chain was targeted. For example, for “PC(17:1/18:1)+CH3COO”, the FA 17:1 chain was targeted in MS3; for “PC(17:1/18:1)+CH3COO _2”, the FA 18:1 chain was targeted.

| **Unsaturated lipid species** | **MS2 epoxide precursor ion (*m/z*)** | **MS3 precursor ion (*m/z*)** |
| --- | --- | --- |
| FA(16:1)-H | 269.21 |  |
| FA(17:1)-H | 283.23 |  |
| FA(18:1)-H | 297.24 |  |
| D17-FA(18:1)-H (IS) | 314.35 |  |
| FA(19:1)-H | 311.26 |  |
| FA(20:1)-H | 325.27 |  |
| FA(22:1)-H | 353.31 |  |
| FA(24:1)-H | 381.34 |  |
| LPC(16:1)+CH3COO | 568.32 | 269.21 |
| LPC(18:1)+CH3COO | 596.36 | 297.24 |
| LPG(18:1)+CH3COO | 525.28 | 297.24 |
| LPI(18:1)+CH3COO | 613.30 | 297.24 |
| PC(16:0/16:1)+CH3COO_2 | 806.55 | 269.21 |
| PC(16:0/18:1)+CH3COO_2 | 834.59 | 297.24 |
| PC(16:1/16:1)+CH3COO | 820.53 | 269.21 |
| PC(16:1/18:1)+CH3COO | 848.57 | 269.21 |
| PC(16:1/18:1)+CH3COO_2 | 848.57 | 297.24 |
| PC(16:1/18:2)+CH3COO | 862.54 | 269.21 |
| PC(17:1/18:1)+CH3COO | 862.58 | 283.23 |
| PC(17:1/18:1)+CH3COO _2 | 862.58 | 297.24 |
| PC(18:0/16:1)+CH3COO _2 | 834.59 | 269.21 |
| PC(18:0/18:1)+CH3COO _2 | 862.62 | 297.24 |
| PC(18:1/18:1)+CH3COO | 876.60 | 297.24 |
| PC(18:1/18:2)+CH3COO | 890.58 | 297.24 |
| PE(16:0/18:1)-H _2 | 732.52 | 297.24 |
| PE(18:0/18:1)-H _2 | 760.55 | 297.24 |
| PE(18:1/18:1)-H | 774.53 | 297.24 |
| PE(18:1/18:2)-H | 788.51 | 297.24 |
| PE(18:1/20:4)-H | 844.50 | 297.24 |
| PE(18:1/20:5)-H | 858.48 | 297.24 |
| PE(20:0/18:1)-H _2 | 788.58 | 297.24 |
| PE(20:1/18:1)-H | 802.56 | 325.27 |
| PE(20:1/18:1)-H _2 | 802.56 | 297.24 |
| PE(21:1/24:6)-H | 970.57 | 339.29 |
| PE(24:5/21:1)-H _2 | 956.59 | 339.29 |
| PE(32:1/20:4)-H | 1040.72 | 493.46 |
| PE(32:1/22:6)-H | 1096.71 | 493.46 |
| PE(34:1/20:4)-H | 1068.75 | 521.49 |
| PI(16:0/18:1)-H _2 | 851.53 | 297.24 |
| PI(18:0/18:1)-H _2 | 879.56 | 297.24 |
| PI(18:1/18:1)-H | 893.54 | 297.24 |
| PI(18:1/18:2)-H | 907.52 | 297.24 |
| PI(18:1/20:4)-H | 963.51 | 297.24 |
| PS(18:0/18:1)-H _2 | 804.54 | 297.24 |
| PS(18:1/24:7)-H | 986.49 | 297.24 |

**Table S4. The LC-MS-tPRM target ion list for the analysis of polyunsaturated lipids in human serum lipid extract**

| **Unsaturated lipid species** | **MS2 epoxide precursor ion (*m/z*)** | **MS3 precursor ion (*m/z*)** |
| --- | --- | --- |
| FA(18:2)-H | 295.23 |  |
| FA(18:3)-H | 293.21 |  |
| FA(18:4)-H | 291.20 |  |
| FA(20:2)-H | 323.26 |  |
| FA(20:3)-H | 321.24 |  |
| FA(20:4)-H | 319.23 |  |
| FA(20:5)-H | 317.21 |  |
| FA(22:4)-H | 347.26 |  |
| FA(22:5)-H | 345.24 |  |
| FA(22:6)-H | 343.23 |  |
| LPC(18:2)+CH3COO | 594.34 | 295.23 |
| LPC(20:4)+CH3COO | 618.34 | 319.23 |
| LPE(18:2)-H | 492.27 | 295.23 |
| LPE(20:4)-H | 516.27 | 319.23 |
| LPE(22:6)-H | 540.27 | 343.23 |
| PC(16:0/18:2)+CH3COO_2 | 832.57 | 295.23 |
| PC(16:0/20:3)+CH3COO_2 | 858.59 | 321.24 |
| PC(16:0/20:4)+CH3COO_2 | 856.57 | 319.23 |
| PC(18:0/18:2)+CH3COO_2 | 860.60 | 295.23 |
| PC(18:1/18:2)+CH3COO_2 | 858.59 | 295.23 |
| PC(18:2/18:2)+CH3COO | 856.57 | 295.23 |
| PC(20:3/20:5)+CH3COO | 904.57 | 321.24 |
| PC(20:3/20:5)+CH3COO_2 | 904.57 | 317.21 |
| PC(20:5/18:2)+CH3COO | 878.56 | 317.21 |
| PC(20:5/18:2)+CH3COO_2 | 878.56 | 295.23 |
| PC(20:5/20:4)+CH3COO | 902.56 | 317.21 |
| PC(20:5/20:4)+CH3COO_2 | 902.56 | 319.23 |
| PE(16:0/18:2)-H_2 | 730.50 | 295.23 |
| PE(16:0/20:4)-H_2 | 754.50 | 319.23 |
| PE(16:0/22:6)-H_2 | 778.50 | 343.23 |
| PE(18:0/18:2)-H_2 | 758.53 | 295.23 |
| PE(18:0/20:4)-H_2 | 782.53 | 319.23 |
| PE(18:0/22:6)-H_2 | 806.53 | 343.23 |
| PE(18:1/18:2)-H_2 | 756.52 | 295.23 |
| PE(18:1/20:5)-H_2 | 778.50 | 317.21 |
| PE(20:2/18:2)-H | 782.53 | 323.26 |
| PE(20:2/18:2)-H_2 | 782.53 | 295.23 |
| PG(19:0/24:7)-H_2 | 877.56 | 369.24 |
| PG(21:2/24:7)-H | 901.56 | 337.27 |
| PG(21:2/24:7)-H_2 | 901.56 | 369.24 |
| PG(25:4/20:4)-H | 903.58 | 389.30 |
| PG(25:4/20:4)-H_2 | 903.58 | 319.23 |
| PI(16:0/18:2)-H_2 | 849.51 | 295.23 |
| PI(16:0/20:4)-H_2 | 873.51 | 319.23 |
| PI(18:0/18:2)-H_2 | 877.54 | 295.23 |
| PI(18:0/18:3)-H_2 | 875.53 | 293.21 |
| PI(18:0/20:3)-H_2 | 903.56 | 321.24 |
| PI(18:0/20:4)-H_2 | 901.54 | 319.23 |
| PI(18:1/18:2)-H_2 | 875.53 | 295.23 |
| PS(18:0/24:6)-H_2 | 878.56 | 371.26 |

**S4-2. Identification and quantification of C=C isomers in human serum**

**S4-2-1. Monounsaturated lipids**

**Table S5. Identification and quantification of monounsaturated C=C isomers in human serum.**

The quantification results are represented by "averaged value ± standard deviation" (n = 10). Relative proportions of C=C isomers were estimated by XIC AUC of the diagnostic ion pairs.

| Lipid class | Lipid species | Molecular ion | MS/MS precursor ion (*m/z*) | MS/MS/MS precursor ion (*m/z*) | Relative proportion (µ ± SD %, n = 10) | | | | | | | | |
| --- | --- | --- | --- | --- | --- | --- | --- | --- | --- | --- | --- | --- | --- |
|  |  |  |  |  | Δ6 | Δ7 | Δ8 | Δ9 | Δ10 | Δ11 | Δ12 | Δ13 | Δ15 |
| FA | FA 16:1 | [M-H]^-^ | 269.21 | --- | 0.5 ± 0.3 | 7.2 ± 2.5 |  | 90.2 ± 2.8 |  | 2.1 ± 0.8 | O (< 1%) |  |  |
|  | FA 17:1 | [M-H]^-^ | 283.23 | --- |  |  | 44.3 ± 10.4 | 39.0 ± 14.6 | 15.7 ± 4.4 |  | 1.1 ± 0.3 |  |  |
|  | FA 18:1 | [M-H]^-^ | 297.24 | --- |  |  |  | 91.8 ± 0.9 |  | 7.8 ± 0.9 |  | 0.3 ± 0.1 |  |
|  | FA 20:1 | [M-H]^-^ | 325.27 | --- |  | 0.5 ± 0.8 |  | 5.8 ± 2.6 |  | 82.2 ± 4.7 |  | 11.1 ± 5.1 | 1.0 ± 0.4 |
|  | FA 22:1 | [M-H]^-^ | 353.31 | --- |  |  |  |  |  | O (major) |  | O (major) | O |
|  | FA 24:1 | [M-H]^-^ | 381.34 | --- |  |  |  |  |  |  |  | O (minor, < 5 %) | O (major, > 95 %) |
| LPLs | LPC 16:1 | [M+CH_3_COO]^-^ | 568.32 | 269.21 |  | 10.4 ± 3.0 |  | 82.7 ± 2.3 |  | 6.9 ± 1.5 |  |  |  |
|  | LPC 18:1* | [M+CH_3_COO]^-^ | 596.36 | 297.24 |  |  |  | 85.7 ± 2.8* |  | 14.3 ± 2.8* |  |  |  |
|  | LPG 18:1 | [M-H]^-^ | 525.28 | 297.24 |  |  |  | 57.9 ± 11.0** |  | 42.1 ± 11.0** |  |  |  |
|  | LPI 18:1 | [M-H]^-^ | 613.30 | 297.24 |  |  |  | O (major) |  | O |  |  |  |
| PLs | PC **16:1**_16:0 | [M+CH_3_COO]^-^ | 806.55 | 269.21 |  | 5.4 ± 3.0 |  | 94.6 ± 3.0 |  |  |  |  |  |
|  | PC **16:1_**18:1 | [M+CH_3_COO]^-^ | 848.57 | 269.21 |  | O |  | O (major) |  |  |  |  |  |
|  | PC **16:1_**18:2 | [M+CH_3_COO]^-^ | 862.54 | 269.21 |  | O |  | O (major) |  | O |  |  |  |
|  | PC **16:1**_18:0 | [M+CH_3_COO]^-^ | 834.59 | 269.21 |  |  |  | O |  |  |  |  |  |
|  | PC **18:1**_16:0 | [M+CH_3_COO]^-^ | 834.59 | 297.24 |  |  |  | 78.6 ± 6.0 |  | 21.4 ± 6.0 |  |  |  |
|  | PC **18:1**_16:1 | [M+CH_3_COO]^-^ | 848.57 | 297.24 |  |  |  | 94.1 ± 4.6 |  | 5.8 ± 4.6 |  |  |  |
|  | PC **18:1**_18:0 | [M+CH_3_COO]^-^ | 862.62 | 297.24 |  |  |  | 82.3 ± 5.2 |  | 17.7 ± 5.2 |  |  |  |
|  | PC **18:1_18:1** | [M+CH_3_COO]^-^ | 876.60 | 297.24 |  |  |  | 90.3 ± 1.9 |  | 9.7 ± 1.9 |  |  |  |
|  | PC **18:1_**18:2 | [M+CH_3_COO]^-^ | 890.58 | 297.24 |  |  |  | 78.0 ± 3.6 |  | 20.7 ± 3.6 |  | 1.3 ± 0.5 |  |
|  | PI **18:1**_16:0 | [M-H]^-^ | 851.53 | 297.24 |  |  |  | O (major) |  | O |  |  |  |
|  | PI **18:1**_18:0 | [M-H]^-^ | 879.56 | 297.24 |  |  |  | O (major) |  | O |  |  |  |
|  | PI **18:1_18:1** | [M-H]^-^ | 893.54 | 297.24 |  |  |  | O (major) |  | O |  |  |  |
|  | PI **18:1_**20:4 | [M-H]^-^ | 963.51 | 297.24 |  |  |  | O (major) |  | O |  |  |  |
| Total lipid species: **23**.  Identified C=C isomers: **58**.  O – quantification was not available due to ion interference from other co-eluted isomers, or insufficient detection of full chromatograms of the diagnostic ions  * - two sets of Δ9 and Δ11 isomers were observed (possibly *sn*-substitutional isomers separated by LC)  ** - estimated by the fully-separated MS1 XIC AUC | | | | | | | | | | | | | |

**S4-2-2. Polyunsaturated lipids**

**Table S6. Identification and quantification of polyunsaturated C=C lipid isomers in human serum.**

The major polyunsaturated C=C isomer in each polyunsaturated lipid species was determined by the most abundant mono-epoxide product, and the minor isomers were listed in order of the mono-epoxide abundance.

| Lipid class | Lipid species | Molecular ion | MS/MS precursor ion (*m/z*) | MS/MS/MS precursor ion (*m/z*) | Major isomer | Minor isomer-1 | Minor isomer-2 |
| --- | --- | --- | --- | --- | --- | --- | --- |
| FA | FA 18:2 | [M-H]^-^ | 295.23 | --- | ω-6 (9, 12) |  |  |
|  | FA 18:3 | [M-H]^-^ | 293.21 | --- | ω-3 (9, 12, 15) | ω-6 (6, 9, 12) |  |
|  | FA 18:4 | [M-H]^-^ | 291.20 | --- | ω-3 (6, 9, 12, 15)^a^ |  |  |
|  | FA 20:2 | [M-H]^-^ | 323.26 | --- | ω-6 (11, 14) | ω-9 (8, 11) |  |
|  | FA 20:3 | [M-H]^-^ | 321.24 | --- | ω-6 (8, 11, 14) | ω-3 (11, 14, 17) | ω-9 (5, 8, 11) |
|  | FA 20:4 | [M-H]^-^ | 319.23 | --- | ω-6 (5, 8, 11, 14) | ω-3 (8, 11, 14, 17)^b^ |  |
|  | FA 20:5 | [M-H]^-^ | 317.21 | --- | ω-3 (5, 8, 11, 14, 17) |  |  |
|  | FA 22:4 | [M-H]^-^ | 347.26 | --- | ω-6 (7, 10, 13, 16) | ω-3 (10, 13, 16, 19)^c^ |  |
|  | FA 22:5 | [M-H]^-^ | 345.24 | --- | ω-3 (7, 10, 13, 16, 19) | ω-6 (4, 7, 10, 13, 16)^d^ |  |
|  | FA 22:6 | [M-H]^-^ | 343.23 | --- | ω-3 (4, 7, 10, 13, 16, 19) |  |  |
| LPLs | LPC 18:2 | [M+CH_3_COO]^-^ | 594.34 | 295.23 | ω-6 (9, 12) | ω-6 (9, 12)* |  |
|  | LPE 18:2 | [M-H]^-^ | 492.27 | 295.23 | ω-6 (9, 12) |  |  |
|  | LPC 20:4 | [M+CH_3_COO]^-^ | 618.34 | 319.23 | ω-6 (5, 8, 11, 14) | ω-6 (5, 8, 11, 14)* |  |
| PLs | PC 16:0_**18:2** | [M+CH_3_COO]^-^ | 832.57 | 295.23 | ω-6 (9, 12) |  |  |
|  | PC 18:0_**18:2** | [M+CH_3_COO]^-^ | 860.60 | 295.23 | ω-6 (9, 12) |  |  |
|  | PC 18:1_**18:2** | [M+CH_3_COO]^-^ | 858.59 | 295.23 | ω-6 (9, 12) |  |  |
|  | PC **18:2**_**18:2** | [M+CH_3_COO]^-^ | 856.57 | 295.23 | ω-6 (9, 12) |  |  |
|  | PC 20:5_**18:2** | [M+CH_3_COO]^-^ | 878.56 | 295.23 | ω-6 (9, 12) |  |  |
|  | PE 16:0_**18:2** | [M-H]^-^ | 730.50 | 295.23 | ω-6 (9, 12) |  |  |
|  | PE 18:0_**18:2** | [M-H]^-^ | 758.53 | 295.23 | ω-6 (9, 12) | ω-6 (9, 12)* |  |
|  | PE 18:1_**18:2** | [M-H]^-^ | 756.52 | 295.23 | ω-6 (9, 12) | ω-6 (9, 12)* |  |
|  | PE 20:2_**18:2** | [M-H]^-^ | 782.53 | 295.23 | ω-6 (9, 12) |  |  |
|  | PI 16:0_**18:2** | [M-H]^-^ | 849.51 | 295.23 | ω-6 (9, 12) |  |  |
|  | PI 18:0_**18:2** | [M-H]^-^ | 877.54 | 295.23 | ω-6 (9, 12) |  |  |
|  | PI 18:1_**18:2** | [M-H]^-^ | 875.53 | 295.23 | ω-6 (9, 12) | ω-6 (9, 12)* |  |
|  | PE **20:2**_18:2 | [M-H]^-^ | 782.53 | 323.26 | ω-6 (11, 14) |  |  |
|  | PC 16:0_**20:3** | [M+CH_3_COO]^-^ | 858.59 | 321.24 | ω-6 (8, 11, 14) |  |  |
|  | PI 18:0_**20:3** | [M-H]^-^ | 903.56 | 321.24 | ω-6 (8, 11, 14) |  |  |
|  | PC 16:0_**20:4** | [M+CH_3_COO]^-^ | 856.57 | 319.23 | ω-6 (5, 8, 11, 14) |  |  |
|  | PC 20:5_**20:4** | [M+CH_3_COO]^-^ | 902.56 | 319.23 | ω-6 (5, 8, 11, 14) |  |  |
|  | PE 16:0_**20:4** | [M-H]^-^ | 754.50 | 319.23 | ω-6 (5, 8, 11, 14) |  |  |
|  | PE 18:0_**20:4** | [M-H]^-^ | 782.53 | 319.23 | ω-6 (5, 8, 11, 14) |  |  |
|  | PI 16:0_**20:4** | [M-H]^-^ | 873.51 | 319.23 | ω-6 (5, 8, 11, 14) |  |  |
|  | PI 18:0_**20:4** | [M-H]^-^ | 901.54 | 319.23 | ω-6 (5, 8, 11, 14) |  |  |
|  | PE 18:0_**22:6** | [M-H]^-^ | 806.53 | 343.23 | ω-3 (4, 7, 10, 13, 16, 19) |  |  |
|  | PE 18:1_**20:5** | [M-H]^-^ | 778.50 | 317.21 | ω-3 (5, 8, 11, 14, 17)^e^ |  |  |
| Total lipid species: **36**.  Identified C=C positional isomers: **48**  * another set of C=C isomers were observed (possibly *sn*-substitutional isomer separated by LC)  a. Δ6 mono-epoxide was not observed  b. Δ11 epoxide and Δ14 epoxide were not observed  c. Δ13 epoxide and Δ16 epoxide were not observed  d. Δ4 epoxide and Δ10 epoxide were not observed  e. only Δ17 epoxide was identified | | | | | | | |

**S4-3. Data processing: identification of C=C isomers in biological lipid extracts**

The details of data processing for quantification of C=C isomers in biological lipid extracts were elaborated here.

Taking FA 16:1 in human serum for example (**Fig. S13**), multiple peaks were found in the full FT-MS XIC of epoxy-FA 16:1 ([M-H]^-^ = *m/z* 269.21), and subsequent analysis of C=C diagnostic ion pairs enabled the identification and quantification of five C=C isomers (**Fig. S13a**). Notably, for each C=C isomer, its C=C diagnostic ions was co-eluted and aligned with epoxy-FA 16:1 peaks in the MS1 XIC. To rule out the possible MS/MS ion interference from co-eluted analytes or other competing fragments in IT-MS/MS, high-resolution LC-FT-MS/MS analysis was subsequently conducted. As a result, the FT-MS/MS spectra of the five epoxides were collected, and the annotations of C=C positions were thus confirmed by the accurate *m/z* values of the diagnostic ions (mass error < 5 ppm) (**Fig. S13c**). Similarly, such the identification and FT-MS/MS validation of C=C isomers were further applied to the other MOFAs, including FA 17:1, 18:1, 20:1, 22:1, and 24:1, and a highly diverse composition of isomers with C=C positions ranging from Δ6 to Δ15 was revealed in human serum (**Fig. S14-S18**). For other identified human serum C=C isomers in GPLs and PUFAs, their MS^n^ spectra as well as XICs of the diagnostic ions had been carefully investigated, and the representative cases were shown in **Fig. S19-21**. Similarly, for the identified C=C isomers in 3T3-L1 adipocyte lipid extract, high-resolution LC-FT-MS/MS was also performed for the validation of C=C isomers in MOFAs (**Fig. S22-S28**). For other identified C=C isomers, representative LC-MS^n^ chromatograms were shown in **Fig. S29-S31**.

Interestingly, we could identify two sets of identical C=C isomers from one unsaturated lipid species, which might be attributed to *sn*-substitutional isomers of GPLs. As shown in **Fig. S21**, for lyso-PC (LPC) 18:2 in human serum, using MELDI-LC-MS-tPRM we found two pairs of Δ9- and Δ12-epoxy-LPC 18:2, which were possibly the epoxidation products of two *sn*-substitutional isomers, sn-1-LPC (OH/18:2) and sn-2-LPC (18:2/OH). In the previous study, Abe et al. showed that sn-1 and sn-2 LPC isomers could be separated by RPLC, in which sn-1-LPC (OH/18:2) had the shorter retention time ^5^. As a result, we proposed that the four epoxy-LPC 18:2 epoxides were derived from two *sn*-substitutional LPC 18:2 isomers. In addition, similar cases could be also found in, for example, most lyso-GPLs in 3T3-L1 adipocytes (**Fig. S29**), and these cases would be remarked in our data reports.

**S4-4. Data processing: quantification of C=C isomer compositions in biological lipid extracts**

Quantification of relative proportions of C=C isomers in a given MOFA or MOFA-constituting GPL species was estimated by AUC of the C=C diagnostic ions. Taking FA 16:1 as an example, five C=C isomers (Δ6, Δ7, Δ9, Δ11, Δ12) were identified, and the quantity of each isomer (A_Δx_) was estimated by AUC of the diagnostic ions (A_m/z_) in the MS/MS channel at *m/z* 261.2 by:

A_Δ6_ = A_m/z 113_ + A_m/z 129_

A_Δ7_ = A_m/z 127_ + A_m/z 143_

A_Δ9_ = A_m/z 155_ + A_m/z 129_

A_Δ11_ = A_m/z 183_ + A_m/z 199_

A_Δ12_ = A_m/z 197_ + A_m/z 203_

The relative proportion of each C=C isomer (Δx %) was thus estimated by:

Δ6 % = A_Δ6_ / (A_Δ6_ + A_Δ7_ + A_Δ9_ + A_Δ11_ + A_Δ12_)

Δ7 % = A_Δ7_ / (A_Δ6_ + A_Δ7_ + A_Δ9_ + A_Δ11_ + A_Δ12_)

Δ9 % = A_Δ9_ / (A_Δ6_ + A_Δ7_ + A_Δ9_ + A_Δ11_ + A_Δ12_)

Δ11 % = A_Δ11_ / (A_Δ6_ + A_Δ7_ + A_Δ9_ + A_Δ11_ + A_Δ12_)

Δ12 % = A_Δ12_ / (A_Δ6_ + A_Δ7_ + A_Δ9_ + A_Δ11_ + A_Δ12_)

Restrictions in quantifying C=C isomers could be: (i) insufficient acquisitions full peak chromatograms of diagnostic ions (ii) the MS/MS ion interference from other epoxides. For (i), one major reason could be instrumental limitation in the scanning speed. Although we had expanded the number of IT-MS^n^ events up to 25 in an acquisition cycle (cycling time was around 6 seconds), this problem still happened in quantifying C=C isomers in some GPLs due to their high similarity in retention time. For (ii), the issue was further elaborated by the quantification of C=C isomers of FA 22:1 in human serum. As shown in **Fig. S17d**, two C=C diagnostic ions, *m/z* 183 (for Δ11) and *m/*z 255 (for Δ15), showed two overlapped peaks due to ion interferences from the Δ13 isomer in IT-MS/MS. Such interfering ions could be differentiated in high-resolution FT-MS/MS (**Fig. S17e**). A similar case could be also found for the quantification of FA 24:1 C=C isomers (**Fig. S18**). However, applying FT-MS/MS events in a top-25 LC-MS-tPRM module could lead to longer cycling time using our Orbitrap Elite Mass Spectrometer, and thus not applicable if tens of unsaturated lipid epoxides were targeted. In the above situations, when the quantification could possibly lead to bias, the relative proportions of C=C isomers were denoted as “major” or “minor” species in the summarized reports.

For quantification of poly-unsaturated lipid C=C isomers, we estimated the C=C isomer compositions in (i) ω-3 and ω-6 FA 18:3 (ii) ω-6 and ω-9 FA 20:3 (iii) FA 18:3 or FA 20:3 constituting GPLs, which were supported by the built calibration curves in **Main Fig. 4**. For other PUFA species with multiple C=C isomers, their quantities were assigned in the order of “major, minor-1, minor-2…” based on the observed MS1 peak heights of the epoxides.

**S4-5. Representative LC-MS-tPRM results of the identified C=C isomers in the human serum**

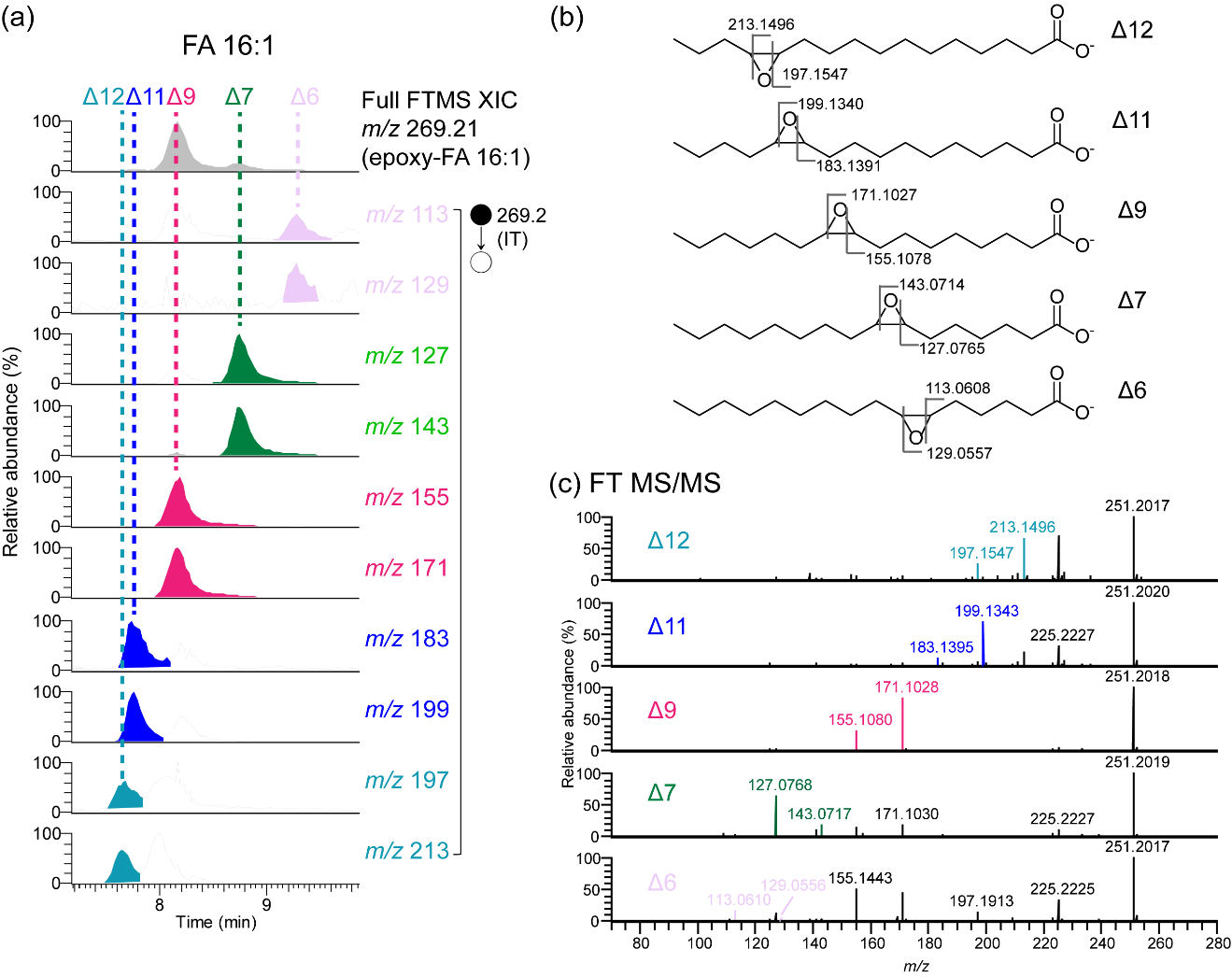

**Figure S13. The identified FA 16:1 C=C isomers in the human serum.**

**(a)** The representative XIC of the epoxidation products of FA 16:1 at FT-MS1 level and the corresponding C=C diagnostic ions in IT-MS2 level. **(b)** The CID scheme of the epoxides, where the theoretical *m/z* values of the proposed fragments were shown. **(c)** The FT-MS/MS spectra of the epoxides.

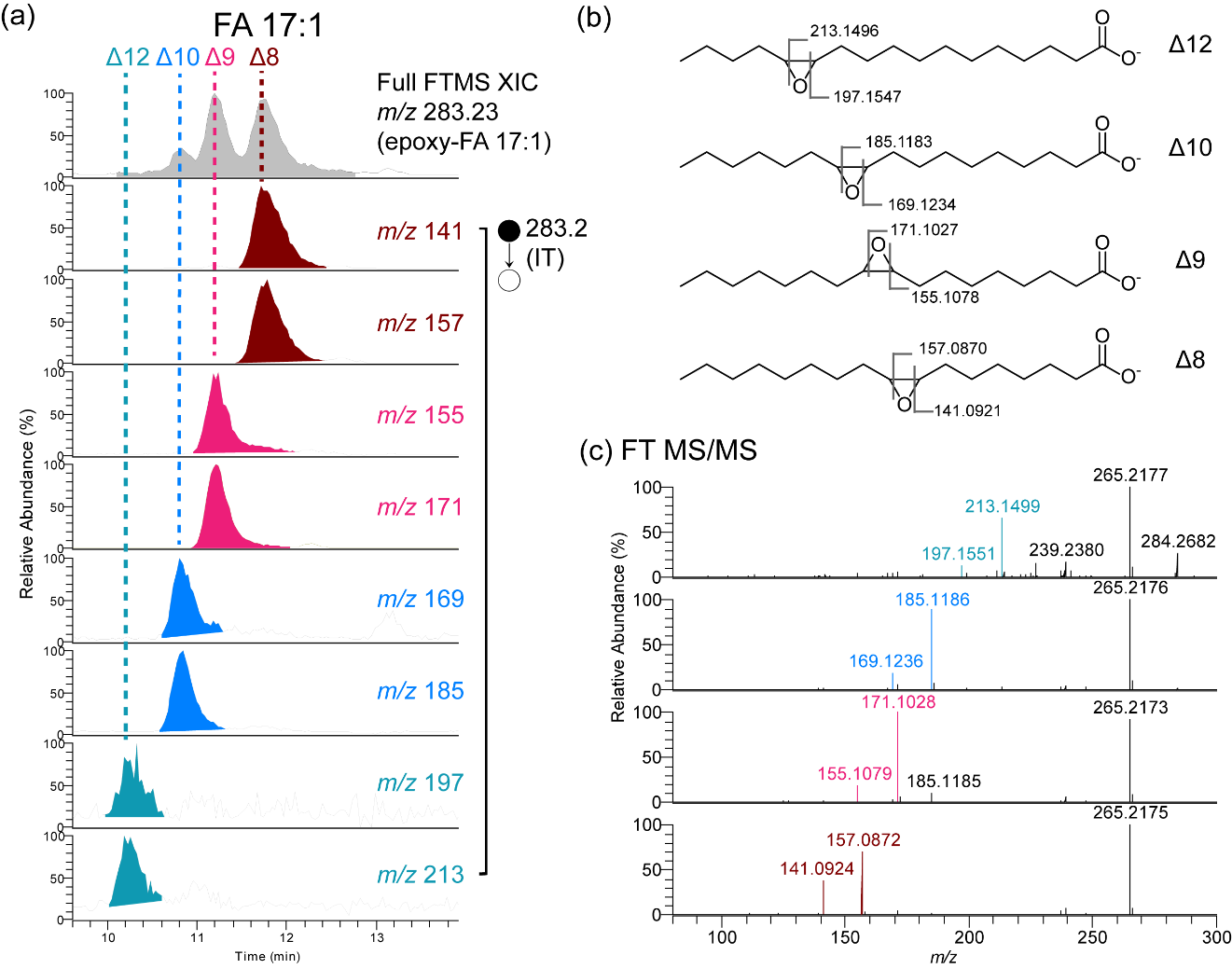

**Figure S14. The identified FA 17:1 C=C isomers in the human serum.**

**(a)** The representative XIC of the epoxidation products of FA 17:1 at FT-MS1 level and the corresponding C=C diagnostic ions in IT-MS2 level. **(b)** The CID scheme of the epoxides, where the theoretical *m/z* values of the proposed fragments were shown. **(c)** The FT-MS/MS spectra of the epoxides.

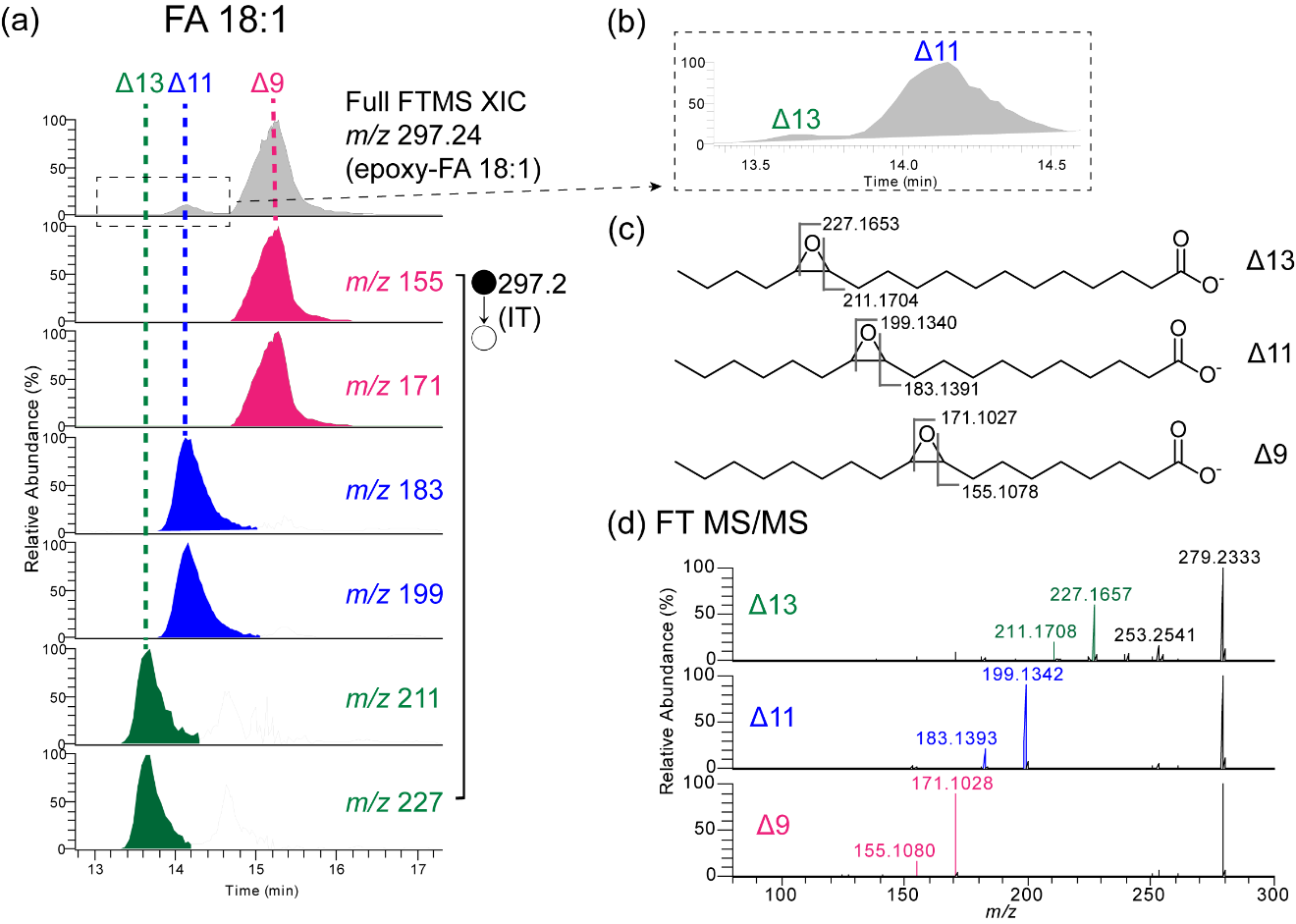

**Figure S15. The identified FA 18:1 C=C isomers in the human serum.**

**(a)** The representative XIC of the epoxidation products of FA 18:1 at FT-MS1 level and the corresponding C=C diagnostic ions in IT-MS2 level. The XIC in retention time during 13.5~14.5 min was highlighted in **(b)**. **(c)** The CID scheme of the epoxides, where the theoretical *m/z* values of the proposed fragments were shown. **(d)** The FT-MS/MS spectra of the epoxides.

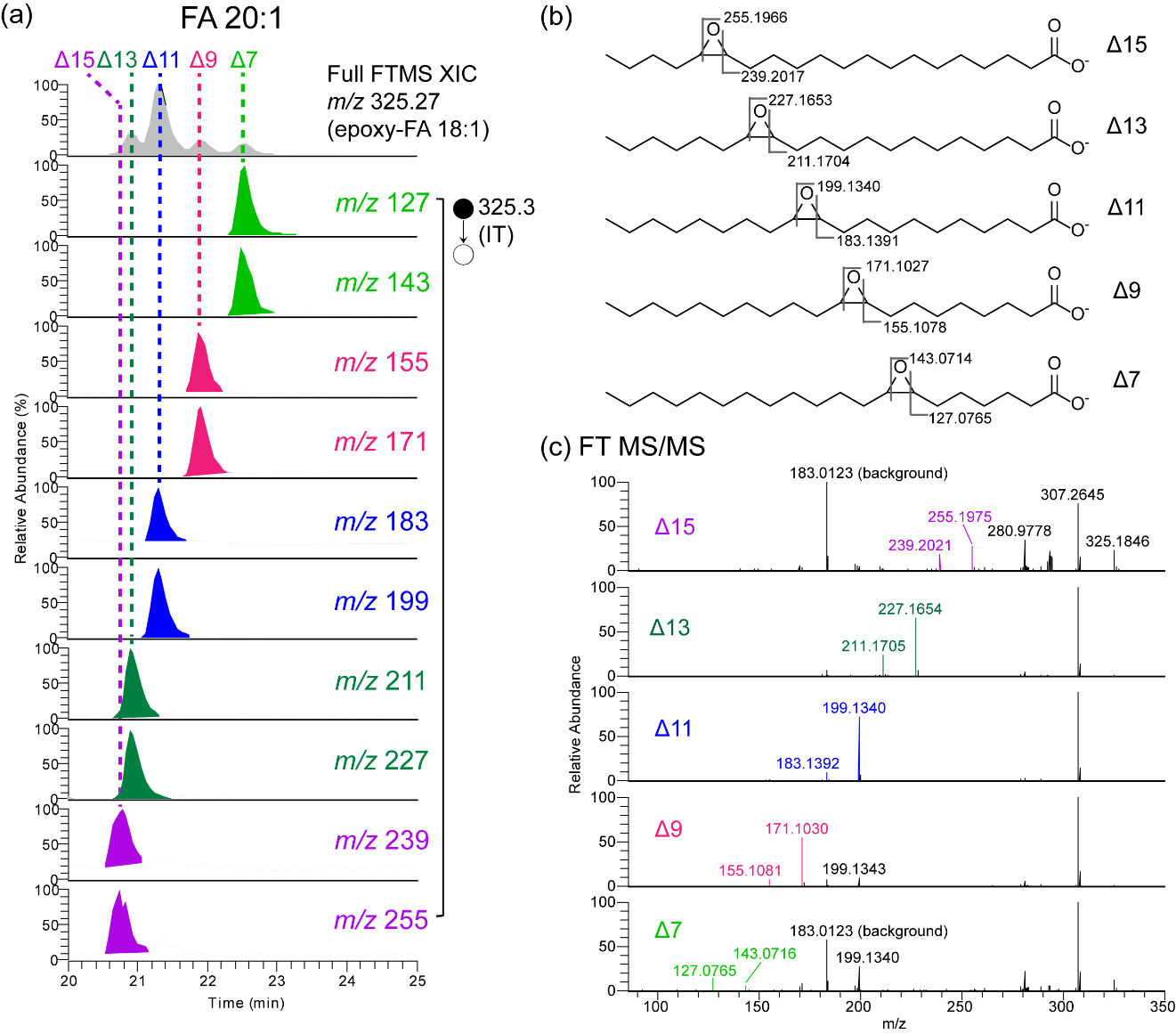

**Figure S16. The identified FA 20:1 C=C isomers in the human serum.**

**(a)** The representative XIC of the epoxidation products of FA 20:1 at FT-MS1 level and the corresponding C=C diagnostic ions in IT-MS2 level. **(b)** The CID scheme of the epoxides, where the theoretical *m/z* values of the proposed fragments were shown. **(c)** The FT-MS/MS spectra of the epoxides.

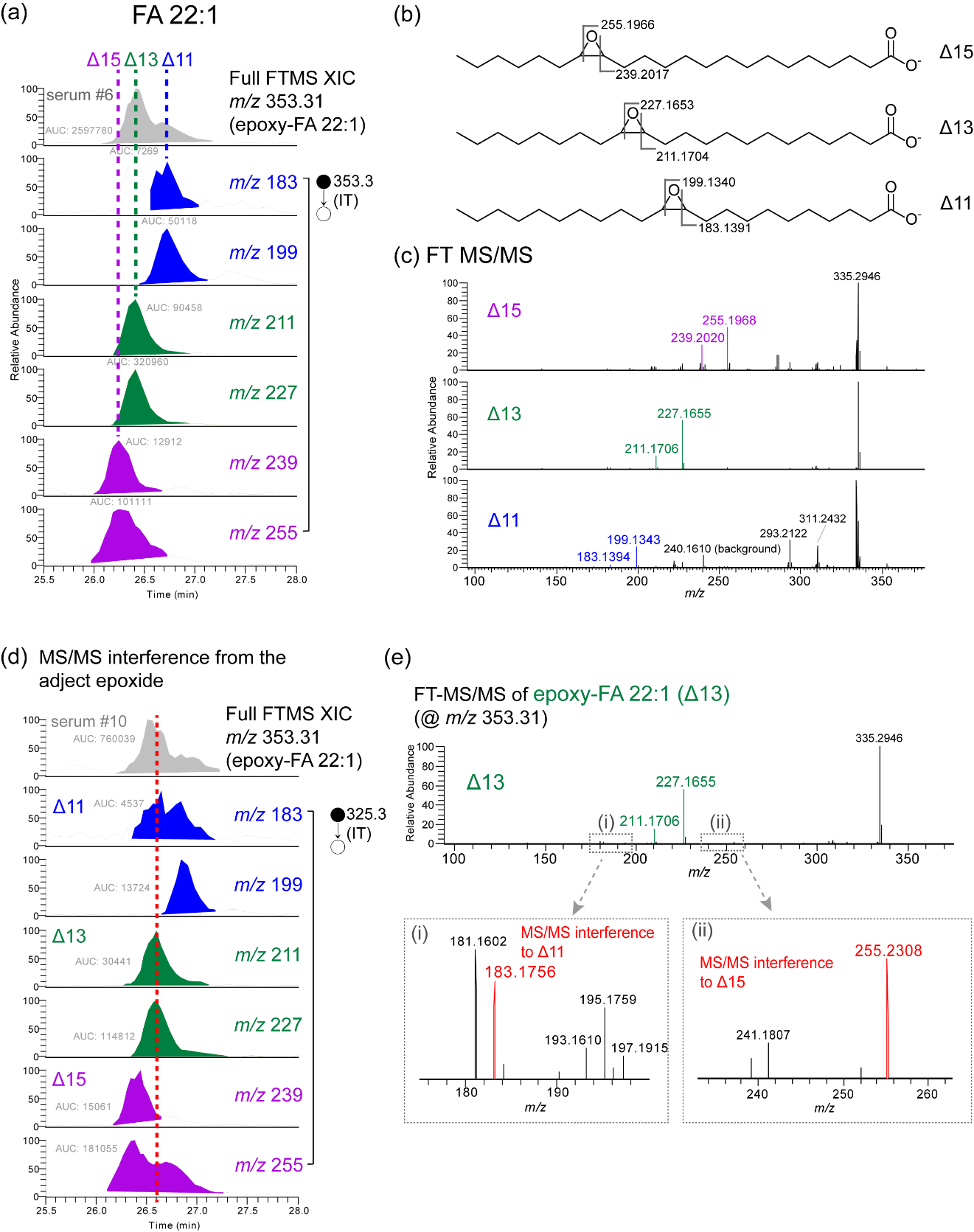

**Figure S17. The identified FA 22:1 C=C isomers in the human serum.**

**(a)** The representative XIC of the epoxidation products of FA 22:1 at FT-MS1 level and the corresponding C=C diagnostic ions in IT-MS2 level. **(b)** The CID scheme of the epoxides, where the theoretical *m/z* values of the proposed fragments were shown. **(c)** The FT-MS/MS spectra of the epoxides. **(d)** The XIC set of another serum sample showed the C=C diagnostic ions of Δ13 in IT-MS/MS were interfered by the adjacent epoxides. **(e)** The MS/MS interfering ions in the FT-MS/MS spectrum of Δ13-epoxy-FA 22:1 were highlighted.

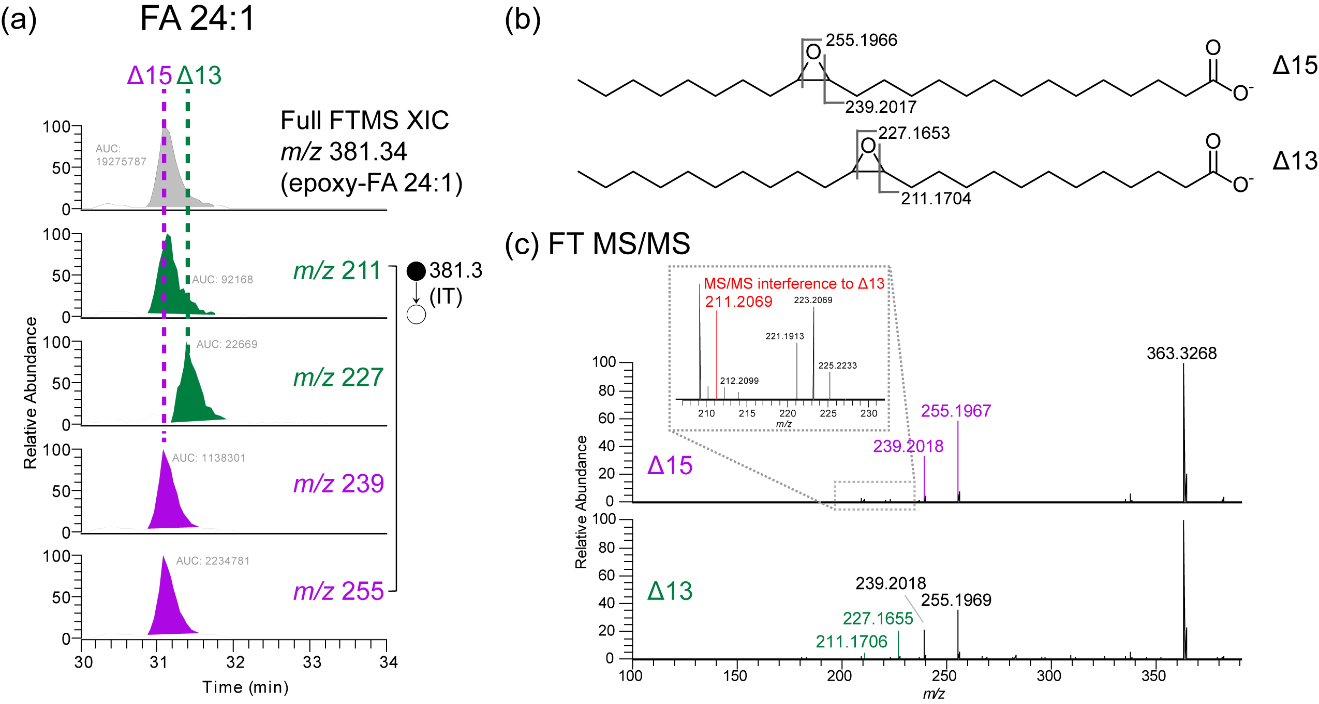

**Figure S18. The identified FA 24:1 C=C isomers in the human serum.**

**(a)** The representative XIC of the epoxidation products of FA 24:1 at FT-MS1 level and the corresponding C=C diagnostic ions in IT-MS2 level. **(b)** The CID scheme of the epoxides, where the theoretical *m/z* values of the proposed fragments were shown. **(c)** The FT-MS/MS spectra of the epoxides. The MS/MS interfering ion was highlighted.

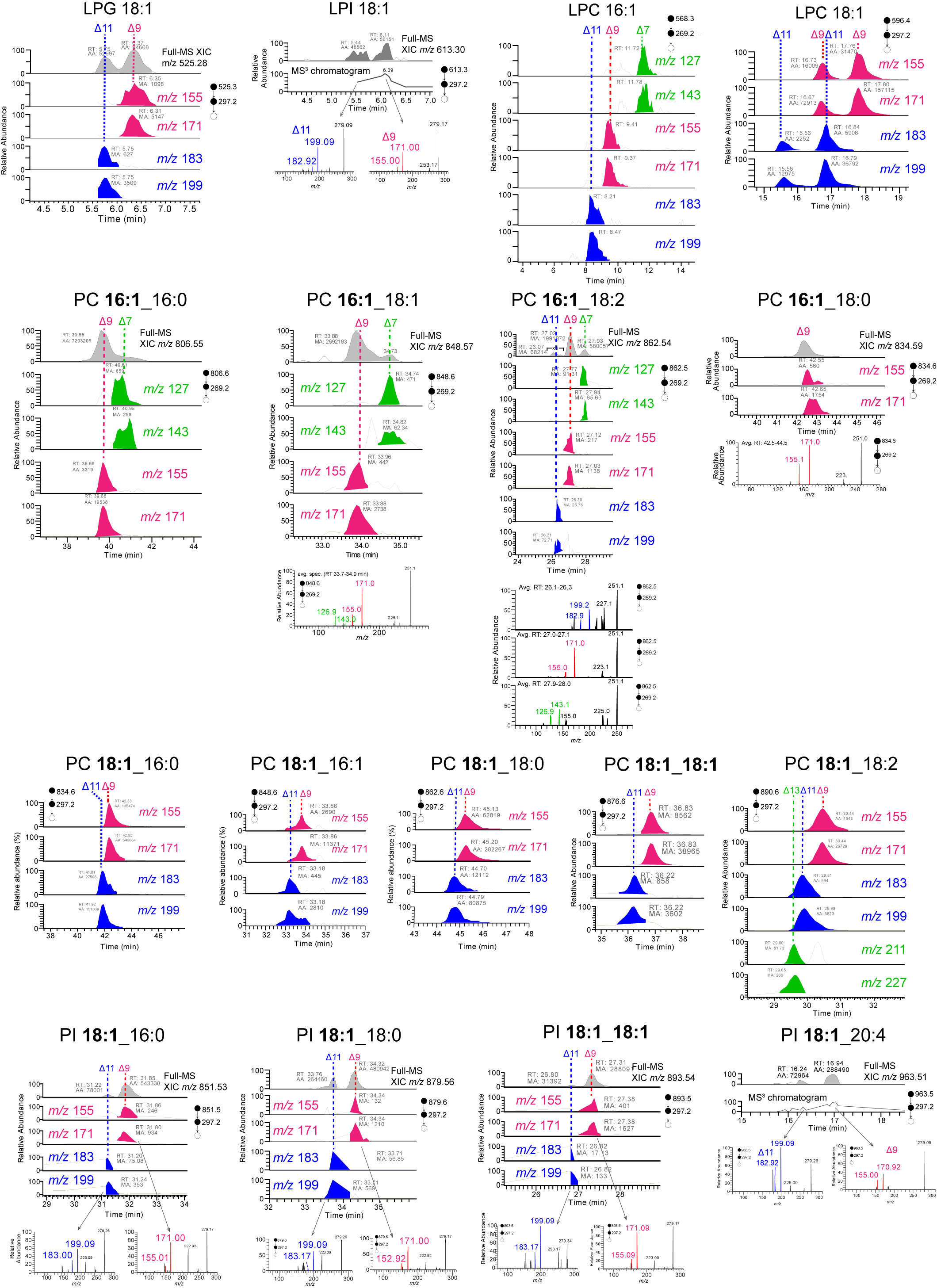

**Figure S19. The identified monounsaturated GPL C=C isomers in the human serum.**

The representative XIC of the epoxidation products of the monounsaturated GPLs at FT-MS1 level and the corresponding C=C diagnostic ions in IT-MS2 level were shown.

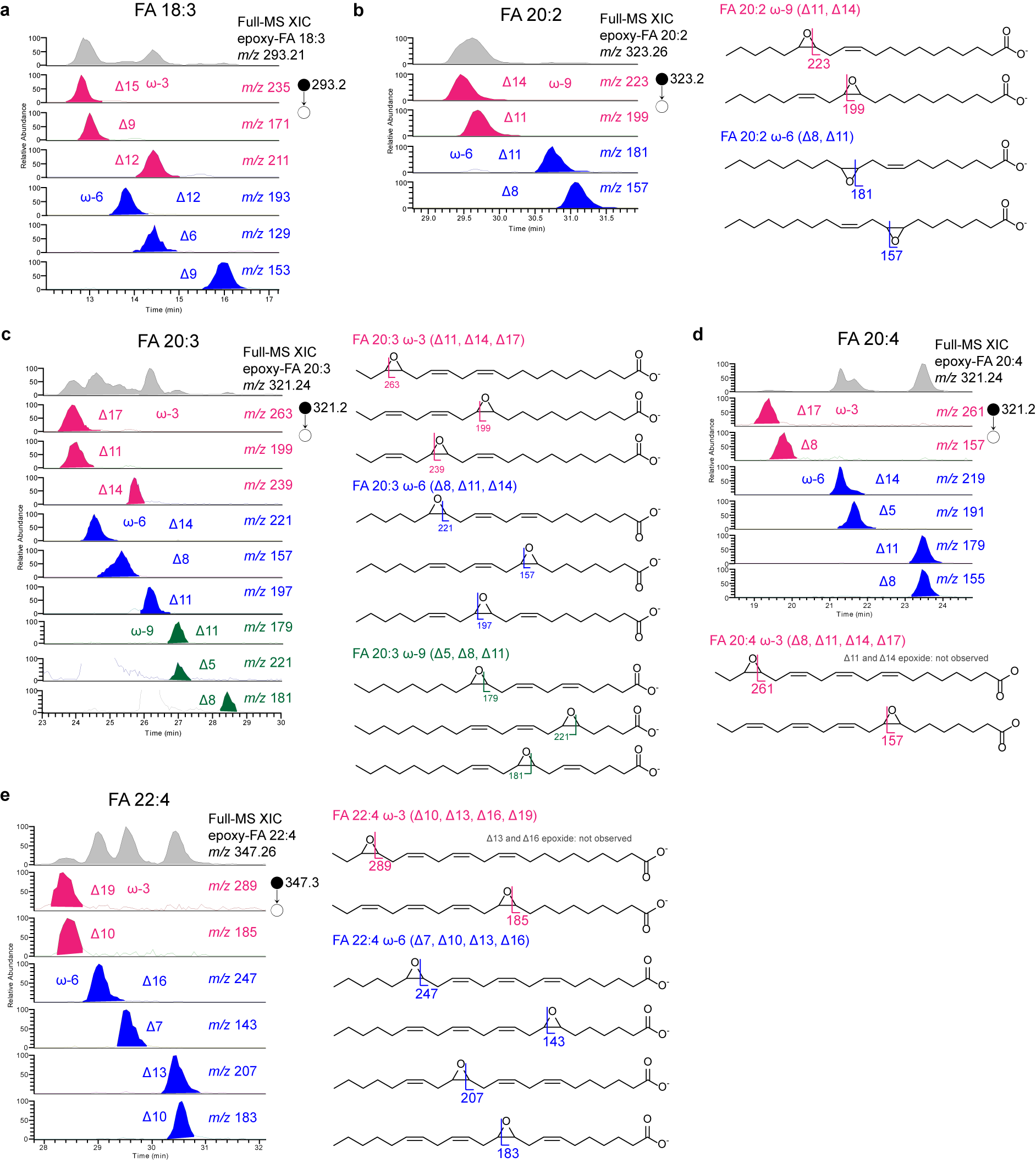

**Figure S20. The identified PUFA C=C isomers in the human serum.**

The representative XIC of the mono-epoxidation products of **(a)** FA 18:3 **(b)** FA 20:2 **(c)** FA 20:3 **(d)** FA 20:4 **(e)** FA 22:4 at FT-MS1 level and the corresponding C=C diagnostic ions in IT-MS2 level. The CID scheme of the epoxides were shown.

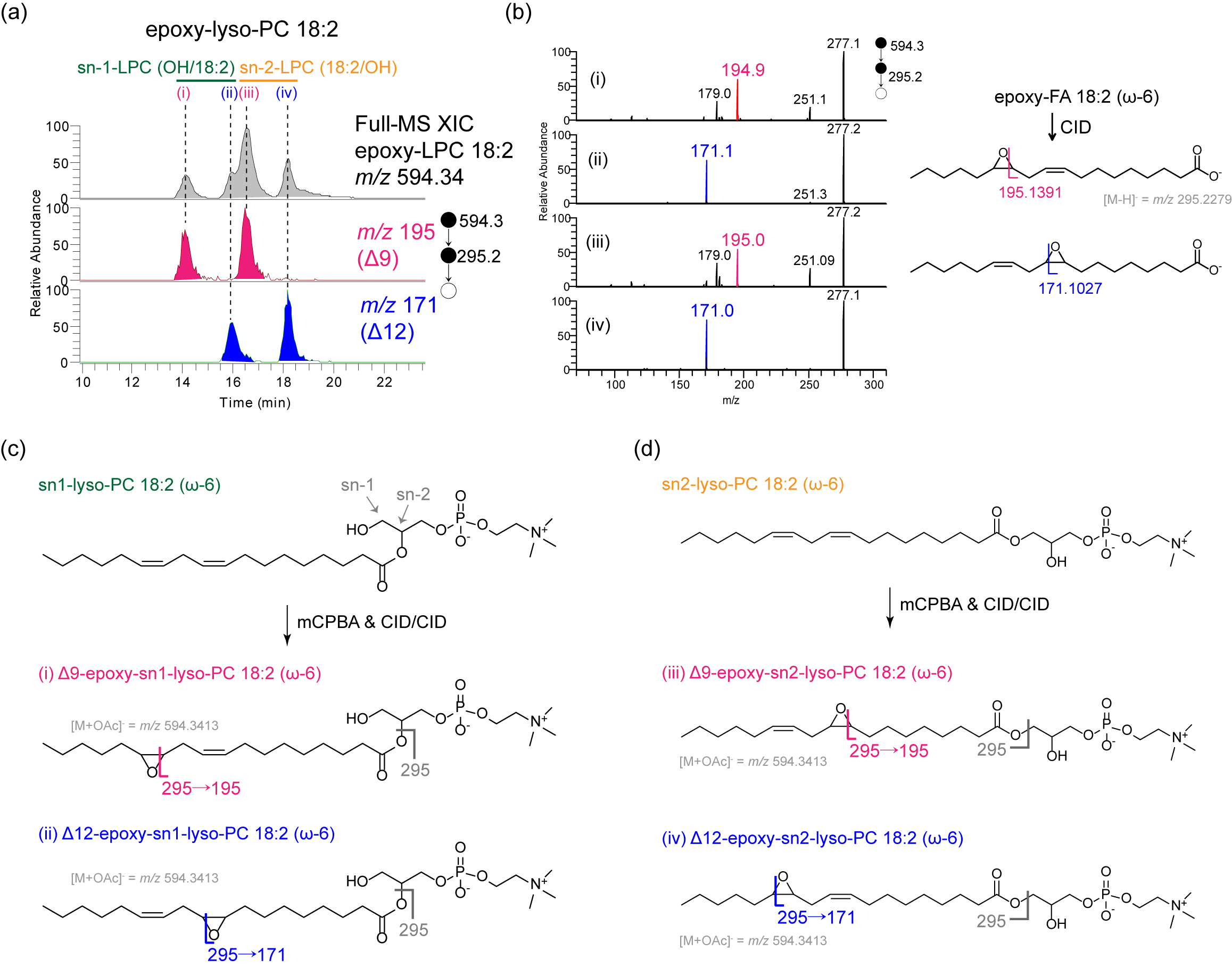

**Figure S21. The possible *sn-*substitutional lyso-PC isomers were found in the human serum.**

**(a)** The representative XIC of the mono-epoxidation products lyso-PC 18:2, where two sets of lyso-PC 18:2 (Δ9, Δ12) were found.

**(b)** The corresponding IT-MS^3^ spectra of the epoxides. The diagnostic ions were indicated. The proposed CID schemes the sn1-LPC 18:2 and sn-2-LPC 18:2 were shown in **(c)** and **(d)**, respectively.

**S4-6. Healthy vs breast cancerous human serum**

**Table S7. Comparison of C=C isomer compositions between healthy human serum and breast cancerous human serum.**

| Lipid species | C=C position | Healthy (n = 4) | | | | | | Breast cancerous (n = 5) | | | | | | | Two-tailed p-value (Student's t test) | Significant difference (*p < 0.05) |
| --- | --- | --- | --- | --- | --- | --- | --- | --- | --- | --- | --- | --- | --- | --- | --- | --- |
|  |  | Sample-1 | Sample-2 | Sample-3 | Sample-4 | Avg. | S.D. | Sample-1 | Sample-2 | Sample-3 | Sample-4 | Sample-5 | Avg. | S.D. |  |  |
| FA 16:1 | Δ6 | 0.5 | 2.7 | 1.1 | 1.3 | 1.4 | 0.9 | 1.3 | 0.8 | 0.4 | 0.4 | 3.1 | 1.2 | 1.1 | 0.818 |  |
|  | Δ7 | 6.3 | 6.4 | 9.2 | 7.1 | 7.2 | 1.3 | 10.6 | 7.0 | 6.6 | 8.8 | 11.7 | 8.9 | 2.2 | 0.224 |  |
|  | Δ9 | 91.3 | 89.0 | 87.3 | 89.0 | 89.2 | 1.6 | 86.0 | 90.4 | 90.7 | 88.8 | 81.9 | 87.6 | 3.7 | 0.445 |  |
|  | Δ11 | 1.8 | 1.9 | 2.4 | 2.6 | 2.2 | 0.4 | 2.0 | 1.9 | 2.2 | 1.9 | 3.3 | 2.3 | 0.6 | 0.796 |  |
| FA 17:1 | Δ8 | 44.0 | 40.4 | 59.7 | 43.2 | 46.8 | 8.7 | 47.3 | 33.6 | 30.3 | 29.6 | 56.2 | 39.4 | 11.8 | 0.328 |  |
|  | Δ9 | 40.7 | 46.0 | 18.8 | 39.2 | 36.2 | 11.9 | 35.6 | 53.5 | 54.9 | 57.4 | 18.5 | 44.0 | 16.7 | 0.458 |  |
|  | Δ10 | 13.7 | 12.1 | 19.6 | 16.0 | 15.4 | 3.3 | 16.0 | 11.1 | 13.0 | 11.1 | 22.5 | 14.7 | 4.8 | 0.829 |  |
|  | Δ12 | 1.6 | 1.4 | 1.9 | 1.5 | 1.6 | 0.2 | 1.1 | 1.9 | 1.7 | 1.9 | 2.8 | 1.9 | 0.6 | 0.417 |  |
| FA 18:1 | Δ9 | 92.5 | 90.6 | 92.0 | 92.2 | 91.8 | 0.8 | 92.1 | 92.0 | 90.6 | 89.8 | 90.9 | 91.1 | 1.0 | 0.254 |  |
|  | Δ11 | 7.1 | 8.8 | 7.5 | 6.6 | 7.5 | 0.9 | 7.5 | 7.6 | 8.9 | 9.8 | 8.2 | 8.4 | 0.9 | 0.214 |  |
|  | Δ13 | 0.3 | 0.5 | 0.5 | 1.2 | 0.6 | 0.4 | 0.4 | 0.4 | 0.6 | 0.4 | 0.9 | 0.5 | 0.2 | 0.589 |  |
| LPC 18:1 | Δ9 | 82.0 | 87.8 | 86.9 | 86.4 | 85.8 | 2.6 | 83.3 | 82.8 | 80.2 | 85.0 | 80.9 | 82.4 | 1.9 | 0.063 |  |
|  | Δ11 | 18.0 | 12.2 | 13.1 | 13.6 | 14.2 | 2.6 | 16.7 | 17.2 | 19.8 | 15.0 | 19.1 | 17.6 | 1.9 | 0.063 |  |
| PC **16:1**_18:2 | Δ9 | 95.3 | 85.7 | 86.7 | 87.5 | 88.8 | 4.4 | 90.4 | 86.1 | 81.3 | 82.3 | 97.6 | 87.5 | 6.7 | 0.755 |  |
|  | Δ11 | 4.7 | 14.3 | 13.3 | 12.5 | 11.2 | 4.4 | 9.6 | 13.9 | 18.7 | 17.7 | 2.4 | 12.5 | 6.7 | 0.755 |  |
| PC **18:1**_16:0 | Δ9 | 80.5 | 80.5 | 80.6 | 81.5 | 80.8 | 0.5 | 80.6 | 82.3 | 73.1 | 71.6 | 77.6 | 77.1 | 4.6 | 0.148 |  |
|  | Δ11 | 19.5 | 19.5 | 19.4 | 18.5 | 19.2 | 0.5 | 19.4 | 17.7 | 26.9 | 28.4 | 22.4 | 22.9 | 4.6 | 0.148 |  |
| PC **18:1**_18:0 | Δ9 | 88.9 | 89.5 | 89.7 | 86.2 | 88.6 | 1.6 | 86.3 | 88.4 | 80.0 | 82.4 | 84.4 | 84.3 | 3.3 | **0.049** | ***** |
|  | Δ11 | 11.1 | 10.5 | 10.3 | 13.8 | 11.4 | 1.6 | 13.7 | 11.6 | 20.0 | 17.6 | 15.6 | 15.7 | 3.3 | **0.049** | ***** |
| PC **18:1**_**18:1** | Δ9 | 95.0 | 96.1 | 95.6 | 96.3 | 95.8 | 0.6 | 92.9 | 93.2 | 93.5 | 93.3 | 87.7 | 92.1 | 2.5 | **0.028** | ***** |
|  | Δ11 | 5.0 | 3.9 | 4.4 | 3.7 | 4.2 | 0.6 | 7.1 | 6.8 | 6.5 | 6.7 | 12.3 | 7.9 | 2.5 | **0.028** | ***** |
| PC **18:1**_18:2 | Δ9 | 80.6 | 74.1 | 82.3 | 75.7 | 78.2 | 3.9 | 81.9 | 85.4 | 75.9 | 86.8 | 84.0 | 82.8 | 4.2 | 0.136 |  |
|  | Δ11 | 19.4 | 25.9 | 17.7 | 24.3 | 21.8 | 3.9 | 18.1 | 14.6 | 24.1 | 13.2 | 16.0 | 17.2 | 4.2 | 0.136 |  |

**S5-1. The LC-MS-tPRM target ion list for 3T3-L1 adipocyte lipid extracts**

In another study on C=C isomers in 3T3-L1 adipocytes, the target ion list of monounsaturated lipids was provided in **Table S5**, which included 1 internal standard (D_17­_-FA 18:1) and 68 endogenous mono-unsaturated lipid species (9 FA, 16 LPL, and 43 GPL). Eventually, C=C isomers in 20 targeted lipid species were confidently identified (**Table S10**). In addition, the target ion list of polyunsaturated lipids was provided in **Table S6**, which included 72 endogenous poly-unsaturated lipid species (14 FA, 10 LPL, and 48 GPL), and finally, C=C isomers in 46 targeted polyunsaturated lipids species were identified (**Table S11**).

**Table S8. The LC-MS-tPRM target ion list for the analysis of monounsaturated lipids in 3T3-L1 adipocyte lipid extract**

| **Unsaturated lipid species** | **MS2 epoxide precursor ion (*m/z*)** | **MS3 precursor ion (*m/z*)** |
| --- | --- | --- |
| FA(16:1)-H | 269.2122 |  |
| FA(17:1)-H | 283.2279 |  |
| FA(18:1)-H | 297.2435 |  |
| D17-FA(18:1)-H (IS) | 314.3483 |  |
| FA(19:1)-H | 311.2592 |  |
| FA(20:1)-H | 325.2748 |  |
| FA(22:1)-H | 353.3061 |  |
| FA(22:2)-H | 367.2854 |  |
| FA(24:1)-H | 381.3374 |  |
| FA(26:1)-H | 409.3687 |  |
| LPA(16:1)-H | 423.2153 | 269.2109 |
| LPA(18:1)-H | 451.2466 | 297.2421 |
| LPC(18:1)+CH3COO | 596.3569 | 297.2421 |
| LPE(16:1)-H | 466.2575 | 269.2109 |
| LPE(17:1)-H | 480.2732 | 283.2265 |
| LPE(18:1)-H | 494.2888 | 297.2421 |
| LPE(19:1)-H | 508.3045 | 311.2577 |
| LPE(22:1)-H | 550.3514 | 353.3045 |
| LPG(16:1)-H | 497.2521 | 269.2109 |
| LPG(18:1)-H | 525.2834 | 297.2421 |
| LPI(16:1)-H | 585.2681 | 269.2109 |
| LPI(18:1)-H | 613.2994 | 297.2421 |
| LPI(19:1)-H | 627.3151 | 311.2577 |
| LPI(20:1)-H | 641.3307 | 325.2733 |
| LPS(16:1)-H | 510.2473 | 269.2109 |
| LPS(18:1)-H | 538.2786 | 297.2421 |
| PA(16:0/18:1)-H_2 | 689.4763 | 297.2421 |
| PA(18:0/16:1)-H_2 | 689.4763 | 269.2109 |
| PA(18:0/18:1)-H_2 | 717.5076 | 297.2421 |
| PA(18:1/18:1)-H | 731.4868 | 297.2421 |
| PA(18:1/18:1)-H_2 | 731.4868 | 297.2421 |
| PA(22:1/20:4)-H | 857.5185 | 353.3045 |
| PE(16:0/18:1)-H_2 | 732.5185 | 297.2421 |
| PE(16:1/18:1)-H | 746.4977 | 269.2109 |
| PE(16:1/18:1)-H_2 | 746.4977 | 297.2421 |
| PE(16:1/20:1)-H | 774.529 | 269.2109 |
| PE(16:1/20:1)-H_2 | 774.529 | 325.2733 |
| PE(17:1/21:3)-H | 830.5188 | 283.2265 |
| PE(18:0/16:1)-H_2 | 732.5185 | 269.2109 |
| PE(18:0/18:1)-H_2 | 760.5498 | 297.2421 |
| PE(18:1/18:1)-H | 774.529 | 297.2421 |
| PE(18:1/18:1)-H_2 | 774.529 | 297.2421 |
| PE(18:1/20:4)-H | 844.4981 | 297.2421 |
| PE(19:1/18:2)-H | 802.5239 | 311.2577 |
| PE(20:1/18:1)-H | 802.5603 | 325.2733 |
| PE(20:1/18:1)-H_2 | 802.5603 | 297.2421 |
| PG(16:0/18:1)-H_2 | 763.5131 | 297.2421 |
| PG(16:2/18:1)-H_2 | 791.4716 | 297.2421 |
| PG(18:0/16:1)-H_2 | 763.5131 | 269.2109 |
| PG(18:0/18:1)-H_2 | 791.5444 | 297.2421 |
| PG(18:1/18:1)-H | 805.5236 | 297.2421 |
| PG(18:1/18:1)-H_2 | 805.5236 | 297.2421 |
| PI(16:0/18:1)-H_2 | 851.5291 | 297.2421 |
| PI(18:0/16:1)-H_2 | 851.5291 | 269.2109 |
| PI(18:0/18:1)-H_2 | 879.5604 | 297.2421 |
| PI(18:1/18:1)-H | 893.5397 | 297.2421 |
| PI(18:1/18:1)-H_2 | 893.5397 | 297.2421 |
| PI(18:1/18:3)-H | 921.4982 | 297.2421 |
| PI(18:1/20:3)-H | 949.5295 | 297.2421 |
| PI(18:1/20:4)-H | 963.5087 | 297.2421 |
| PI(20:1/20:2)-H | 963.5815 | 325.2733 |
| PS(16:0/18:1)-H_2 | 776.5083 | 297.2421 |
| PS(16:1/15:1)-H | 748.4406 | 269.2109 |
| PS(16:1/15:1)-H_2 | 748.4406 | 255.1953 |
| PS(17:0/18:1)-H_2 | 790.524 | 297.2421 |
| PS(18:0/16:1)-H_2 | 776.5083 | 269.2109 |
| PS(18:0/18:1)-H_2 | 804.5396 | 297.2421 |
| PS(18:1/18:1)-H | 818.5189 | 297.2421 |
| PS(18:1/18:1)-H_2 | 818.5189 | 297.2421 |

**Table S9. The LC-MS-tPRM target ion list for the analysis of polyunsaturated lipids in 3T3-L1 adipocyte lipid extract**

| **Unsaturated lipid species** | **MS2 epoxide precursor ion (*m/z*)** | **MS3 precursor ion (*m/z*)** |
| --- | --- | --- |
| FA(18:2)-H | 295.2279 |  |
| FA(18:3)-H | 293.2122 |  |
| FA(20:2)-H | 323.2592 |  |
| FA(20:3)-H | 321.2435 |  |
| FA(20:4)-H | 319.2279 |  |
| FA(20:5)-H | 317.2122 |  |
| FA(22:2)-H | 351.2905 |  |
| FA(22:3)-H | 349.2748 |  |
| FA(22:4)-H | 347.2592 |  |
| FA(22:5)-H | 345.2435 |  |
| FA(22:6)-H | 343.2279 |  |
| FA(24:2)-H | 379.3218 |  |
| FA(24:5)-H | 373.2748 |  |
| FA(24:6)-H | 371.2592 |  |
| LPE(18:2)-H | 492.2732 | 295.2265 |
| LPE(20:2)-H | 520.3045 | 323.2577 |
| LPE(20:3)-H | 518.2888 | 321.2421 |
| LPE(20:4)-H | 516.2732 | 319.2265 |
| LPE(20:5)-H | 514.2575 | 317.2109 |
| LPE(22:4)-H | 544.3045 | 347.2577 |
| LPE(22:5)-H | 542.2888 | 345.2421 |
| LPE(22:6)-H | 540.2732 | 343.2265 |
| LPI(20:3)-H | 637.2994 | 321.2421 |
| LPI(20:4)-H | 635.2838 | 319.2265 |
| PA(18:0/18:2)-H_2 | 715.4919 | 295.2265 |
| PA(20:0/24:6)-H_2 | 819.5545 | 371.2577 |
| PA(22:1/20:4)-H_2 | 793.5389 | 319.2265 |
| PC(18:1/20:5)+CH3COO_2 | 880.5709 | 317.2109 |
| PC(20:3/20:5)+CH3COO | 904.5709 | 321.2421 |
| PC(20:3/20:5)+CH3COO_2 | 904.5709 | 317.2109 |
| PC(20:5/20:4)+CH3COO | 902.5553 | 317.2109 |
| PC(20:5/20:4)+CH3COO_2 | 902.5553 | 319.2265 |
| PC(20:5/20:5)+CH3COO | 900.5396 | 317.2109 |
| PE(16:0/20:4)-H_2 | 754.5028 | 319.2265 |
| PE(17:1/21:3)-H_2 | 782.5341 | 335.2577 |
| PE(18:0/18:2)-H_2 | 758.5341 | 295.2265 |
| PE(18:0/20:4)-H_2 | 782.5341 | 319.2265 |
| PE(18:0/20:5)-H_2 | 780.5185 | 317.2109 |
| PE(18:0/22:4)-H_2 | 810.5654 | 347.2577 |
| PE(18:0/22:6)-H_2 | 806.5341 | 343.2265 |
| PE(18:1/18:2)-H_2 | 756.5185 | 295.2265 |
| PE(18:1/20:4)-H_2 | 780.5185 | 319.2265 |
| PE(18:2/18:2)-H | 754.5028 | 295.2265 |
| PE(19:0/18:3)-H_2 | 770.5341 | 293.2109 |
| PE(19:0/20:4)-H_2 | 796.5498 | 319.2265 |
| PE(19:1/18:2)-H_2 | 770.5341 | 295.2265 |
| PE(20:0/18:2)-H_2 | 786.5654 | 295.2265 |
| PE(20:3/18:2)-H | 780.5185 | 321.2421 |
| PE(20:3/18:2)-H_2 | 780.5185 | 295.2265 |
| PG(16:2/18:1)-H | 759.4818 | 267.1953 |
| PG(18:0/18:2)-H_2 | 789.5287 | 295.2265 |
| PI(16:0/20:4)-H_2 | 873.5135 | 319.2265 |
| PI(16:0/22:4)-H_2 | 901.5448 | 347.2577 |
| PI(18:0/18:2)-H_2 | 877.5448 | 295.2265 |
| PI(18:0/20:2)-H_2 | 905.5761 | 323.2577 |
| PI(18:0/20:3)-H_2 | 903.5604 | 321.2421 |
| PI(18:0/20:4)-H_2 | 901.5448 | 319.2265 |
| PI(18:0/22:4)-H_2 | 929.5761 | 347.2577 |
| PI(18:1/18:3)-H_2 | 873.5135 | 293.2109 |
| PI(18:1/20:3)-H_2 | 901.5448 | 321.2421 |
| PI(18:1/20:4)-H_2 | 899.5291 | 319.2265 |
| PI(20:1/20:2)-H_2 | 931.5917 | 323.2577 |
| PS(18:0/18:2)-H_2 | 802.524 | 295.2265 |
| PS(18:0/20:4)-H_2 | 826.524 | 319.2265 |
| PS(18:0/22:5)-H_2 | 852.5396 | 345.2421 |
| PS(18:0/22:6)-H_2 | 850.524 | 343.2265 |
| PS(18:0/24:6)-H_2 | 878.5553 | 371.2577 |
| PS(18:1/22:4)-H_2 | 852.5396 | 347.2577 |
| PS(24:5/16:0)-H | 852.5396 | 373.2733 |
| PS(24:5/18:0)-H | 880.5709 | 373.2733 |
| PS(24:5/20:4)-H | 900.5396 | 373.2733 |
| PS(24:5/20:4)-H_2 | 900.5396 | 319.2265 |

**S5-2. Identification and quantification of C=C isomers in 3T3-L1 adipocytes**

**S5-2-1. Monounsaturated lipids**

**Table S10. Identification and quantification of C=C isomers in the differentiated 3T3-L1 adipocytes.**

The quantification results are represented by "averaged value ± standard deviation" (n = 4).

| Lipid class | Lipid species | MS/MS precursor ion (*m/z*) | MS/MS/MS precursor ion (*m/z*) | Relative proportion (%) | | | | | | | | | | |
| --- | --- | --- | --- | --- | --- | --- | --- | --- | --- | --- | --- | --- | --- | --- |
|  |  |  |  | Δ6 | Δ7 | Δ8 | Δ9 | Δ10 | Δ11 | Δ12 | Δ13 | Δ14 | Δ15 | Δ17 |
| FA | FA 16:1 | 269.21 | --- | 4.5 ± 1.6 | 18.5 ± 1.4 |  | 68.0 ± 6.3 |  | 3.2 ± 1.0 | 5.8 ± 5.0 |  |  |  |  |
|  | FA 17:1 | 283.23 | --- | 1.0 ± 0.2 | 1.6 ± 0.1 | 23.2 ± 6.5 | 54.8 ± 12.0 | 13.9 ± 5.9 | 2.9 ± 0.4 | 2.7 ± 0.3 |  |  |  |  |
|  | FA 18:1 | 297.24 | --- |  |  | 2.2 ± 0.1 | 69.7 ± 1.6 |  | 25.8 ± 1.5 |  | 1.7 ± 0.1 | 0.6 ± 0.1 |  |  |
|  | FA 19:1 | 311.26 | --- |  |  | 7.4 ± 0.9 | 9.9 ± 1.6 |  | 79.4 ± 2.8 |  | 3.4 ± 0.6 |  |  |  |
|  | FA 20:1 | 325.27 | --- |  | 3.4 ± 0.2 | 3.5 ± 0.2 | 1.1 ± 0.1 |  | 60.6 ± 1.9 |  | 31.4 ± 2.3 |  |  |  |
|  | FA 22:1 | 353.31 | --- |  |  |  | 8.4 ± 0.5 | 2.7 ± 0.1 |  |  | 75.1 ± 0.3 |  | 13.8 ± 0.9 |  |
|  | FA 24:1 | 381.34 | --- |  |  |  | 1.2 ± 0.1 |  | 1.0 ± 0.1 | 0.6 ± 0.04 |  |  | 86.8 ± 1.0 | 10.4 ± 0.9 |
| LPLs | LPE 18:1^a^ | 494.29 | 297.24 |  |  | 1.1 ± 0.1 | 92.0 ± 1.8 |  | 6.7 ± 1.7 |  | 0.12 ± 0.02 |  |  |  |
|  | LPG 18:1^a^ | 525.28 | 297.24 |  |  |  | 56.7 ± 2.6 |  | 43.3 ± 2.6 |  |  |  |  |  |
|  | LPI 18:1 | 613.30 | 297.24 |  |  | 2.0 ± 0.2 | 64.1 ± 2.4 |  | 32.6 ± 2.5 |  | 1.2 ± 0.2 |  |  |  |
|  | LPI 20:1 | 641.33 | 325.27 |  |  |  |  |  | 78.4 ± 5.7 |  | 21.6 ± 5.7 |  |  |  |
| PLs | PG **18:1**_16:0 | 763.51 | 297.24 |  |  |  | 24.4 ± 2.0 |  | 75.3 ± 2.0 |  | 0.24 ± 0.01 |  |  |  |
|  | PG **18:1_18:1** | 805.52 | 297.24 |  |  |  | 71.0 ± 1.5 |  | 28.5 ± 1.5 |  | 0.6 ± 0.2 |  |  |  |
|  | PI **16:1_**18:0 | 851.53 | 269.21 |  | 64.2 ± 2.5 |  | 35.8 ± 2.5 |  |  |  |  |  |  |  |
|  | PI **18:1_**16:0 | 851.53 | 297.24 |  |  |  | 79.6 ± 2.1 |  | 20.4 ± 2.1 |  |  |  |  |  |
|  | PI **18:1**_20:4 | 963.51 | 297.24 |  |  | 1.5 ± 0.2 | 68.6 ± 1.3 |  | 28.6 ± 0.9 |  | 1.3 ± 0.3 |  |  |  |
|  | PE **18:1**_**18:1** | 774.53 | 297.24 |  |  |  | 84.1 ± 0.7 |  | 15.4 ± 0.7 |  | 0.4 ± 0.1 |  |  |  |
|  | PE **16:1**_18:0 | 732.52 | 269.21 | 3.9 ± 0.6 | 13.1 ± 1.1 |  | 80.6 ± 1.8 |  | 2.5 ± 0.3 |  |  |  |  |  |
|  | PE **18:1**_16:0 | 732.52 | 297.24 |  |  |  | 86.6 ± 0.7 |  | 13.4 ± 0.7 |  |  |  |  |  |
|  | PS **18:1**_18:0 | 804.54 | 297.24 |  |  |  | 83.5 ± 2.4 |  | 16.5 ± 2.4 |  |  |  |  |  |
| Total species: **20**  Identified C=C isomers: **78**  a. two sets of the Δ8, Δ9, Δ11, and Δ13 isomers were observed (possibly *sn*-substitutional isomers separated by LC) | | | | | | | | | | | | | | |

**S5-2-2. Polyunsaturated lipids**

**Table S11. - Identification and quantification of polyunsaturated C=C lipid isomers in the differentiated 3T3-L1 adipocytes.**

The major polyunsaturated C=C isomers in each polyunsaturated lipid species were determined by the most abundant mono-epoxide products, and the minor isomers were listed in order of the mono-epoxide.

| Lipid class | Lipid species | Molecular ion | MS/MS precursor ion (*m/z*) | MS/MS/MS precursor ion (*m/z*) | Major isomer | Minor isomer-1 | Minor isomer-2 | Minor isomer-3 | Minor isomer-4 | Minor isomer-5 |
| --- | --- | --- | --- | --- | --- | --- | --- | --- | --- | --- |
| FA | FA 18:2 | [M-H]^-^ | 295.228 | --- | ω-6 (9, 12) | ω-7 (8,11) | ω-9 (6, 9) |  |  |  |
|  | FA 18:3 | [M-H]^-^ | 293.212 | --- | ω-6 (6, 9, 12) | ω-7 (5, 8, 11) | ω-3 (9, 12, 15)^a^ |  |  |  |
|  | FA 20:2 | [M-H]^-^ | 323.259 | --- | ω-9 (8, 11) | ω-6 (11, 14) | ω-7 (10, 13) | conj. ω-6 (12, 14) | conj. ω-7 (11, 13) |  |
|  | FA 20:3 | [M-H]^-^ | 321.244 | --- | ω-6 (8, 11, 14) | ω-9 (5, 8, 11) |  |  |  |  |
|  | FA 20:4 | [M-H]^-^ | 319.228 | --- | ω-6 (5, 8, 11, 14) |  |  |  |  |  |
|  | FA 20:5 | [M-H]^-^ | 317.212 | --- | ω-3 (5, 8, 11, 14, 17) |  |  |  |  |  |
|  | FA 22:2 | [M-H]^-^ | 351.290 | --- | ω-9 (10, 13) | ω-6 (13, 16) | ω-7 (12, 15) | ω-10 (9, 12) |  |  |
|  | FA 22:3 | [M-H]^-^ | 349.275 | --- | ω-9 (7 ,10, 13) | ω-6 (10 ,13, 16) | ω-7 (9, 12, 15) |  |  |  |
|  | FA 22:4 | [M-H]^-^ | 347.259 | --- | ω-6 (7, 10, 13, 16) |  |  |  |  |  |
|  | FA 22:5 | [M-H]^-^ | 345.244 | --- | ω-3 (7, 10, 13, 16, 19) | ω-6 (4, 7, 10, 13, 16)^b^ |  |  |  |  |
|  | FA 22:6 | [M-H]^-^ | 343.228 | --- | ω-3 (4, 7, 10, 13, 16, 19) |  |  |  |  |  |
|  | FA 24:2 | [M-H]^-^ | 379.322 | --- | ω-6 (15, 18) | ω-9 (12, 15) |  |  |  |  |
|  | FA 24:6 | [M-H]^-^ | 371.259 | --- | ω-3 (6, 9, 12, 15, 18, 21) |  |  |  |  |  |
| LPLs | LPE 18:2 | [M-H]^-^ | 492.273 | 295.227 | ω-6 (9, 12) | ω-6 (9, 12)* | ω-7 (8,11) | ω-7 (8,11)* |  |  |
|  | LPE 20:2 | [M-H]^-^ | 520.304 | 323.258 | ω-9 (8, 11) | ω-9 (8, 11)* | ω-6 (11, 14) | ω-6 (11, 14)* | ω-7 (10, 13) | ω-10 (7, 10) |
|  | LPE 20:3 | [M-H]^-^ | 518.289 | 321.242 | ω-9 (5, 8, 11) | ω-9 (5, 8, 11)* | ω-6 (8, 11, 14) | ω-6 (8, 11, 14)* |  |  |
|  | LPE 20:4 | [M-H]^-^ | 516.273 | 319.227 | ω-6 (5, 8, 11, 14) | ω-6 (5, 8, 11, 14)* |  |  |  |  |
|  | LPE 20:5 | [M-H]^-^ | 514.258 | 317.211 | ω-3 (5, 8, 11, 14, 17) | ω-3 (5, 8, 11, 14, 17)^c^ |  |  |  |  |
|  | LPE 22:4 | [M-H]^-^ | 544.304 | 347.258 | ω-6 (7, 10, 13, 16) | ω-6 (7, 10, 13, 16)* |  |  |  |  |
|  | LPE 22:5 | [M-H]^-^ | 542.289 | 345.242 | ω-3 (7, 10, 13, 16, 19) | ω-3 (7, 10, 13, 16, 19)* |  |  |  |  |
|  | LPE 22:6 | [M-H]^-^ | 540.273 | 343.227 | ω-3 (4, 7, 10, 13, 16, 19) |  |  |  |  |  |
|  | LPI 20:3 | [M-H]^-^ | 637.299 | 321.242 | ω-9 (5, 8, 11) | ω-9 (5, 8, 11)* | ω-6 (8, 11, 14) | ω-6 (8, 11, 14)* |  |  |
|  | LPI 20:4 | [M-H]^-^ | 635.284 | 319.227 | ω-6 (5, 8, 11, 14) | ω-6 (5, 8, 11, 14)* |  |  |  |  |
| PLs | PA 18:0_**18:2** | [M-H]^-^ | 715.492 | 295.227 | ω-6 (9, 12) |  |  |  |  |  |
|  | PE 16:0_**20:4** | [M-H]^-^ | 754.503 | 319.227 | ω-6 (5, 8, 11, 14) |  |  |  |  |  |
|  | PE 18:0_**18:2** | [M-H]^-^ | 758.534 | 295.227 | ω-6 (9, 12) | ω-6 (9, 12)* | ω-7 (8,11) |  |  |  |
|  | PE 18:0_**20:4** | [M-H]^-^ | 782.534 | 319.227 | ω-6 (5, 8, 11, 14) | ω-6 (5, 8, 11, 14)* |  |  |  |  |
|  | PE 18:0_**20:5** | [M-H]^-^ | 780.518 | 317.211 | ω-3 (5, 8, 11, 14, 17) | ω-3 (5, 8, 11, 14, 17)^d^ |  |  |  |  |
|  | PE 18:0_**22:4** | [M-H]^-^ | 810.565 | 347.258 | ω-6 (7, 10, 13, 16) | ω-6 (7, 10, 13, 16)* |  |  |  |  |
|  | PE 18:0_**22:6** | [M-H]^-^ | 806.534 | 343.227 | ω-3 (4, 7, 10, 13, 16, 19) |  |  |  |  |  |
|  | PE 18:1_**18:2** | [M-H]^-^ | 756.518 | 295.227 | ω-6 (9, 12) | ω-7 (8,11) |  |  |  |  |
|  | PE **18:2**_**18:2** | [M-H]^-^ | 754.503 | 295.227 | ω-6 (9, 12) | ω-7 (8,11) |  |  |  |  |
|  | PE 20:0_**18:2** | [M-H]^-^ | 786.565 | 295.227 | ω-6 (9, 12) | ω-6 (9, 12)* | ω-7 (8,11) | ω-7 (8,11)* |  |  |
|  | PE **20:3**_18:2 | [M-H]^-^ | 780.518 | 321.242 | ω-6 (8, 11, 14) |  |  |  |  |  |
|  | PG 18:0_**18:2** | [M-H]^-^ | 789.529 | 295.227 | ω-6 (9, 12) | ω-6 (9, 12)* |  |  |  |  |
|  | PI 16:0_**20:4** | [M-H]^-^ | 873.513 | 319.227 | ω-6 (5, 8, 11, 14) |  |  |  |  |  |
|  | PI 16:0_**22:4** | [M-H]^-^ | 901.545 | 347.258 | ω-6 (7, 10, 13, 16) |  |  |  |  |  |
|  | PI 18:0_**18:2** | [M-H]^-^ | 877.545 | 295.227 | ω-6 (9, 12) | ω-7 (8,11) |  |  |  |  |
|  | PI 18:0_**20:3** | [M-H]^-^ | 903.560 | 321.242 | ω-9 (5, 8, 11) | ω-6 (8, 11, 14) |  |  |  |  |
|  | PI 18:0_**20:4** | [M-H]^-^ | 901.545 | 319.227 | ω-6 (5, 8, 11, 14) |  |  |  |  |  |
|  | PI 18:0_**22:4** | [M-H]^-^ | 929.576 | 347.258 | ω-6 (7, 10, 13, 16) |  |  |  |  |  |
|  | PI 18:1_**20:4** | [M-H]^-^ | 899.529 | 319.227 | ω-6 (5, 8, 11, 14) |  |  |  |  |  |
|  | PS 18:0_**18:2** | [M-H]^-^ | 802.524 | 295.227 | ω-6 (9, 12) | ω-7 (8,11) |  |  |  |  |
|  | PS 18:0_**20:4** | [M-H]^-^ | 826.524 | 319.227 | ω-6 (5, 8, 11, 14) |  |  |  |  |  |
|  | PS 18:0_**22:5** | [M-H]^-^ | 852.540 | 345.242 | ω-3 (7, 10, 13, 16, 19) |  |  |  |  |  |
|  | PS 24:5_**20:4** | [M-H]^-^ | 900.540 | 319.227 | ω-6 (5, 8, 11, 14) |  |  |  |  |  |
| Total lipid species: **46**  Identified C=C positional isomers: **96**  * another set of C=C isomers (possibly the *sn*-substitutional isomers separated by LC)  a. Δ12 epoxide was not observed  b. Δ4 and Δ10 epoxides were not observed  c. Δ5 and Δ11 epoxides were not observed  d. Δ5, Δ8, Δ11 mono-epoxides were not observed N/A | | | | | | | | | | |

**S5-3. Representative LC-MS-tPRM results of the identified C=C isomers in the 3T3-L1 adipocyte**

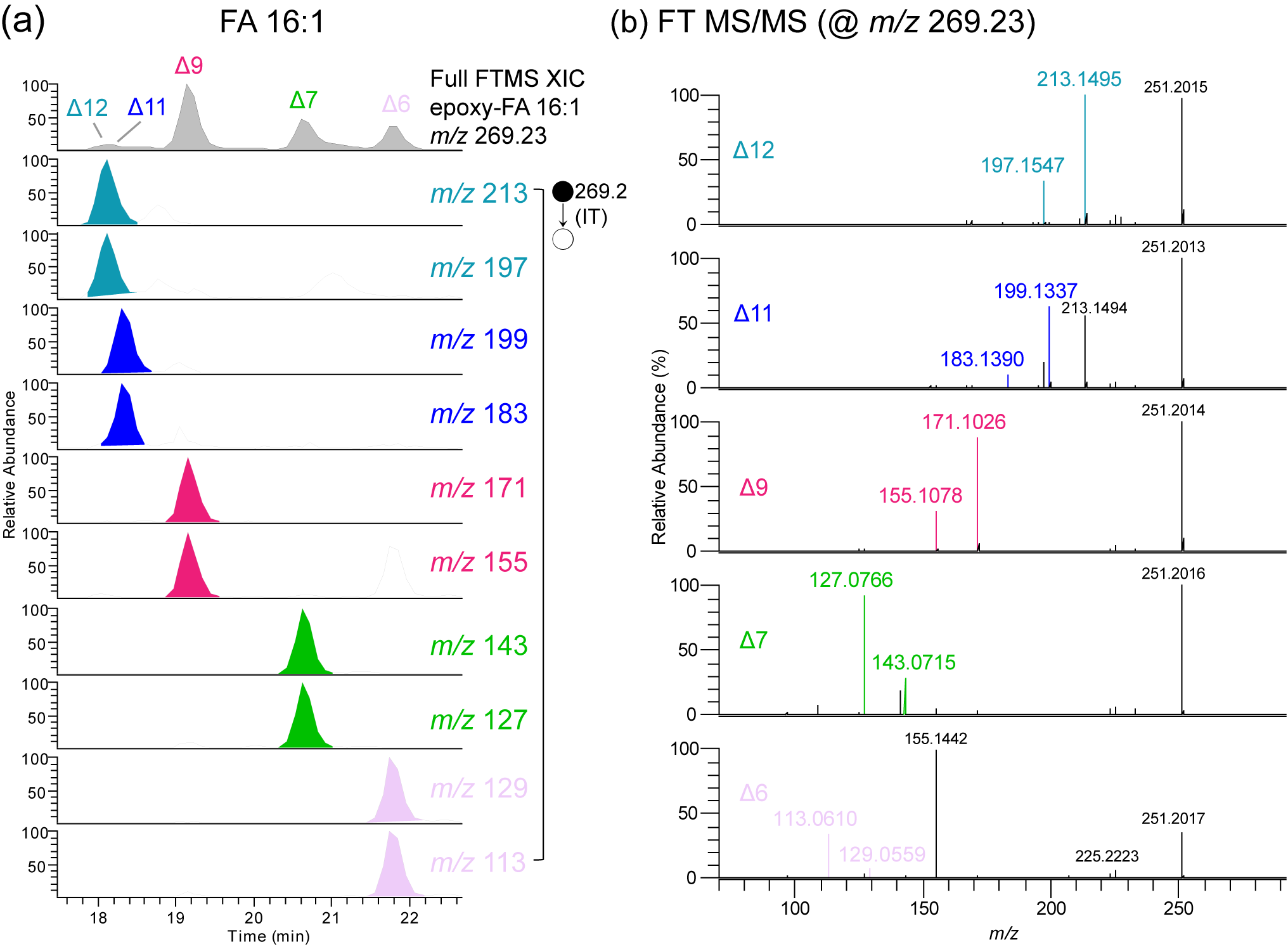

**Figure S22. The identified FA 16:1 C=C isomers in the 3T3-L1 adipocyte.**

**(a)** The representative XIC of the epoxidation products of FA 16:1 at FT-MS1 level and the corresponding C=C diagnostic ions in IT-MS2 level. **(b)** The FT-MS/MS spectra of the epoxides.

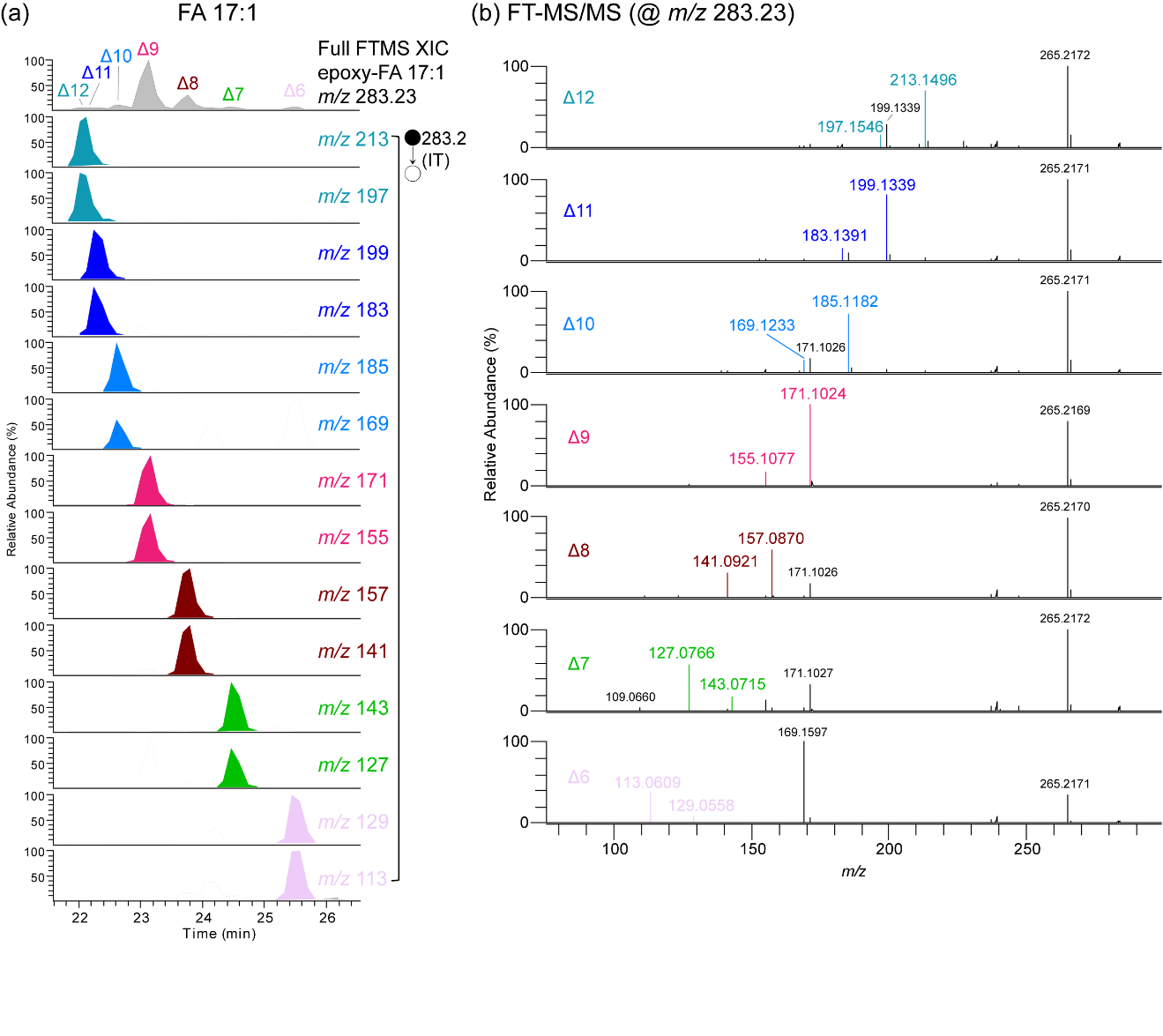

**Figure S23. The identified FA 17:1 C=C isomers in the 3T3-L1 adipocyte.**

**(a)** The representative XIC of the epoxidation products of FA 17:1 at FT-MS1 level and the corresponding C=C diagnostic ions in IT-MS2 level. **(b)** The FT-MS/MS spectra of the epoxides.

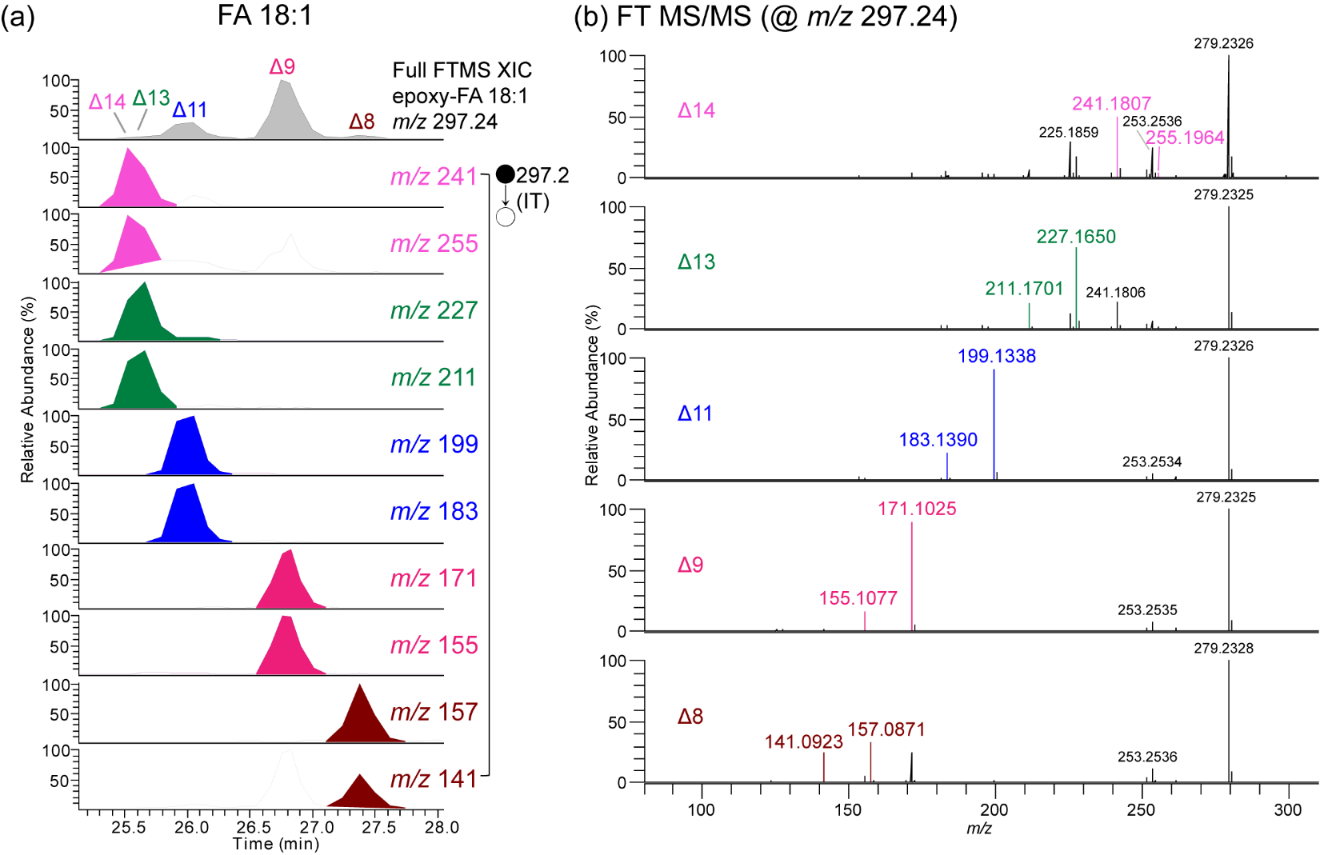

**Figure S24. The identified FA 18:1 C=C isomers in the 3T3-L1 adipocyte.**

**(a)** The representative XIC of the epoxidation products of FA 18:1 at FT-MS1 level and the corresponding C=C diagnostic ions in IT-MS2 level. **(b)** The FT-MS/MS spectra of the epoxides.

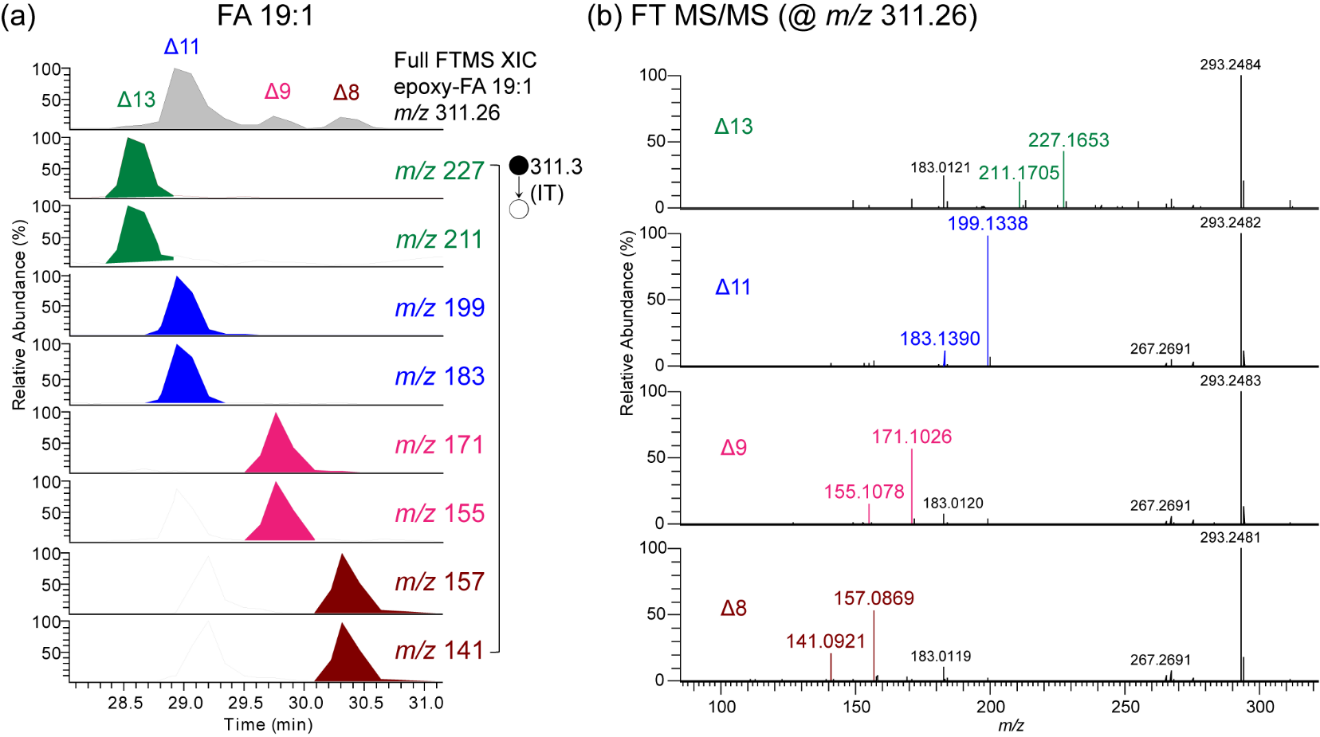

**Figure S25. The identified FA 19:1 C=C isomers in the 3T3-L1 adipocyte.**

**(a)** The representative XIC of the epoxidation products of FA 19:1 at FT-MS1 level and the corresponding C=C diagnostic ions in IT-MS2 level. **(b)** The FT-MS/MS spectra of the epoxides.

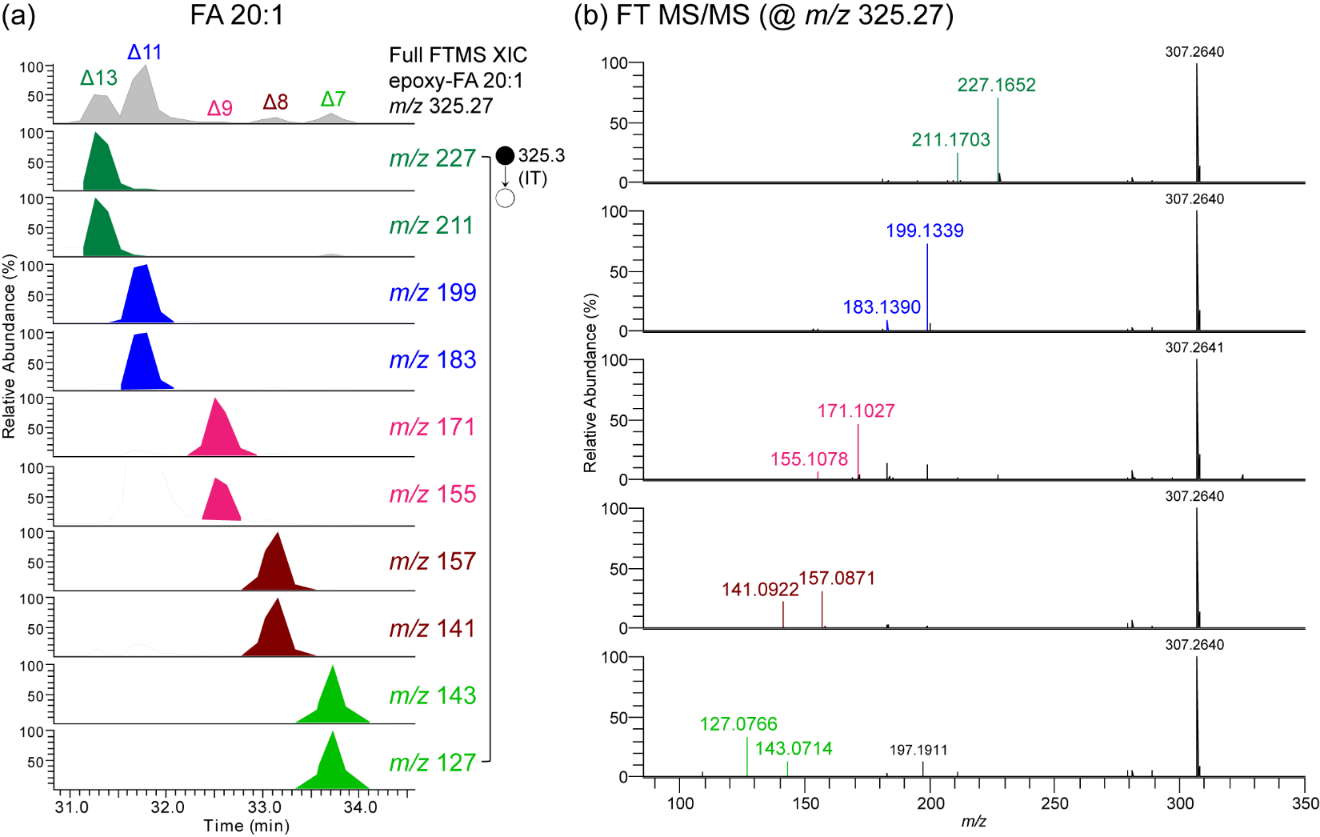

**Figure S26. The identified FA 20:1 C=C isomers in the 3T3-L1 adipocyte.**

**(a)** The representative XIC of the epoxidation products of FA 20:1 at FT-MS1 level and the corresponding C=C diagnostic ions in IT-MS2 level. **(b)** The FT-MS/MS spectra of the epoxides.

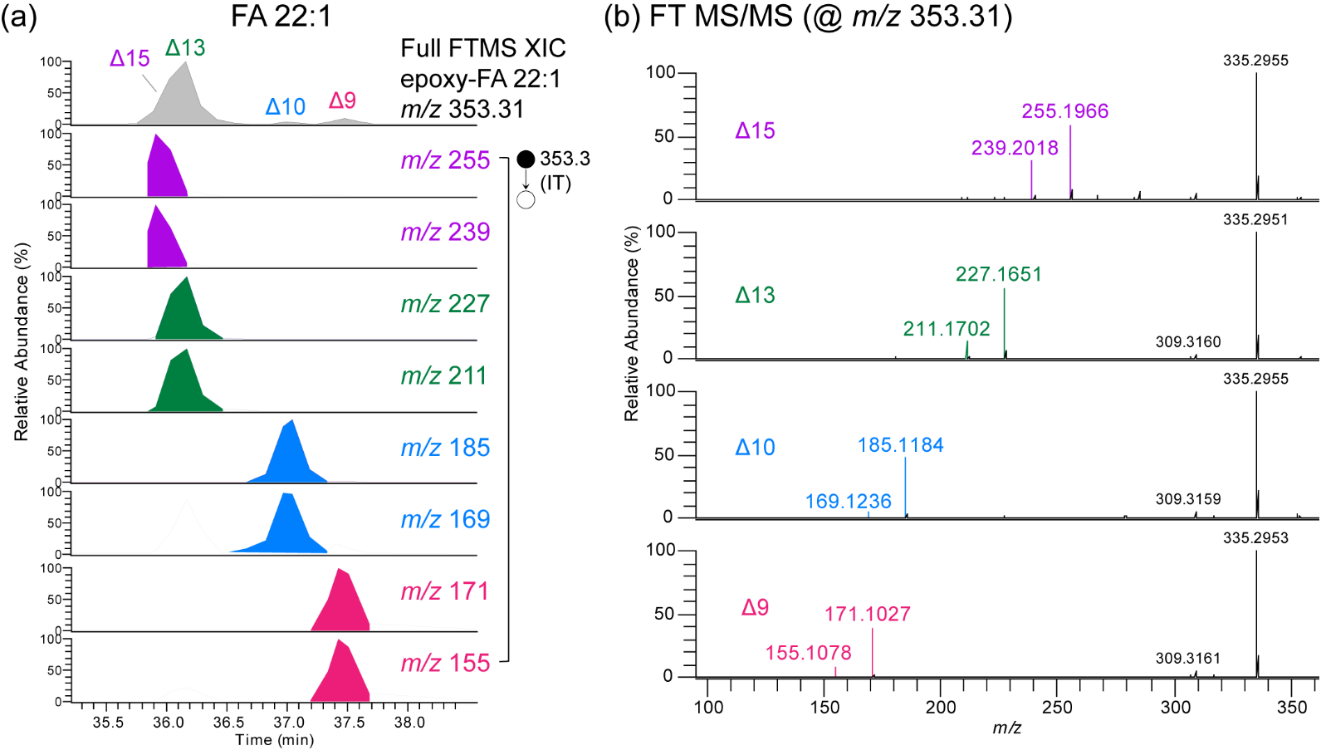

**Figure S27. The identified FA 22:1 C=C isomers in the 3T3-L1 adipocyte.**

**(a)** The representative XIC of the epoxidation products of FA 22:1 at FT-MS1 level and the corresponding C=C diagnostic ions in IT-MS2 level. **(b)** The FT-MS/MS spectra of the epoxides.

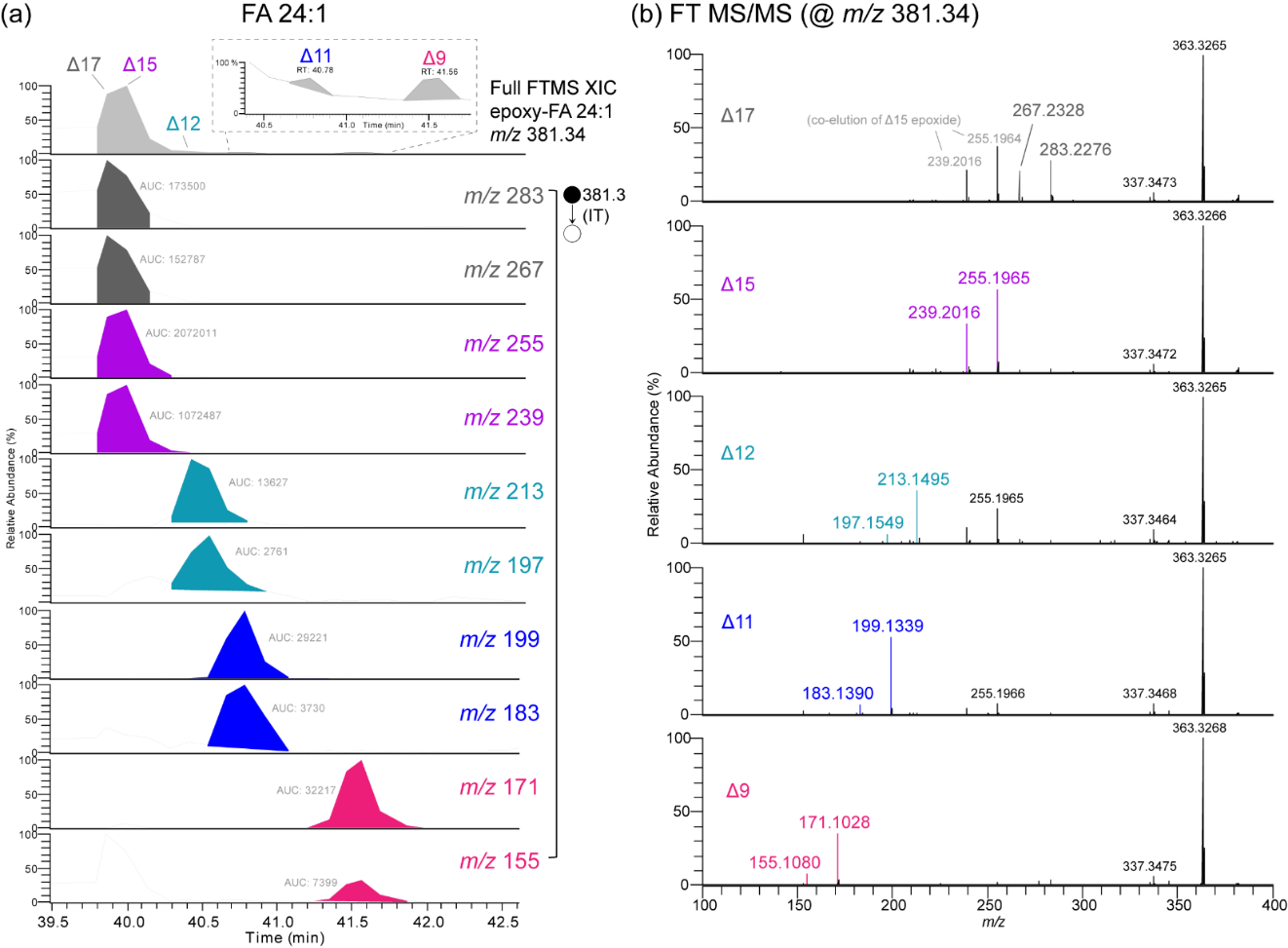

**Figure S28. The identified FA 24:1 C=C isomers in the 3T3-L1 adipocyte.**

**(a)** The representative XIC of the epoxidation products of FA 24:1 at FT-MS1 level and the corresponding C=C diagnostic ions in IT-MS2 level. **(b)** The FT-MS/MS spectra of the epoxides.

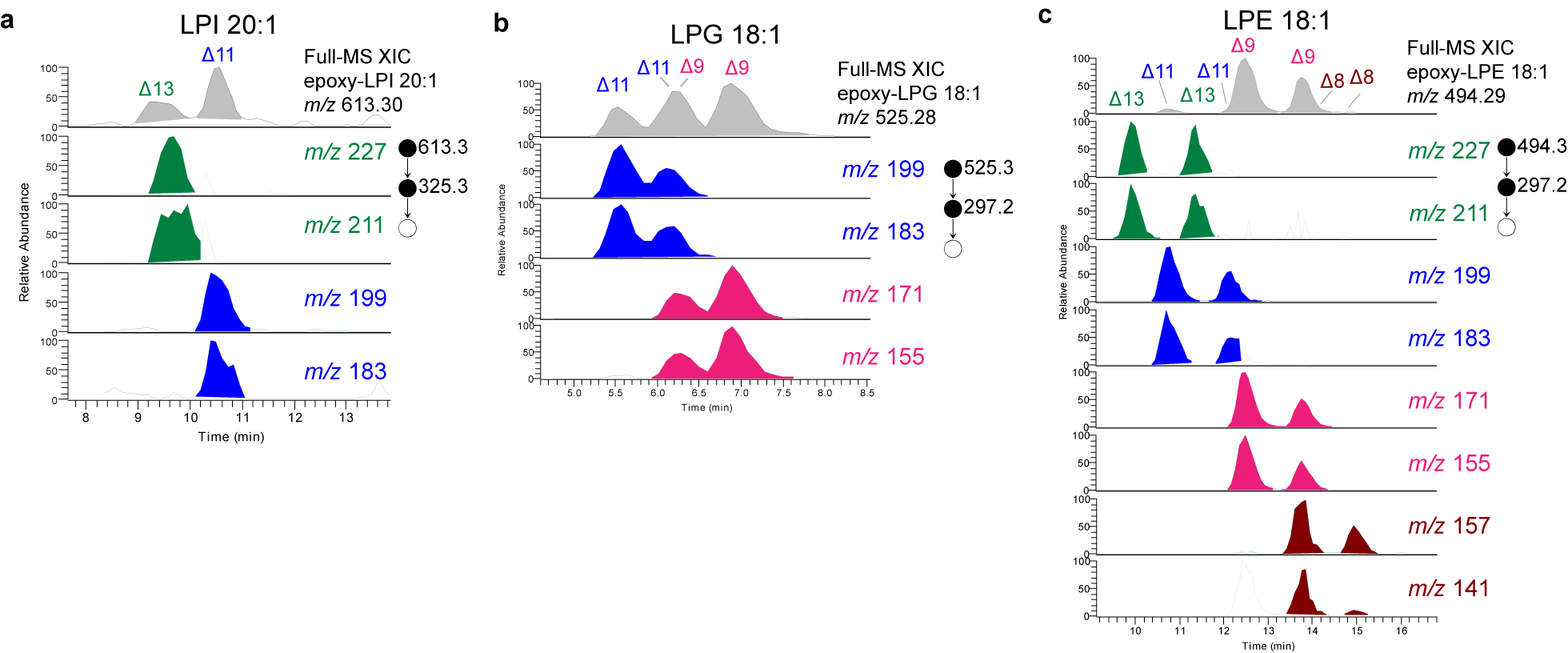

**Figure S29. The identified monounsaturated lyso-GPL C=C isomers in the 3T3-L1 adipocyte.**

The representative XIC of the epoxidation products of **(a)** LPI 20:1 **(b)** LPG 18:1 **(c)** LPE 18:1 at FT-MS1 level and the corresponding C=C diagnostic ions in IT-MS2 level were shown.

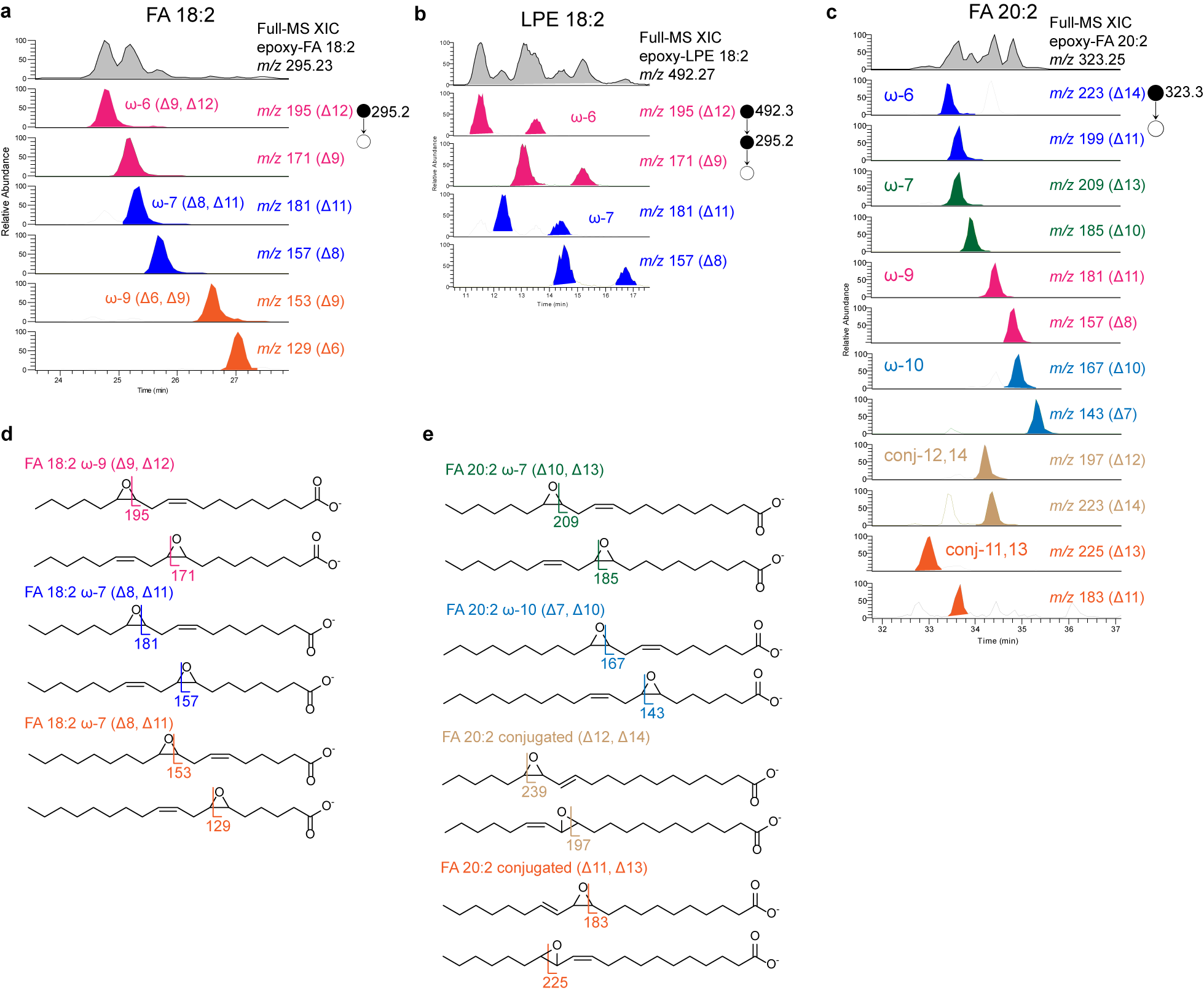

**Figure S30. The identified polyunsaturated isomers with two C=C bonds in the 3T3-L1 adipocyte.**

The representative XIC of the epoxidation products of **(a)** FA 18:2 **(b)** LPE 18:2 **(c)** FA 20:2 at FT-MS1 level and the corresponding C=C diagnostic ions in IT-MS2 (or MS3 for LPE) level were shown. **(d)** The CID scheme of the epoxy-FA 18:2 isomers. **(e)** The CID scheme of the epoxy-FA 20:2 isomers.

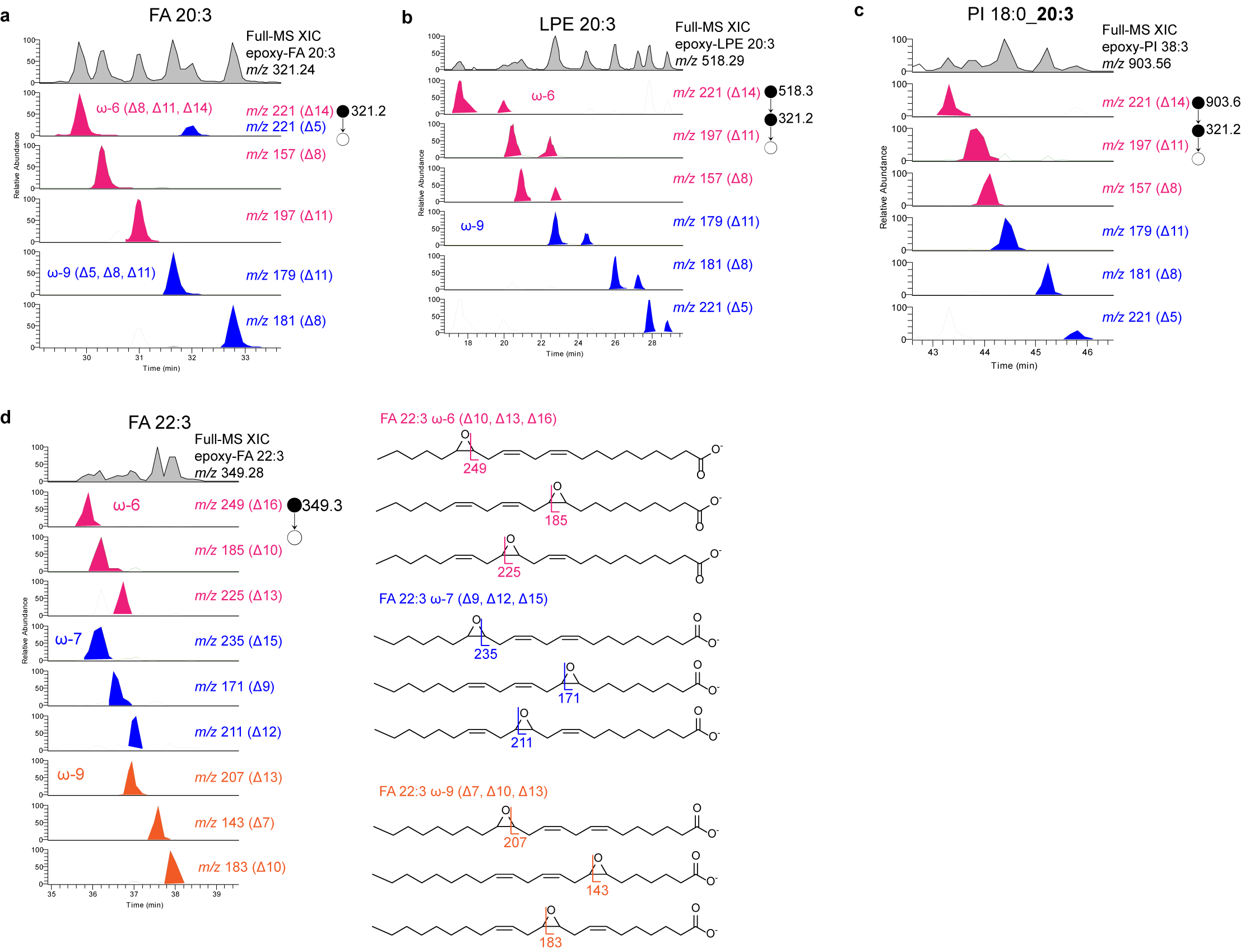

**Figure S31. The identified polyunsaturated isomers with three C=C bonds in the 3T3-L1 adipocyte.**

The representative XIC of the epoxidation products of **(a)** FA 20:3 **(b)** LPE 20:3 **(c)** PI 18:0_20:3 **(d)** FA 22:3 at FT-MS1 level and the corresponding C=C diagnostic ions in IT-MS2 (or MS3 for GPL) level were shown.

**S5-4. Normal vs SCD1 inhibitor-treated**

**Table S12. Quantitative comparison of C=C isomer compositions of monounsaturated lipids between the SCD1 inhibitor-treated and the normal 3T3-L1 adipocytes.**

| Lipid species | C=C position | + DMSO (n= 4) | | | | | | + CAY10655 (n = 4) | | | | | | Two-tailed p-value (Student's t test) | Significant difference * p < 0.05  ** p < 0.01  *** p < 0.005 |
| --- | --- | --- | --- | --- | --- | --- | --- | --- | --- | --- | --- | --- | --- | --- | --- |
|  |  | Sample-1 | Sample-2 | Sample-3 | Sample-4 | Avg. | S.D. | Sample-1 | Sample-2 | Sample-3 | Sample-4 | Avg. | S.D. |  |  |
| FA 16:1 | Δ6 | 7.0 | 3.6 | 4.0 | 3.6 | 4.5 | 1.6 | 6.7 | 7.4 | 6.8 | 7.2 | 7.1 | 0.3 | 5.4E-02 |  |
|  | Δ7 | 16.4 | 18.6 | 19.4 | 19.5 | 18.5 | 1.4 | 16.8 | 17.7 | 16.8 | 17.0 | 17.1 | 0.4 | 1.0E-01 |  |
|  | Δ9 | 58.6 | 71.8 | 70.7 | 70.9 | 68.0 | 6.3 | 62.3 | 59.8 | 61.2 | 62.7 | 61.5 | 1.1 | 1.3E-01 |  |
|  | Δ11 | 4.7 | 2.7 | 2.5 | 2.7 | 3.2 | 1.0 | 5.0 | 5.0 | 5.0 | 4.7 | 4.9 | 0.1 | **3.9E-02** | * |
|  | Δ12 | 13.3 | 3.3 | 3.4 | 3.3 | 5.8 | 5.0 | 9.2 | 10.1 | 10.2 | 8.4 | 9.4 | 0.7 | 2.4E-01 |  |
| FA 17:1 | Δ6 | 0.7 | 1.2 | 1.1 | 1.0 | 1.0 | 0.2 | 1.5 | 1.4 | 1.7 | 1.5 | 1.6 | 0.1 | **4.1E-03** | *** |
|  | Δ7 | 1.6 | 1.5 | 1.6 | 1.7 | 1.6 | 0.1 | 1.9 | 1.5 | 1.8 | 1.7 | 1.7 | 0.1 | 2.1E-01 |  |
|  | Δ8 | 32.1 | 17.2 | 23.2 | 20.2 | 23.2 | 6.5 | 13.0 | 13.3 | 11.1 | 12.2 | 12.6 | 1.0 | **4.3E-02** | * |
|  | Δ9 | 38.2 | 66.6 | 55.0 | 59.4 | 54.8 | 12.0 | 69.0 | 69.7 | 70.8 | 71.2 | 69.8 | 1.2 | 8.3E-02 |  |
|  | Δ10 | 22.1 | 8.0 | 13.3 | 12.2 | 13.9 | 5.9 | 6.1 | 6.0 | 4.9 | 5.8 | 5.8 | 0.6 | 6.9E-02 |  |
|  | Δ11 | 2.2 | 3.2 | 3.0 | 3.1 | 2.9 | 0.4 | 5.3 | 4.8 | 6.1 | 4.8 | 5.2 | 0.6 | **8.0E-04** | *** |
|  | Δ12 | 2.9 | 2.4 | 2.9 | 2.4 | 2.7 | 0.3 | 3.3 | 3.2 | 3.6 | 3.0 | 3.3 | 0.2 | **1.8E-02** | * |
| FA 18:1 | Δ8 | 2.2 | 2.1 | 2.2 | 2.2 | 2.2 | 0.1 | 3.2 | 2.8 | 3.0 | 3.2 | 3.0 | 0.2 | **1.8E-04** | *** |
|  | Δ9 | 67.4 | 70.7 | 70.1 | 70.8 | 69.7 | 1.6 | 62.4 | 62.3 | 63.2 | 64.5 | 63.1 | 1.0 | **4.4E-04** | *** |
|  | Δ11 | 28.0 | 25.0 | 25.5 | 24.8 | 25.8 | 1.5 | 31.6 | 32.0 | 31.3 | 29.7 | 31.2 | 1.0 | **1.1E-03** | *** |
|  | Δ13 | 1.8 | 1.7 | 1.7 | 1.7 | 1.7 | 0.1 | 2.1 | 2.1 | 1.9 | 1.9 | 2.0 | 0.1 | **2.9E-03** | *** |
|  | Δ14 | 0.7 | 0.6 | 0.5 | 0.6 | 0.6 | 0.1 | 0.8 | 0.8 | 0.6 | 0.7 | 0.7 | 0.1 | 5.4E-02 |  |
| FA 19:1 | Δ8 | 7.6 | 8.0 | 6.1 | 7.9 | 7.4 | 0.9 | 7.7 | 6.9 | 7.7 | 8.1 | 7.6 | 0.5 | 7.0E-01 |  |
|  | Δ9 | 10.8 | 10.7 | 7.5 | 10.5 | 9.9 | 1.6 | 8.3 | 8.7 | 8.3 | 7.8 | 8.3 | 0.4 | 1.4E-01 |  |
|  | Δ11 | 77.4 | 78.1 | 83.5 | 78.5 | 79.4 | 2.8 | 81.6 | 81.5 | 81.4 | 82.0 | 81.6 | 0.2 | 2.0E-01 |  |
|  | Δ13 | 4.3 | 3.2 | 2.9 | 3.1 | 3.4 | 0.6 | 2.4 | 2.9 | 2.6 | 2.1 | 2.5 | 0.3 | **4.5E-02** | ***** |
| FA 20:1 | Δ7 | 3.6 | 3.5 | 3.4 | 3.1 | 3.4 | 0.2 | 3.7 | 4.4 | 4.3 | 3.9 | 4.1 | 0.3 | **1.5E-02** | ***** |
|  | Δ8 | 3.6 | 3.5 | 3.6 | 3.2 | 3.5 | 0.2 | 5.5 | 5.6 | 5.9 | 5.9 | 5.7 | 0.2 | **2.9E-06** | *** |
|  | Δ9 | 1.1 | 1.1 | 1.0 | 1.1 | 1.1 | 0.1 | 1.3 | 1.8 | 1.3 | 1.4 | 1.4 | 0.2 | **1.7E-02** | * |
|  | Δ11 | 61.1 | 62.7 | 60.8 | 58.0 | 60.6 | 1.9 | 58.1 | 54.7 | 61.7 | 59.8 | 58.6 | 3.0 | 2.8E-01 |  |
|  | Δ13 | 30.7 | 29.2 | 31.3 | 34.6 | 31.4 | 2.3 | 31.5 | 33.6 | 26.7 | 29.1 | 30.2 | 3.0 | 5.4E-01 |  |
| FA 22:1 | Δ9 | 8.0 | 9.0 | 8.7 | 7.9 | 8.4 | 0.5 | 9.4 | 8.3 | 8.7 | 11.2 | 9.4 | 1.3 | 2.1E-01 |  |
|  | Δ10 | 2.8 | 2.7 | 2.8 | 2.5 | 2.7 | 0.1 | 1.9 | 1.7 | 2.0 | 3.0 | 2.2 | 0.6 | 1.7E-01 |  |
|  | Δ13 | 75.0 | 75.4 | 75.2 | 74.7 | 75.1 | 0.3 | 78.5 | 81.0 | 83.5 | 73.2 | 79.0 | 4.4 | 1.7E-01 |  |
|  | Δ15 | 14.2 | 12.9 | 13.3 | 14.9 | 13.8 | 0.9 | 10.1 | 9.1 | 5.9 | 12.6 | 9.4 | 2.8 | **2.3E-02** | * |
| FA 24:1 | Δ9 | 1.3 | 1.2 | 1.1 | 1.3 | 1.2 | 0.1 | 3.2 | 2.9 | 3.4 | 3.2 | 3.2 | 0.2 | **2.57E-07** | *** |
|  | Δ11 | 1.0 | 1.1 | 0.9 | 1.1 | 1.0 | 0.1 | 2.4 | 2.5 | 2.2 | 2.3 | 2.3 | 0.1 | **2.11E-07** | *** |
|  | Δ12 | 0.6 | 0.5 | 0.5 | 0.6 | 0.6 | 0.04 | 0.8 | 1.4 | 1.2 | 1.4 | 1.2 | 0.3 | **4.52E-03** | *** |
|  | Δ15 | 87.1 | 87.2 | 87.6 | 85.4 | 86.8 | 1.0 | 76.3 | 74.7 | 76.5 | 76.0 | 76.1 | 0.9 | **5.6E-07** | *** |
|  | Δ17 | 10.0 | 10.0 | 9.9 | 11.7 | 10.4 | 0.9 | 17.3 | 18.5 | 16.8 | 17.1 | 17.5 | 0.7 | **2.38E-06** | *** |
| LPE 18:1 | Δ8 | 1.1 | 1.0 | 1.2 | 1.2 | 1.1 | 0.1 | 2.1 | 2.1 | 2.0 | 2.1 | 2.0 | 0.1 | **9.6E-06** | *** |
|  | Δ9 | 91.1 | 94.7 | 91.4 | 90.9 | 92.0 | 1.8 | 90.3 | 89.5 | 90.3 | 89.8 | 90.0 | 0.4 | 1.1E-01 |  |
|  | Δ11 | 7.6 | 4.2 | 7.3 | 7.8 | 6.7 | 1.7 | 7.5 | 8.2 | 7.6 | 8.0 | 7.8 | 0.4 | 3.0E-01 |  |
|  | Δ13 | 0.1 | 0.1 | 0.1 | 0.1 | 0.1 | 0.0 | 0.1 | 0.2 | 0.1 | 0.1 | 0.1 | 0.0 | 3.7E-01 |  |
| LPG 18:1 | Δ9 | 53.7 | 58.9 | 55.4 | 58.8 | 56.7 | 2.6 | 39.3 | 39.9 | 40.6 | 43.6 | 40.8 | 1.9 | **6.0E-05** | *** |
|  | Δ11 | 46.3 | 41.1 | 44.6 | 41.2 | 43.3 | 2.6 | 60.7 | 60.1 | 59.4 | 56.4 | 59.2 | 1.9 | **6.0E-05** | *** |
| LPI 18:1 | Δ8 | 2.1 | 1.8 | 2.3 | 1.9 | 2.0 | 0.2 | 3.6 | 4.6 | 5.1 | 5.0 | 4.6 | 0.7 | **3.8E-04** | *** |
|  | Δ9 | 62.3 | 66.6 | 65.8 | 61.7 | 64.1 | 2.4 | 58.1 | 54.8 | 55.3 | 56.4 | 56.1 | 1.5 | **1.4E-03** | *** |
|  | Δ11 | 34.2 | 30.6 | 30.4 | 35.3 | 32.6 | 2.5 | 36.2 | 38.2 | 37.7 | 36.4 | 37.1 | 1.0 | **1.5E-02** | * |
|  | Δ13 | 1.4 | 1.0 | 1.5 | 1.1 | 1.2 | 0.2 | 2.1 | 2.5 | 1.9 | 2.3 | 2.2 | 0.2 | **1.2E-03** | *** |
| LPI 20:1 | Δ9 | 72.4 | 81.8 | 84.5 | 74.9 | 78.4 | 5.7 | 68.6 | 72.7 | 70.5 | 72.9 | 71.2 | 2.0 | 5.4E-02 |  |
|  | Δ11 | 27.6 | 18.2 | 15.5 | 25.1 | 21.6 | 5.7 | 31.4 | 27.3 | 29.5 | 27.1 | 28.8 | 2.0 | 5.4E-02 |  |
| PG **18:1**_16:0 | Δ9 | 23.9 | 25.1 | 22.0 | 26.8 | 24.4 | 2.0 | 18.2 | 24.0 | 17.8 | 21.4 | 20.3 | 2.9 | 6.1E-02 |  |
|  | Δ11 | 75.9 | 74.7 | 77.8 | 73.0 | 75.3 | 2.0 | 81.3 | 75.5 | 81.9 | 78.3 | 79.3 | 3.0 | 7.1E-02 |  |
|  | Δ13 | 0.2 | 0.3 | 0.2 | 0.2 | 0.2 | 0.0 | 0.4 | 0.5 | 0.3 | 0.3 | 0.4 | 0.1 | **3.6E-02** | * |
| PG **18:1**_**18:1** | Δ9 | 68.9 | 72.4 | 71.6 | 70.9 | 71.0 | 1.5 | 55.6 | 55.7 | 60.5 | 59.2 | 57.8 | 2.5 | **9.8E-05** | *** |
|  | Δ11 | 30.4 | 27.0 | 27.7 | 28.8 | 28.5 | 1.5 | 43.4 | 43.3 | 38.5 | 39.7 | 41.2 | 2.5 | **1.2E-04** | *** |
|  | Δ13 | 0.7 | 0.6 | 0.7 | 0.3 | 0.6 | 0.2 | 1.1 | 1.0 | 1.1 | 1.1 | 1.0 | 0.1 | **4.9E-03** | *** |
| PI **16:1**_18:0 | Δ7 | 66.1 | 60.5 | 64.5 | 65.6 | 64.2 | 2.5 | 62.9 | 57.8 | 54.5 | 59.0 | 58.6 | 3.5 | **3.9E-02** | * |
|  | Δ9 | 33.9 | 39.5 | 35.5 | 34.4 | 35.8 | 2.5 | 37.1 | 42.2 | 45.5 | 41.0 | 41.4 | 3.5 | **3.9E-02** | * |
| PI **18:1**_16:0 | Δ9 | 77.7 | 82.6 | 79.0 | 79.0 | 79.6 | 2.1 | 61.9 | 60.6 | 61.7 | 61.3 | 61.4 | 0.6 | **2.9E-06** | *** |
|  | Δ11 | 22.3 | 17.4 | 21.0 | 21.0 | 20.4 | 2.1 | 38.1 | 39.4 | 38.3 | 38.7 | 38.6 | 0.6 | **2.9E-06** | *** |
| PI **18:1**_20:4 | Δ8 | 1.5 | 1.7 | 1.7 | 1.2 | 1.5 | 0.2 | 8.6 | 8.5 | 9.5 | 8.1 | 8.7 | 0.6 | **4.6E-07** | *** |
|  | Δ9 | 68.2 | 67.3 | 68.5 | 70.4 | 68.6 | 1.3 | 55.6 | 54.4 | 59.0 | 54.3 | 55.8 | 2.2 | **5.9E-05** | *** |
|  | Δ11 | 28.9 | 29.6 | 28.2 | 27.6 | 28.6 | 0.9 | 33.8 | 34.5 | 29.7 | 35.5 | 33.4 | 2.5 | **1.2E-02** | * |
|  | Δ13 | 1.3 | 1.4 | 1.6 | 0.8 | 1.3 | 0.3 | 1.9 | 2.6 | 1.8 | 2.1 | 2.1 | 0.3 | **1.3E-02** | * |
| PE **18:1**_**18:1** | Δ9 | 83.1 | 84.4 | 84.7 | 84.3 | 84.1 | 0.7 | 87.2 | 89.2 | 86.6 | 85.7 | 87.2 | 1.5 | **9.7E-03** | ** |
|  | Δ11 | 16.4 | 15.1 | 14.9 | 15.3 | 15.4 | 0.7 | 12.3 | 10.3 | 12.9 | 13.7 | 12.3 | 1.4 | **7.3E-03** | ** |
|  | Δ13 | 0.5 | 0.5 | 0.4 | 0.4 | 0.4 | 0.1 | 0.5 | 0.4 | 0.5 | 0.6 | 0.5 | 0.1 | 7.2E-02 |  |
| PE **16:1**_18:0 | Δ6 | 4.6 | 3.9 | 3.6 | 3.4 | 3.9 | 0.6 | 5.4 | 5.2 | 5.4 | 4.7 | 5.2 | 0.3 | **5.9E-03** | * |
|  | Δ7 | 14.2 | 13.9 | 12.1 | 12.3 | 13.1 | 1.1 | 10.2 | 11.1 | 11.1 | 10.9 | 10.8 | 0.4 | **8.1E-03** | ** |
|  | Δ9 | 78.7 | 79.3 | 82.1 | 82.1 | 80.6 | 1.8 | 81.5 | 81.1 | 80.7 | 81.9 | 81.3 | 0.5 | 4.6E-01 |  |
|  | Δ11 | 2.4 | 2.9 | 2.3 | 2.3 | 2.5 | 0.3 | 2.9 | 2.7 | 2.8 | 2.4 | 2.7 | 0.2 | 2.6E-01 |  |
| PE **18:1**_16:0 | Δ9 | 87.2 | 86.3 | 85.8 | 87.2 | 86.6 | 0.7 | 89.4 | 87.1 | 89.6 | 88.4 | 88.6 | 1.2 | **2.6E-02** | * |
|  | Δ11 | 12.8 | 13.7 | 14.2 | 12.8 | 13.4 | 0.7 | 10.6 | 12.9 | 10.4 | 11.6 | 11.4 | 1.2 | **2.6E-02** | * |
| PS **18:1**_18:0 | Δ9 | 80.3 | 83.3 | 85.6 | 84.8 | 83.5 | 2.4 | 86.8 | 88.5 | 89.1 | 90.1 | 88.7 | 1.4 | **9.2E-03** | ** |
|  | Δ11 | 19.7 | 16.7 | 14.4 | 15.2 | 16.5 | 2.4 | 13.2 | 11.5 | 10.9 | 9.9 | 11.3 | 1.4 | **9.2E-03** | ** |

**Table S13. Quantitative comparison of C=C isomer compositions of FA 20:3 and its constituting GPLs in the SCD1 inhibitor-treated and the normal 3T3-L1 adipocytes.**

| Lipid species | C=C position | + DMSO (n = 4) | | | | | | + CAY10655 (n = 4) | | | | | | Two-tailed p-value (Student's t test) | Significant difference  *** p < 0.001 |
| --- | --- | --- | --- | --- | --- | --- | --- | --- | --- | --- | --- | --- | --- | --- | --- |
|  |  | Sample-1 | Sample-2 | Sample-3 | Sample-4 | Avg. | S.D. | Sample-1 | Sample-2 | Sample-3 | Sample-4 | Avg. | S.D. |  |  |
| FA 20:3 | ω-6 (Δ8, Δ11, Δ14) | 51.2 | 52.8 | 49.8 | 53.0 | 51.7 | 1.5 | 74.4 | 81.4 | 77.6 | 74.4 | 77.0 | 3.3 | 8.9E-06 | *** |
|  | ω-9 (Δ5, Δ8, Δ11) | 48.8 | 47.2 | 50.2 | 47.0 | 48.3 | 1.5 | 25.6 | 18.6 | 22.4 | 25.6 | 23.0 | 3.3 | 8.9E-06 | *** |
| LPE 20:3 | ω-6 (Δ8, Δ11, Δ14) | 43.6 | 43.2 | 43.8 | 40.6 | 40.9 | 5.8 | 62.6 | 64.4 | 60.2 | 60.5 | 63.0 | 2.4 | 4.2E-04 | *** |
|  | ω-9 (Δ5, Δ8, Δ11) | 56.4 | 56.8 | 56.2 | 59.4 | 59.1 | 5.8 | 37.4 | 35.6 | 39.8 | 39.5 | 37.0 | 2.4 | 4.2E-04 | *** |
| PI 18:0_20:3 | ω-6 (Δ8, Δ11, Δ14) | 28.4 | 26.9 | 28.2 | 29.9 | 28.3 | 1.2 | 71.8 | 68.7 | 66.9 | 67.2 | 68.6 | 2.3 | 7.4E-08 | *** |
|  | ω-9 (Δ5, Δ8, Δ11) | 71.6 | 73.1 | 71.8 | 70.1 | 71.7 | 1.2 | 28.2 | 31.3 | 33.1 | 32.8 | 31.4 | 2.3 | 7.4E-08 | *** |

**S6.**

**S6. MELDI-DESI-MS imaging platform for mapping C=C isomers in tissue sections**

**S6-1. Quantification of C=C isomer standards using MELDI-DESI-MS/MS**

Herein the ability of MELDI-DESI-MS/MS in quantifying C=C isomers was validated. We prepared a series of isomeric mixtures of FA 18:1 (9Z) and FA 18:1 (11Z), with the 9Z isomer proportion varying in 0 %, 20 %, 40 %, 60 %, 80 % and 100 %, and total concentration of FA 18:1 was held at 100 mM. For reproducible sample deposition, we used the Omni Slide Micro-24 Array (Prosolia, Inc.), a glass slide with a printed 24 hydrophobic spot array, whereas each spot had a diameter of 3-mm and a center-to-center separation of 4 mm. Then, 2 µL of each isomer mixture (200 pmole of FA) was deposited onto the slide and dried. Subsequently, the epoxidation reagent (200 mM mCPBA in chloroform) was prepared, and 2 uL of the reagent was deposited onto each spot for on-site epoxidation in ambient, and the sample would dry in several seconds (**Fig. 32a**). The sample array was examined by targeted DESI-IT-MS/MS in the line-scanning mode (moving speed: 150 µm/sec; maximum injection time: 500 ms; acquisition time ~1.6 sec), in which the precursor ion was set at *m/z* 297.2, the deprotonated ion of epoxy-FA 18:1. As shown in **Fig S32b**, we then extracted the ion chromatograms of *m/z* 171 and *m/z* 199 (the diagnostic ions for Δ9 and Δ11, respectively), and their area under curve (AUC) in each sample spot was calculated, denoted as A*_m/z_* _171_ and A*_m/z_* _199_. The calibration curve for quantification of Δ9/Δ11 isomer mixtures was constructed by plotting fractions of Δ9-AUC against the molar fractions of FA 18:1 (9Z) isomer (**Fig. S32c**), showing a good linearity (R^2^ = 0.995). This result suggested that quantification of Δ9/Δ11 isomer proportions in FA 18:1 and its constituting GPLs was feasible using MELDI-DESI-MS/MS.

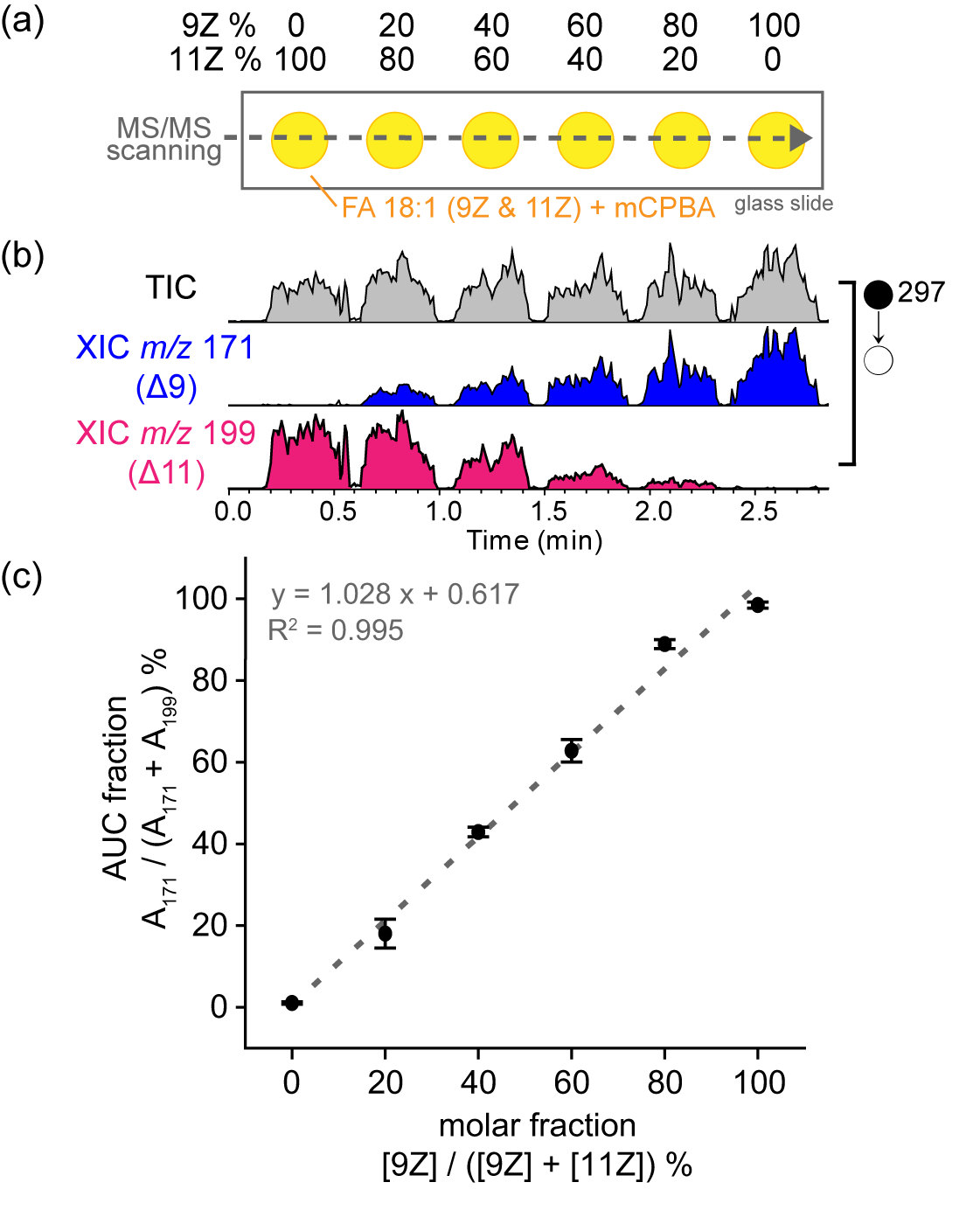

**Figure S32. Quantification of C=C isomer standards using MELDI-DESI-MS/MS.**

**(a)** Quantification of C=C isomers was validated using a series of isomeric mixtures of FA 18:1 (9Z) and FA 18:1 (11Z), with 9Z isomer proportion varying in 0 %, 20 %, 40 %, 60 %, 80 % and 100 %, and total amount of FA 18:1 was held at 200 nmole. The mixtures were deposited onto an Omni Slide with hydrophobic arrays (yellow), epoxidized by mCPBA, and analyzed by targeted DESI IT-MS/MS at the precursor ion *m/z* 297 (deprotonated epoxy-FA 18:1). **(b)** The XIC AUC of C=C diagnostic ions (A*_m/z_* _171_ for 9Z; A*_m/z_* _199_ for 11Z) was extracted. **(c)** The calibration curve was constructed by plotting AUC fractions against the molar fractions of FA 18:1 (9Z) isomer. Error bars represent standard deviations calculated from technical triplicates.

**S6-2 Limit of detection of C=C diagnostic ions with MELDI-DESI MS/MS.**

A series of FA 18:1 (9Z) standards (10^2^, 10^1^, 10^0^, 10^-1^, 10^-2^ mM, and a blank sample) were examined similarly. Again, 2 µL of each sample was deposited (10^4^, 10^3^, 10^2^, 10^1^, 10^0^, and 0 pmole of FA), dried, reacted with mCPBA, subjected to DESI-MS/MS line-scanning analysis (**Fig S33a)**. The AUC of *m/z* 171 was then extracted and plotted against the number of FA molecule per spot (**Fig. S33b**). As a result, we showed that the ion signal from 10 to 1 pmole was not significantly changed, and the LOD was concluded in the range of 10~100 pmole (**Fig. S33c**).

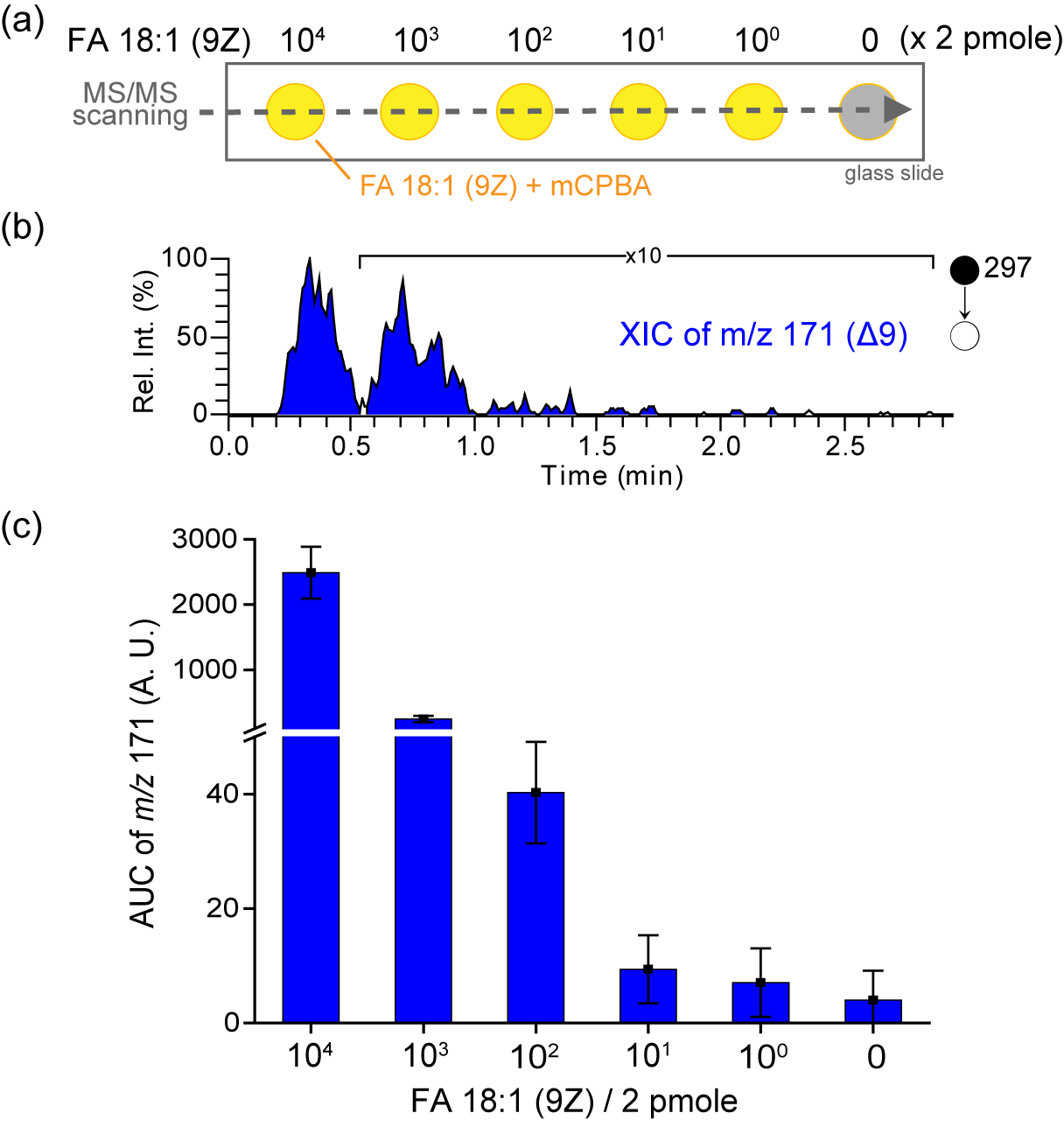

**Figure S33. Limit of detection of C=C diagnostic ions of FA 18:1 (9Z) with MELDI-DESI MS/MS.**

**(a)** Limit of detection (LOD) of C=C isomers was validated using a series of FA 18:1 (9Z) standards and a blank sample. **(b)** The XIC of the Δ9 diagnostic ion (*m/z* 171) under the MS/MS channel at *m/z* 297. **(c)** The AUC of the diagnostic ions corresponding to the samples. Error bars represent standard deviation calculated out of a technical triplicate.

**S6-3. Development of an airbrushing method for *in situ* epoxidation in biological tissue sections**

**S6-3-1. Method development**

We demonstrated a protocol for *in situ* epoxidation of unsaturated lipids on tissue sections for mapping C=C isomers by DESI-MS. The protocol was modified from an airbrush method for matrix deposition in matrix-assisted laser desorption ionization mass spectrometry imaging (MALDI-MSI) experiment, which was previously developed by Prof. Richard M. Caprioli’s group ^6^. Using an airbrush, matrix solution could be homogeneously coated on the tissue surface, forming a thin layer of matrix. Due to its inexpensiveness, easiness, simple operation, and commercial availability, the airbrush method was the most used matrix application method if an expensive robotic sprayer was not accessible.

To demonstrate the universal accessibility of MELDI in mapping C=C isomer *in situ* for the wide mass spectrometric community, we thus utilized and modified the conventional airbrushing protocol for this purpose. First, we replaced the matrix solution in the previous protocol in MALDI-MSI experiment with the epoxidation reagent (100 mM mCPBA, dissolved in methanol). Importantly, reproducible and efficient *in situ* epoxidation was enabled by using pure methanol as the solvent system. We found that both an aqueous solution (water/methanol = 1/1) and dimethylformamide/acetonitrile (v/v = 1/1) solvent that were used in routine DESI-MSI experiment could cause severe droplet accumulation and bad imaging quality. Also, a highly non-polar solvent (e.g. ethyl acetate, dichloromethane, and chloroform) was rapidly evaporated before reaching to the tissue surface, resulting in inefficient derivatization. Finally, the concentration of mCPBA was kept at 100 mM as we found that derivatization using a lower concentration of mCPBA (10 mM) was not efficient, while the higher concentration (e.g. 500 mM ~ 1M) caused severe ion suppression.

The developed method was briefly illustrated in **Fig. S34c**, and a demonstration video was provided (see **section S7-3-2)**. First, the derivatization solution was sprayed onto the tissue section with a controllable commercial airbrush kit. Then, a low solvent flow was adjusted, and the mCPBA crystal outside the tissue margin was carefully removed with methanol. Finally, the remained mCPBA was cleaned prior to DESI-MSI analysis.

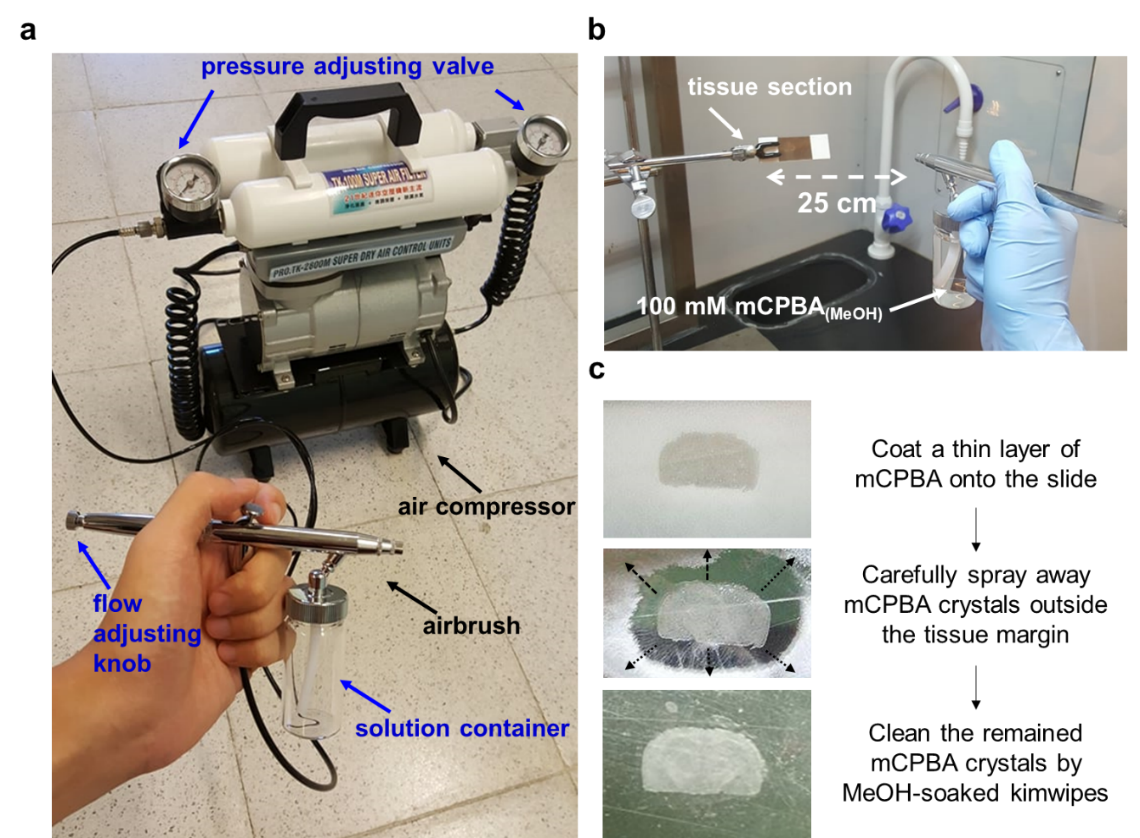

**Figure S34. Illustration of *in situ* epoxidation by spraying mCPBA.**

**(a)** The commercial airbrush kit, which contained an airbrush, and an air compressor with pressures adjusting valves. **(b)** The experimental setup for mCPBA airbrushing on a tissue section. **(c)** Experimental snapshots of the tissue section.

**S6-3-2. Demonstration video**

A demonstration video of mCPBA spraying using an airbrush for *in situ* epoxidation on tissue sections was available at <https://drive.google.com/drive/folders/13Plb4-aY0NnMpS6pT9UURBMOWCDgkQ-N/>

**S6-3-3. Protocol**

- **Materials**

1. mCPBA (powder; ≦77% purity; Sigma Aldrich)
2. Methanol (Honeywell, U.S.A.)
3. Stainless steel plate (76×26×1 mm; Tianmao, Taiwan)
4. Airbrush (RH-BS; Prona, Taiwan)
5. Air compressor (CPM-280A; Xian-Ying, Taiwan)
6. Tissue section on a silane-coating slide (76×26×1 mm; Yeong Jyi Chemical Apparatus, Taiwan)

- **Notification**

For safety issues, the experimental should be carried out in a chemical hood. People should wear face masker and gloves.

- **Experimental steps**

1. **Prepare mCPBA solution (100 mM)**

Add 1.115 g of mCPBA powder into a 50-mL falcon tube and add methanol to total volume of 50 mL to form 100 mM solution.

2. **Spray mCPBA solution onto the tissue section**
   1) Fix the tissue slide on a stainless steel plate by tapes and hold the slide on a three prong clamp.
   2) Add 25 mL of mCPBA solution into the solution container of the airbrush.
   3) Adjust the flow adjusting knob of the airbrush at 1.5 turn. (in high flow mode)
   4) Turn on the air compressor and adjust the air flow pressure at 30 psi.
   5) Make sure that the airbrush can spray fluently.
   6) Spray the mCPBA solution homogeneously onto the tissue with the spray head-to-tissue distance of 20-30 cm.
   7) A thin layer of mCPBA crystals forms after ~20 mL of mCPBA solution has been used.
3. **Clean the airbrush by MeOH**
   1) Wash the solution container with pure MeOH and then add 25 mL MeOH into the solution container.
   2) Clean the solution channel by spraying MeOH with higher flow rate.
   3) Clean the spray head by soaking it into MeOH.
4. **Remove the remained mCPBA crystals outside the tissue margin**
   1) Adjust the flow adjusting knob of the airbrush at 0.25 turn. (in low flow mode)
   2) Carefully clean the mCPBA crystals outside the tissue margin by spraying MeOH in the direction away from tissue section and toward the slide edge.
   3) Use a MeOH-soaked kimwipes to clean the remained mCPBA crystal on the slide before imaging analysis.

**S6-3-4. Validation of spatial resolution**

The spatial resolution of DESI-MSI obtained from an mCPBA pretreated tissue section was validated here. We chose tissue sections of a mouse brain for validation as its DESI-MSI lipid imaging could show distinct regions^7^. Three serial coronal sections of a mouse brain were prepared. One was analyzed by DESI-MSI in full negative FT-MS mode, with a pixel-to-pixel resolution of 200 µm × 200 µm, and the original lipid distribution was obtained, where two distinct distributions of unsaturated lipids were revealed (**Fig S35a**). First, we showed that phosphatidylglycerol (PG) 18:1_16:0 was almost homogeneously distributed, with a slight decrease level in the cerebral deep while matter region. Second, sulfatide (ST) 24:1 was found to specifically locate in the white matter region, which was consistent with the previous report ^7^. Another section was subjected to H&E staining as histological reference.

Subsequently, the other serial section was pretreated by mCPBA with the proposed protocol, and analyzed by DESI-MSI in full FT-MS mode, where the distribution of the epoxidized lipids were investigated (**Fig S35b**). First, we revealed the distribution of the epoxidation product of PG 18:1_16:0, epoxy-PG 18:1_16:0, with a pattern in consistence with that before derivatization. Second, as efficient epoxidation was conducted using the proposed protocol, we spatially revealed diepoxy-ST 24:1, the epoxidation product of ST 24:1 with its two C=C bonds epoxidized. Remarkably, the imaging of diepoxy-ST 24:1 clearly indicated the fine structure of the white matter region that had been revealed by ST 24:1, implied that massive lateral diffusion was not observed, and thus the spatial resolution was retained after the mCPBA pretreatment.

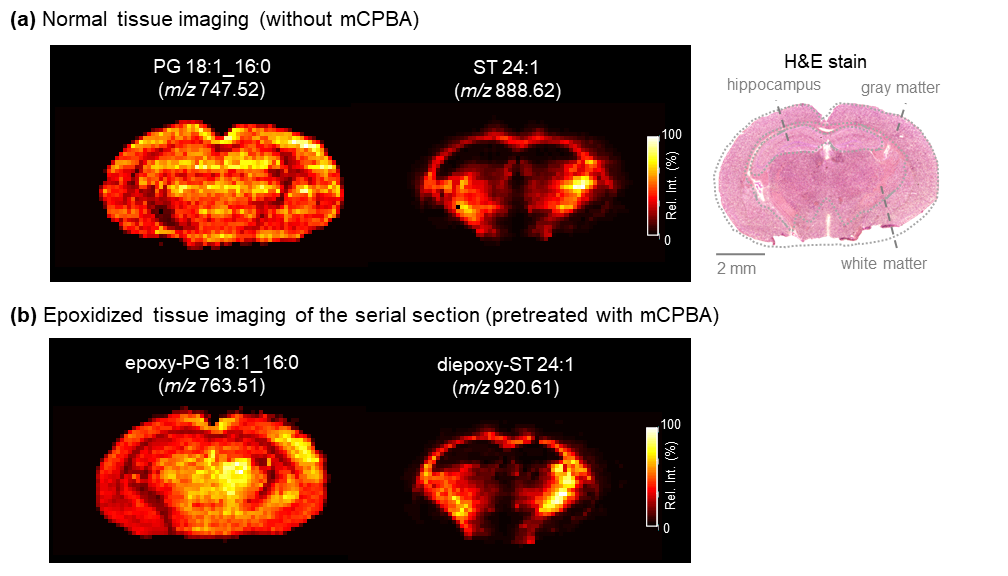

**Figure S35. Validation of DESI-MSI spatial resolution after *in situ* epoxidation using the proposed airbrushing protocol.**

**(a)** Normal DESI-MSI of the coronal section of a mouse brain showed distribution of two unsaturated lipids: PG 18:1_16:0 and sulfatide (ST) 24:1. The H&E staining of the serial section was also shown. **(b)** Another serial section was pretreated through the proposed *in situ* epoxidation method using an airbrush, and subjected to DESI-MSI (in full-MS mode). The epoxidation products of PG 18:1_16:0 and ST 24:1 were shown. The pixel-to-pixel resolution was set at 200 µm × 200 µm.

**S6-4-1. Mouse kidney sections**

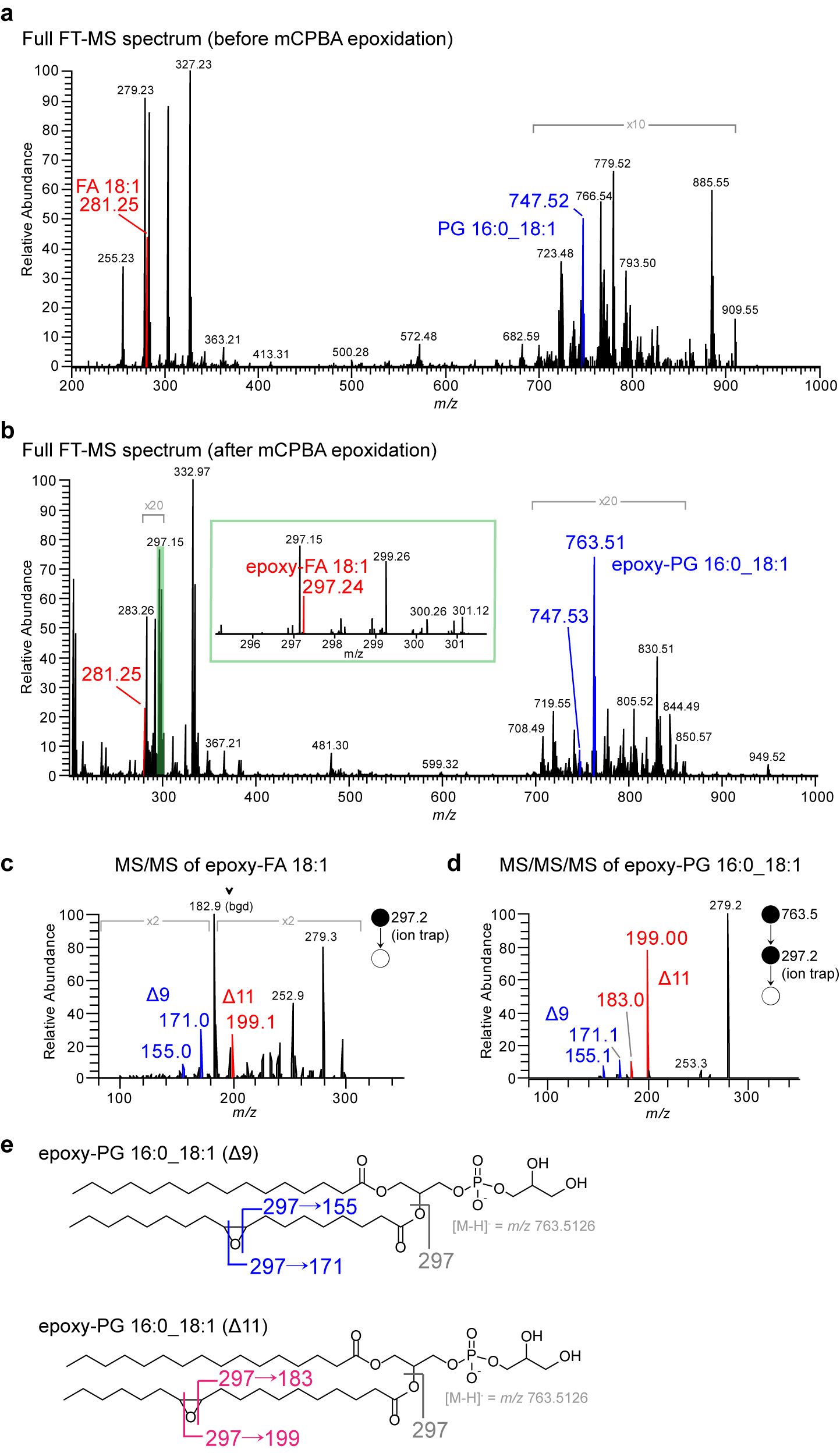

**Figure S36. MELDI-DESI-MS analysis of the mouse kidney section.**

The representative full FT-MS spectra of a mouse kidney section **(a)** prior to mCPBA pretreatment **(b)** after mCPBA pretreatment. From the epoxidized tissue section, we obtained **(c)** *in situ* IT-MS/MS spectrum of epoxidized FA 18:1 ([M-H]^-^ = *m/z* 297.24) **(d)** *in situ* IT-MS^3^ spectrum of epoxidized PG 16:0_18:1 ([M-H]^-^ = *m/z* 763.52), both indicating the presence of Δ9 and Δ11 isomers. **(e)** The CID scheme of epoxidized PG 160_18:1 C=C isomers.

**S6-4-2. Mouse metastatic lung sections**

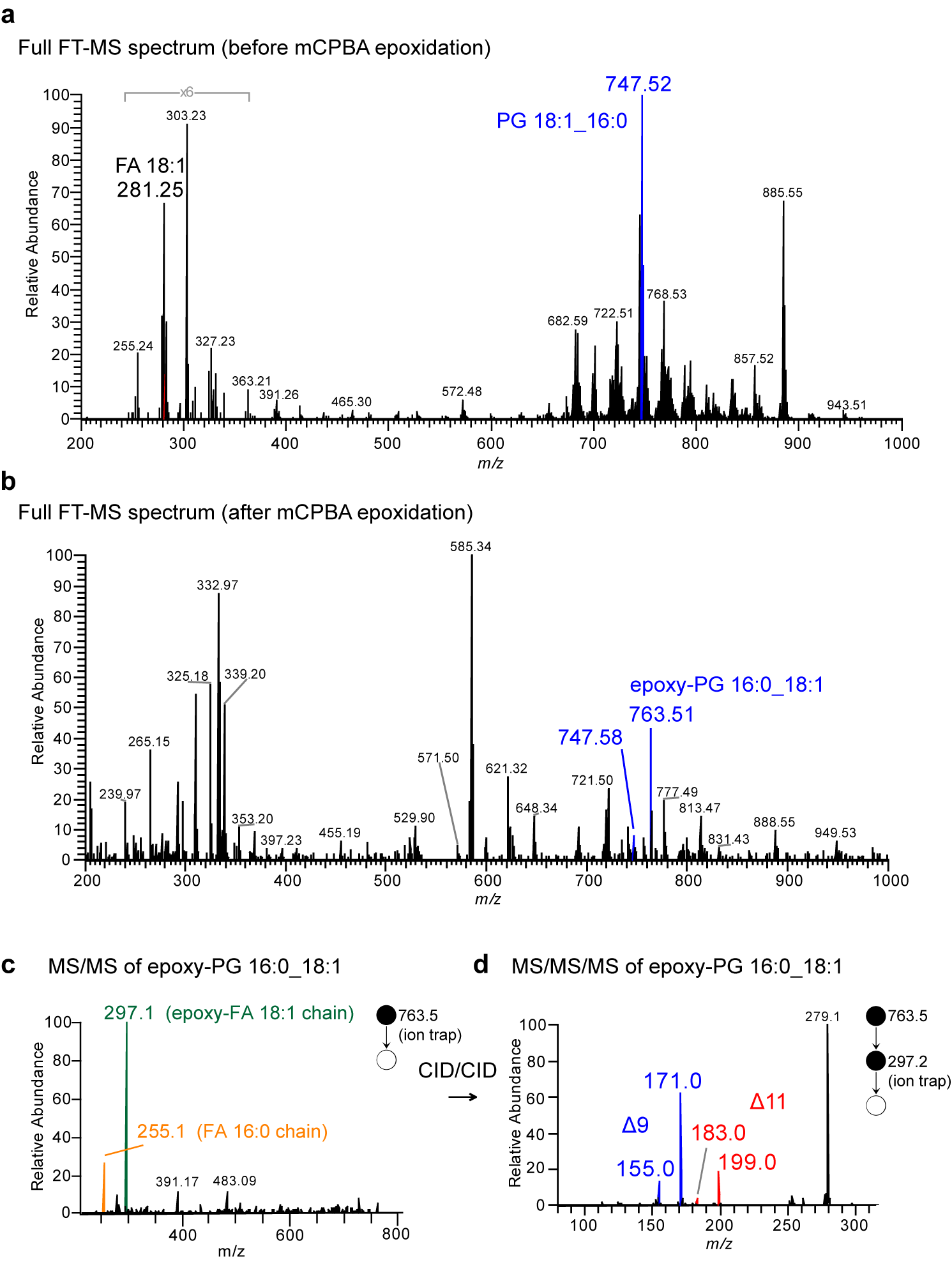

**Figure S36. MELDI-DESI-MS analysis of the mouse metastatic lung section.**

The representative full FT-MS spectra of a mouse metastatic lung section **(a)** prior to mCPBA pretreatment **(b)** after mCPBA pretreatment. From the epoxidized tissue section, the *in situ* IT-MS/MS spectrum of epoxidized PG 16:0_18:1 ([M-H]^-^ = *m/z* 763.52) was shown in (c), indicating the presence of an epoxidized FA 18:1 chain and a FA 16:0 chain. Subsequently, MS^3^ was further applied to the epoxy-FA 18:1 ion (*m/z* 297 in MS/MS), revealing the presence of Δ9 and Δ11 isomers.

**S7. Identification of C=C positions of unsaturated lipids in positive ionization mode**

In comparison to FAs and GPLs, lipids in other categories could be ionized in positive ion mode, which mainly include glycerolipid (GL), sterol lipids, sphingolipids. Also some GPLs such as PC could also be positively ionized. Here the capability of MELDI-MS/MS to the diagnosis of C=C positions of unsaturated lipids in positive ionization mode was investigated (**Fig. S37**).

(i) **Glycerolipid**

A common triacylglycerol (TAG) standard, triolein (TAG 18:1(9Z)/18:1(9Z)/18:1(9Z)) was used for demonstration. After mCPBA epoxidation, its product could be ionized as a sodium adduct ([triepoxyM+Na]^+^ = *m/*z 933.75) and the C=C diagnostic ions could be found in the MS/MS spectrum (**Fig. S37a**).

(ii) **Glycerophospholipid**

PC 18:1(9Z)/18:1(9Z) was epoxidzed to form an epoxy-PC, and subjected to FT-MS/MS analysis. The C=C position was resolved in the CID spectrum of its protonated adduct (**Fig. S37b**).

(iii) **Sphingolipid**

We found that C=C diagnostic ions could be probed in the MS/MS spectrum of an epoxidized unsaturated glycosphingolipid, as demonstrated by glucosylceramide (GlcCer) (d18:1/18:1(9Z)) (**Fig S37c**). However, we failed to find C=C diagnostic ions in other sphingolipid subclasses, such as ceramide (Cer) and sphingomyelin (SM).

(iv) **Sterol lipid**
     Cholesteryl ester (CE) is one of the subclasses of sterol lipid in which C=C positional isomers are present. We showed that for CE 18:1 (9Z), its epoxidation product, epoxy-CE 18:1 (with an epoxide at the original C=C bond in the fatty acyl chain), was ionized as a sodium adduct ([epoxyM+Na]^+^ = *m/*z 764.56) in positive ionization mode (**Fig S37d**). Subsequent MS/MS revealed a abundant fragment of *m/z* 321.24, corresponding to an epoxy-FA 18:1 sodium ion, while annotation of C=C position in MS2 level was not allowed. Applying the MS^3^ strategy similar to that for GPLs in negative ion mode, the Δ9 C=C position was then identified by an aldehyde/alkene diagnostic ion pair (**lower panel in Fig S37d**).

Finally, the overall performance of MELDI for each lipid category was summarized in **Table S14**. In our opinion, the C=C isomer of CE was the most promising target for future analysis in positive ionization mode as its epoxide yielded the most abundant C=C diagnostic ions, yet for other lipid classes, longer MS/MS acusition time or analytes of higher concentration were recommended to ensure the MS/MS spectral quality for C=C diagnosis. To sum up, identification of C=C positions using MELDI was also applicable for many classes of positively ionized lipids.

**Table S14. Performance of MELDI-MS/MS in pinpointing C=C positions of unsaturated lipids of difference classes.**

| Lipid class | Ionization mode / MS^n^ | Studied subclasses | Abundance of C=C diagnostic ions |
| --- | --- | --- | --- |
| Fatty acids (FA) | (-) MS/MS | MOFA, PUFA | High |
| Glycerophospholipid (GPL) | (-) MS/MS/MS  (+) MS/MS | PS, PC, PA, PE, PC, PI  PC | High  Low |
| Sterol Lipids | (+) MS/MS/MS | Cholesteryl ester (CE) | High |
| Glycerolipids (GL) | (+) MS/MS | Triacylglycerol (TAG) | Low |
| Sphingolipids | (+) MS/MS | Glucosylceramide (GlcCer)  Ceramide (Cer)  Sphingomyelin (SM) | Low  N/A  N/A |

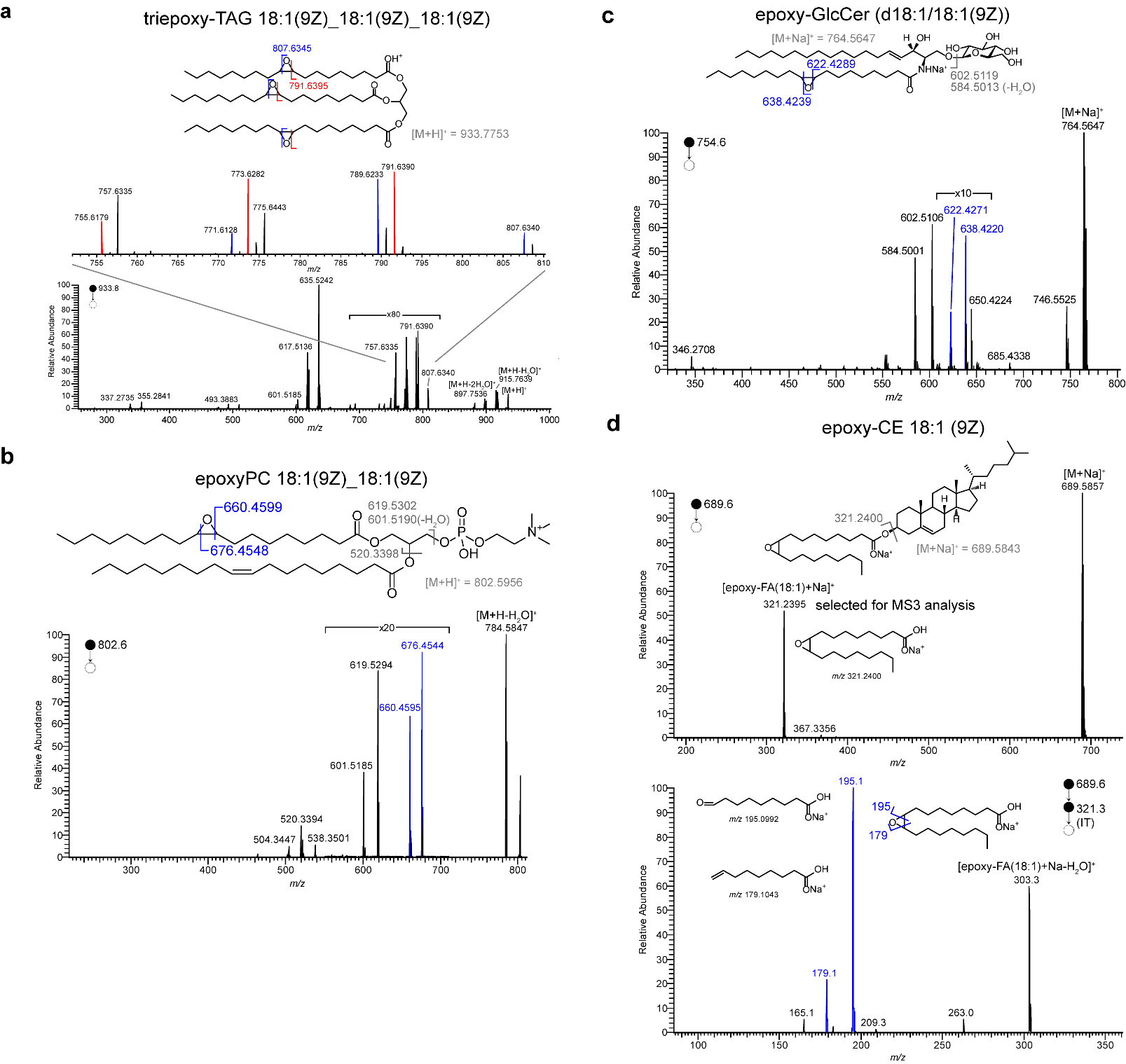

**Figure S37. Pinpointing C=C positions of unsaturated lipids by MELDI-MS^n^ in positive ionization mode.**

The MS/MS (or MS^n^) spectra of the epoxidized lipids in positive ionization mode were shown: **(a)** Fully epoxidized TAG 18:1(9Z)/18:1(9Z)/18:1(9Z) **(b)** mono-epoxidized PC 18:1(9Z)/18:1(9Z) **(c)** mono-epoxidized glucosylceramide (GlcCer) d18:1/18:1 (9Z) **(d)** mono-epoxidized cholesteryl ester (CE) 18:1 (9Z). The spectra were acquired in high resolution FT mode except for the MS^3^ spectrum of the epoxy-CE was acquired in IT mode to ensure the spectral quality.

**S8. Compatibility to commercial mass spectrometers**

MELDI was capable of being integrated to a wide variety of mass spectrometers coupled with difference ionization sources. Generally, when the epoxidized unsaturated lipids could be ionized, they could undergo subsequent CID-based tandem mass spectrometric experiments to reveal the C=C positions. In the main text, we had demonstrated the capability in Orbitrap Elite Mass Spectrometry (Thermo) coupled to a common (LC-)ESI source as well as an ambient ionization source, DESI. Here we further demonstrated MELDI in versatile approaches:

(i) Orbitrap Elite Mass Spectrometer coupled to a paper spray ionization (PSI) source (**Fig. S38**)

First, 10 µL of serum was direct deposited onto a well-cutted filter paper without any pretreatment, dried, and analyzed by PSI-FT-MS in negative ionization mode. Here a solvent of MeOH:CCl_4_:NH_4_OH = 90:10:0.1 was chosen to alleviate corona discharge in negative PSI, and a solvent flow rate was 15 µL/min. The obtained FT-MS spectrum of a non-derivatized serum sample showed abundant signals of unsaturated fatty acids (**Fig. S38b**). Second, 10 µL of mCPBA (200 mM in methanol) was also deposited onto the paper for on-site epoxidation, dried, and analyzed again. Remarkably, the resulting full FT-MS spectrum showed abundant signals of epoxides, including epoxy-FA 16:1, epoxy-FA 18:1, and epoxy-FA 18:2 (**Fig. S38c**), and their C=C positions were revealed by subsequent MS/MS analysis (**Fig. S38d-f**). Remarkably, the overall process took less than 1 minute.

(ii) Orbitrap Elite Mass Spectrometer coupled to an atmospheric pressure chemical ionization (APCI) source (**Fig. S39**)

Unsaturated lipid standards, including FA 18:1 (9Z), FA 18:1 (11Z) and PG 18:1(9Z)/18:1(9Z), were derivatized by mCPBA in organic solution, subjected to APCI, and their C=C positions were identified by the diagnostic ions in the CID spectrum (or CID/CID for PG).

(iii) MALDI-TOF/TOF Mass Spectrometer (Autoflex speed, Bruker Daltonics) (**Fig. S40**)

Three unsaturated lipid standards, FA 18:1 (9Z), FA 18:1 (11Z), and PG 18:1(9Z)/18:1(9Z), were tested. Each sample was derivatized by mCPBA and analyzed by MALDI-TOF/TOF, and the C=C diagnostic ions were revealed. We noted that the ionization of FA was facilitated by a nanogold-coated mesoporous silica substrate ^8^, and 9-aminoacridine (9-AA) was used as matrix to assist ionization of PG.

These results suggest that as mCPBA is commercially available, MELDI does provide a handy way for diagnosing C=C positional isomers for a mass spectrometer that is equipped with CID.

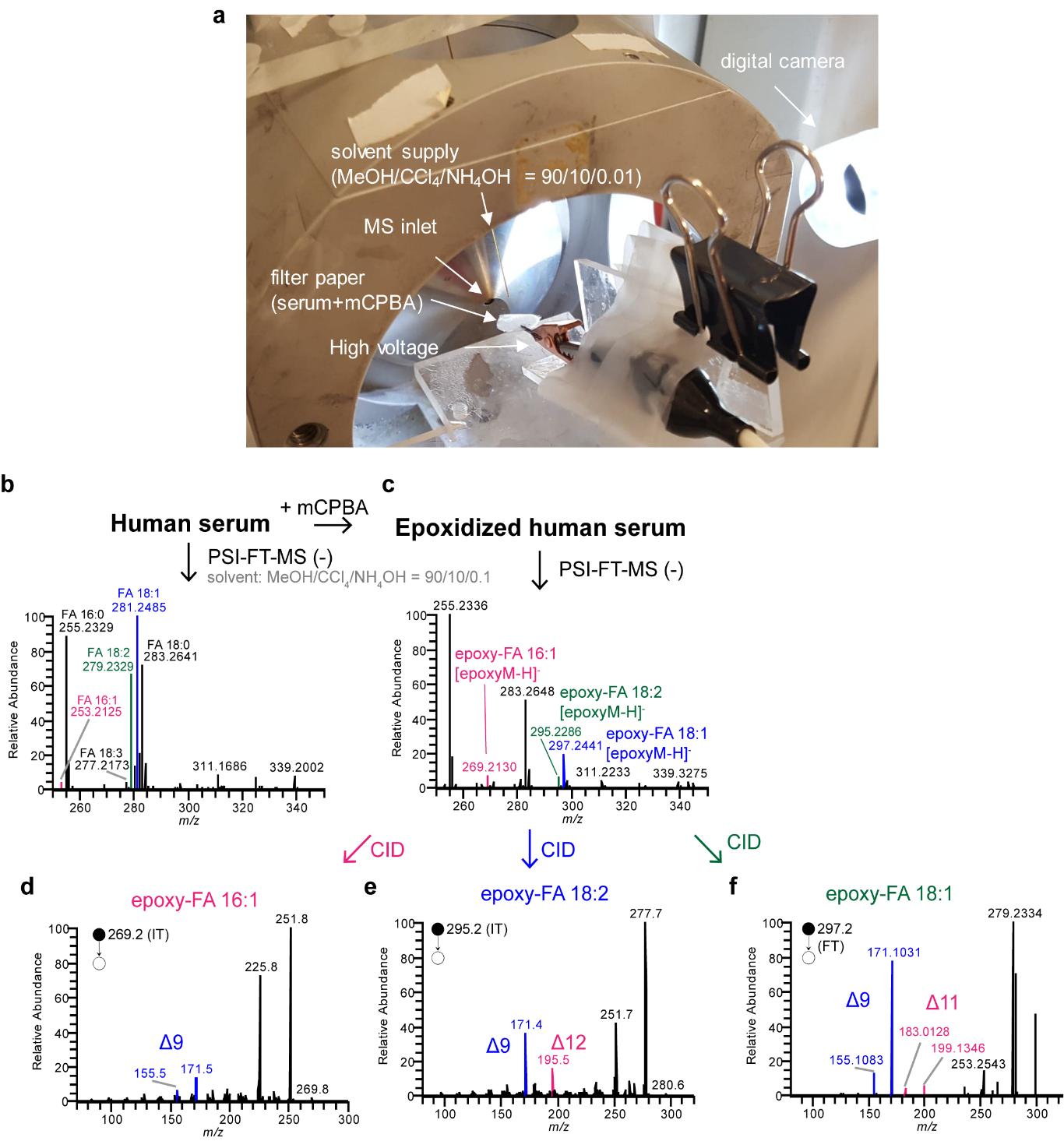

**Figure S38. MELDI-PSI-MS/MS for rapid diagnosis of C=C isomers**.

Orbitrap Elite Mass Spectrometer was coupled with a paper spray ionization source for diagnosis of C=C positional isomers. **(a)** The instrumental setup of MELDI-PSI-MS. The full FT-MS spectra of a human serum sample prior and after on-site epoxidation using mCPBA were shown in **(b)** and **(c)**. **(d-f)** MS/MS spectra of the epoxidized FAs, where C=C diagnostic ions were indicated.

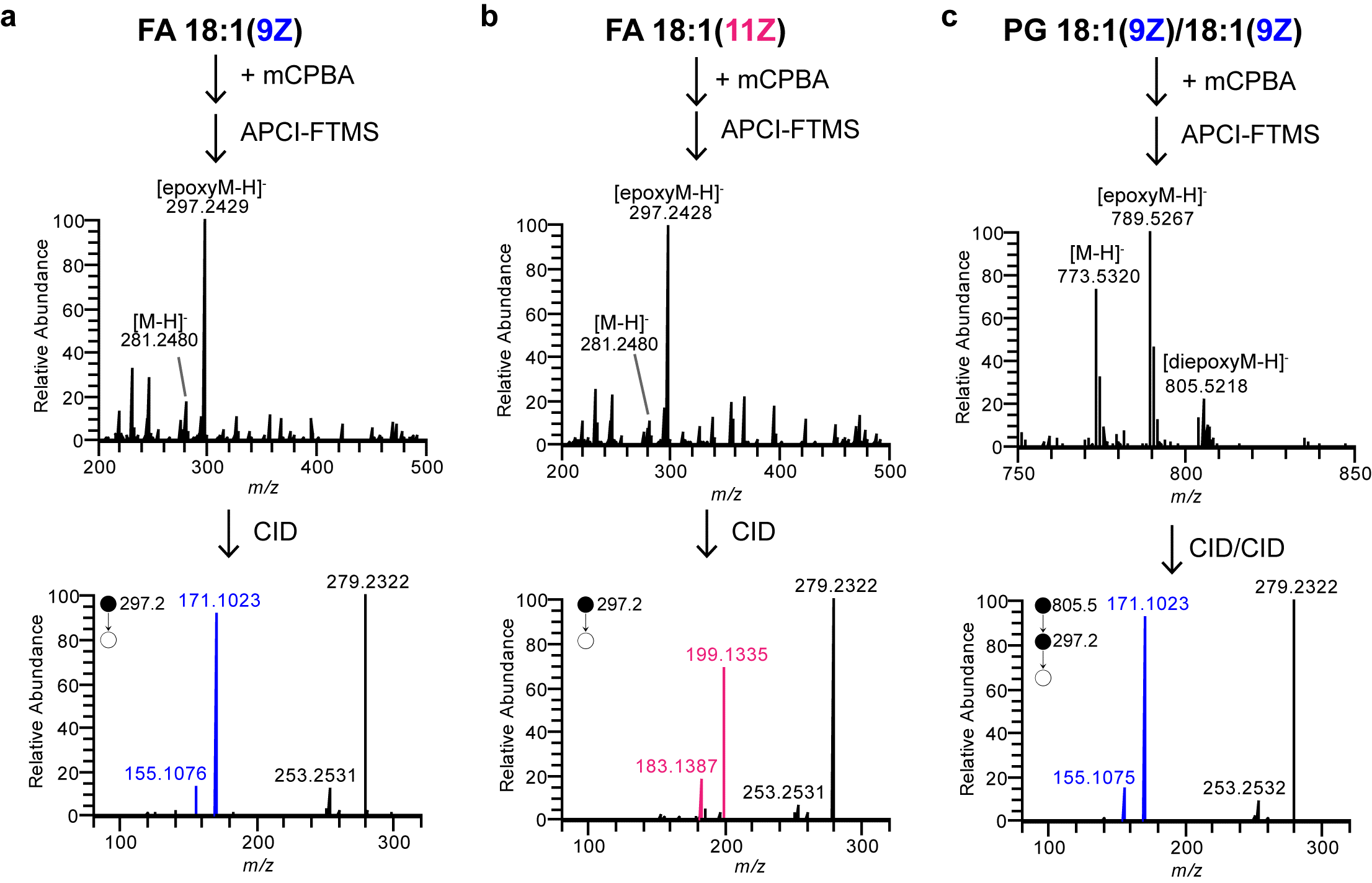

**Figure S39. MELDI-APCI-MS/MS for identification of C=C isomers**.

Orbitrap Elite Mass Spectrometer was coupled with an APCI source for diagnosis of C=C positional isomers. The APCI-MS and APCI-MS^n^ spectra of the epoxidized unsaturated lipid standards were shown: **(a)** FA 18:1 (9Z) **(b)** FA 18:1 (11Z) **(c)** PG 18:1(9Z)/18:1(9Z). The C=C diagnostic ions were colored.

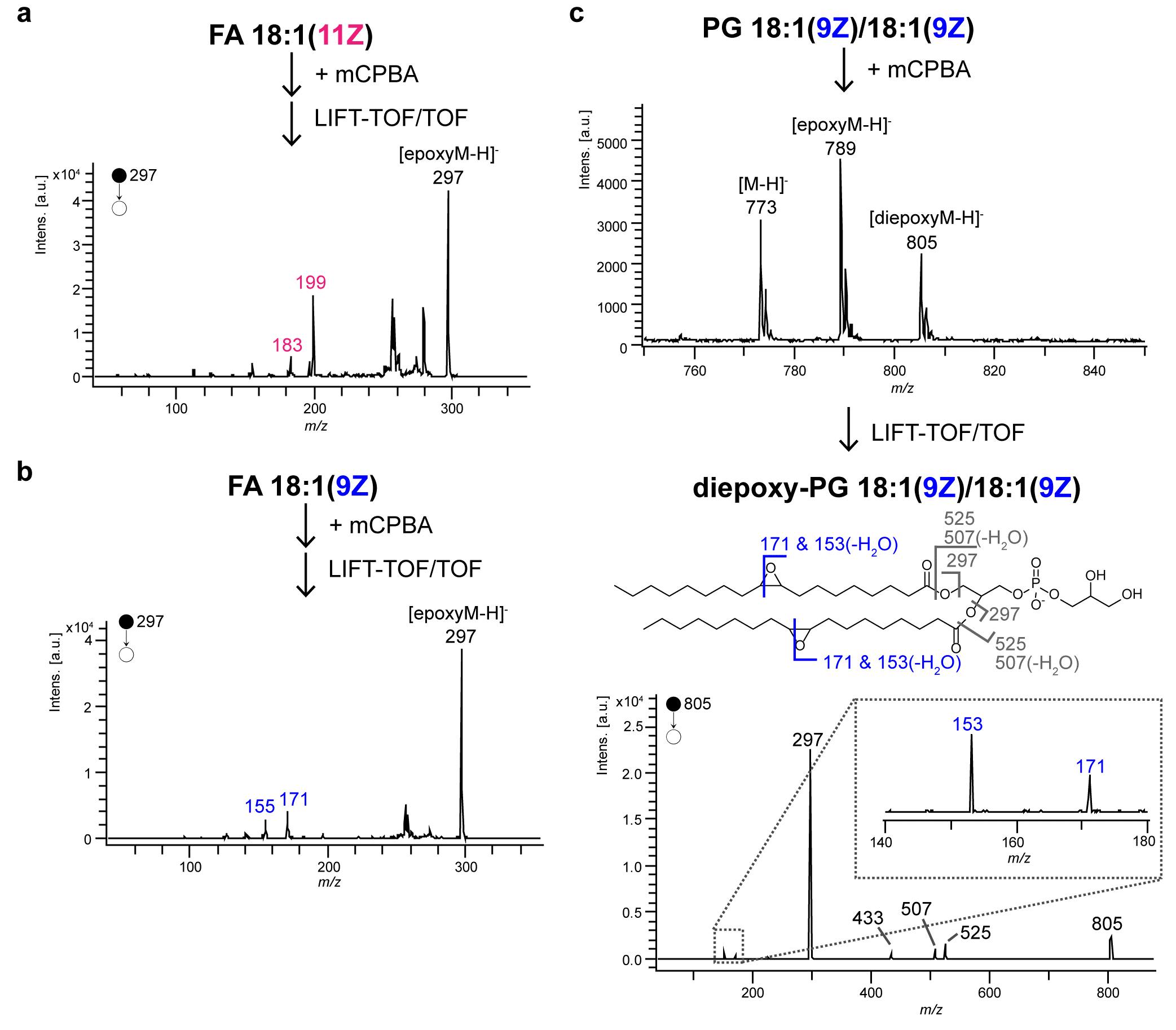

**Figure S40. MELDI-MALDI-TOF/TOF for identification of C=C positions**.

MALDI-TOF/TOF Mass Spectrometer was used to identify C=C positions in unsaturated lipids using MELDI. The LIFT-TOF/TOF spectra were shown: **(a)** epoxy-FA 18:1 (11Z) **(b)** epoxy-FA 18:1 (9Z) **(c)** diepoxy-PG 18:1(9Z)/18:1(9Z).

**S9. Summary of the C=C diagnostic ions of the mono-epoxidized unsaturated lipids**

**Table S14. Table of the MS/MS diagnostic ions of the mono-epoxidized unsaturated lipids.**

The identified diagnostic ions of the mono-epoxidized lipids, represented by their theoretical *m/z* values. For polyunsaturated lipids with multiple diagnostic ions, the most abundant ions were shown in bold.

|  | | **Types of fragments** | **Carboxyl-end fragments** | | | | **Methyl-end fragments** | | | |
| --- | --- | --- | --- | --- | --- | --- | --- | --- | --- | --- |
|  |  | **Bond cleavage**  **Position upon CID** | 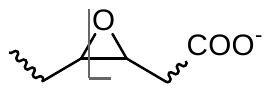 | 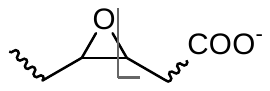 | 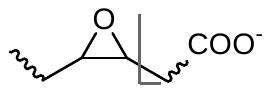 | 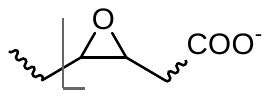 | 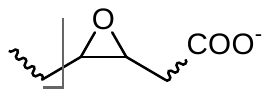 | 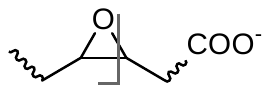 | 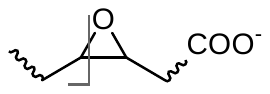 | 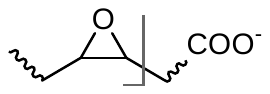 |
| **FA species** | **ω-family** | **Epoxide position** |  |  |  |  |  |  |  |  |
| Mono-unsaturated FA | | Δ6 | 129.0557 | 113.0000 |  |  |  |  |  |  |
|  |  | Δ7 | 143.0714 | 127.0765 |  |  |  |  |  |  |
|  |  | Δ8 | 157.0870 | 141.0921 |  |  |  |  |  |  |
|  |  | Δ9 | 171.1027 | 155.1078 |  |  |  |  |  |  |
|  |  | Δ10 | 185.1183 | 169.1234 |  |  |  |  |  |  |
|  |  | Δ11 | 199.1340 | 183.1391 |  |  |  |  |  |  |
|  |  | Δ12 | 213.1496 | 197.1547 |  |  |  |  |  |  |
|  |  | Δ13 | 227.1653 | 211.1704 |  |  |  |  |  |  |
|  |  | Δ14 | 241.1809 | 225.1860 |  |  |  |  |  |  |
|  |  | Δ15 | 255.1966 | 239.2017 |  |  |  |  |  |  |
|  |  | Δ16 | 269.2122 | 253.2173 |  |  |  |  |  |  |
|  |  | Δ17 | 283.2279 | 267.2330 |  |  |  |  |  |  |
| FA 18:2 | ω-6 (9, 12) | Δ9 | 171.1027 |  |  |  |  |  |  |  |
|  |  | Δ12 |  | 195.1391 |  |  |  |  |  |  |
|  | ω-7 (8,11) | Δ8 | 157.087 |  |  |  |  |  |  |  |
|  |  | Δ11 |  | 181.1234 |  |  |  |  |  |  |
|  | ω-9 (6, 9) | Δ6 | 129.0557 |  |  |  |  |  |  |  |
|  |  | Δ9 |  | 153.0291 |  |  |  |  |  |  |
| FA 18:3 | ω-6 (6, 9, 12) | Δ6 | **129.0557** | 113.0608 |  | 141.0557 |  |  |  | 191.1441 |
|  |  | Δ9 | 169.087 | **153.0921** |  |  |  |  |  |  |
|  |  | Δ12 |  | 193.1234 |  |  |  |  |  |  |
|  | ω-7 (5, 8, 11) | Δ5 |  |  |  |  |  | 193.1598 |  |  |
|  |  | Δ8 |  |  |  |  |  | 153.1285 |  |  |
|  |  | Δ11 |  | 179.1078 |  |  |  |  |  |  |
|  | ω-3 (9, 12, 15) | Δ9 | 171.1027 |  |  |  |  |  |  |  |
|  |  | Δ12 | **211.1340** | 195.1398 |  |  |  |  |  |  |
|  |  | Δ15 |  | 235.1704 |  |  |  |  |  |  |
| FA 18:4 | ω-3 (6, 9, 12, 15) | Δ6 | N/A |  |  |  |  |  |  |  |
|  |  | Δ9 |  | 153.0921 |  |  |  |  |  |  |
|  |  | Δ12 | 209.1183 | **193.1234** | 181.1234 |  |  |  |  |  |
|  |  | Δ15 |  | 233.1547 |  |  |  |  |  |  |
| FA 20:2 | ω-9 (8, 11) | Δ8 | 157.087 |  |  |  |  |  |  |  |
|  |  | Δ11 |  | 181.1234 |  |  |  |  |  |  |
|  | ω-6 (11, 14) | Δ11 | 199.134 |  |  |  |  |  |  |  |
|  |  | Δ14 |  | 223.1704 |  |  |  |  |  |  |
|  | ω-7 (10, 13) | Δ10 | 209.1547 |  |  |  |  |  |  |  |
|  |  | Δ13 |  | 185.1183 |  |  |  |  |  |  |
|  | conj. ω-6 (12, 14) | Δ12 |  | 197.1547 |  |  |  |  |  |  |
|  |  | Δ14 | 239.1653 |  |  |  |  |  |  |  |
|  | conj. ω-7 (11, 13) | Δ11 |  | 183.1391 |  |  |  |  |  |  |
|  |  | Δ13 | 225.1496 |  |  |  |  |  |  |  |
| FA 20:3 | ω-6 (8, 11, 14) | Δ8 | 157.084 |  |  |  |  |  |  |  |
|  |  | Δ11 | 197.1183 |  |  |  |  |  |  |  |
|  |  | Δ14 |  | 221.1547 |  |  |  |  |  |  |
|  | ω-9 (5, 8, 11) | Δ5 |  |  |  |  |  | 221.1911 |  |  |
|  |  | Δ8 |  |  |  |  |  | 181.1598 |  |  |
|  |  | Δ11 |  | 179.1078 |  |  |  |  |  |  |
|  | ω-3 (11, 14, 17) | Δ11 | 199.134 |  |  |  |  |  |  |  |
|  |  | Δ14 | 239.1653 |  |  |  |  |  |  |  |
|  |  | Δ17 |  | 263.2017 |  |  |  |  |  |  |
| FA 20:4 | ω-6 (5, 8, 11, 14) | Δ5 |  |  |  |  | **191.1805** |  |  | 219.1754 |
|  |  | Δ8 | 155.0714 |  |  |  |  |  |  |  |
|  |  | Δ11 |  | 179.1079 |  |  |  |  |  |  |
|  |  | Δ14 |  | 219.1391 |  |  |  |  |  |  |
|  | ω-3 (8, 11, 14, 17) | Δ8 | 157.087 |  |  |  |  |  |  |  |
|  |  | Δ11 | N/A |  |  |  |  |  |  |  |
|  |  | Δ14 | N/A |  |  |  |  |  |  |  |
|  |  | Δ17 |  | 261.186 |  |  |  |  |  |  |
| FA 20:5 | ω-3 (5, 8, 11, 14, 17) | Δ5 |  |  |  |  | **189.1649** | 217.1598 |  |  |
|  |  | Δ8 | 155.0714 |  |  |  |  |  |  |  |
|  |  | Δ11 | 195.1027 | **179.1078** |  |  |  |  |  |  |
|  |  | Δ14 | 235.134 | 219.1391 | **207.1391** |  |  |  |  |  |
|  |  | Δ17 |  | 259.1704 |  |  |  |  |  |  |
| FA 22:2 | ω-9 (10, 13) | Δ10 | 185.1183 |  |  |  |  |  |  |  |
|  |  | Δ13 |  | 209.1547 |  |  |  |  |  |  |
|  | ω-6 (13, 16) | Δ13 | 227.1653 |  |  |  |  |  |  |  |
|  |  | Δ16 |  | 251.2017 |  |  |  |  |  |  |
|  | ω-7 (12, 15) | Δ12 | 213.1496 |  |  |  |  |  |  |  |
|  |  | Δ15 | 253.1809 | 237.186 |  |  |  |  |  |  |
|  | ω-10 (9, 12) | Δ9 | 171.1027 |  |  |  |  |  |  |  |
|  |  | Δ12 |  | 195.1391 |  |  |  |  |  |  |
| FA 22:3 | ω-9 (7 ,10, 13) | Δ7 | **143.0714** |  |  |  |  |  |  |  |
|  |  | Δ10 | **183.1027** |  |  |  |  |  |  |  |
|  |  | Δ13 |  | 207.1391 |  |  |  |  |  |  |
|  | ω-6 (10 ,13, 16) | Δ10 | 185.1183 |  |  |  |  |  |  |  |
|  |  | Δ13 | 225.1496 |  |  |  |  |  |  |  |
|  |  | Δ16 |  | 249.186 |  |  |  |  |  |  |
|  | ω-7 (11, 14, 17) | Δ9 | 171.1027 |  |  |  |  |  |  |  |
|  |  | Δ12 | 211.134 |  |  |  |  |  |  |  |
|  |  | Δ15 |  | 235.1704 |  |  |  |  |  |  |
| FA 22:4 | ω-6 (7, 10, 13, 16) | Δ7 | 143.0714 |  |  | 157.0874 |  |  | 203.1805 |  |
|  |  | Δ10 | 183.1027 | 167.1078 |  |  |  |  |  |  |
|  |  | Δ13 | 223.134 | 207.1391 | **195.1391** |  |  |  |  |  |
|  |  | Δ16 |  | 247.1704 |  |  |  |  |  |  |
|  | ω-3 (10, 13, 16, 19) | Δ10 | 185.1183 |  |  |  |  |  |  |  |
|  |  | Δ13 | N/A |  |  |  |  |  |  |  |
|  |  | Δ16 | N/A |  |  |  |  |  |  |  |
|  |  | Δ19 |  | 289.2173 |  |  |  |  |  |  |
| FA 22:5 | ω-3 (7, 10, 13, 16, 19) | Δ7 | **143.0714** |  |  |  |  |  | 201.1649 |  |
|  |  | Δ10 | **183.1027** | 167.1078 |  |  |  |  |  |  |
|  |  | Δ13 |  | 207.1391 |  |  |  |  |  |  |
|  |  | Δ16 | 263.1653 | 247.1704 | **235.1704** |  |  |  |  |  |
|  |  | Δ19 |  | 287.2017 |  |  |  |  |  |  |
|  | ω-6 (4, 7, 10, 13, 16) | Δ4 | N/A |  |  |  |  |  |  |  |
|  |  | Δ7 |  |  |  |  | 191.1805 |  |  |  |
|  |  | Δ10 | N/A |  |  |  |  |  |  |  |
|  |  | Δ13 |  | 205.1234 |  |  |  |  |  |  |
|  |  | Δ16 |  | 245.1547 |  |  |  |  |  |  |
| FA 22:6 | ω-3 (4, 7, 10, 13, 16, 19) | Δ4 |  |  |  |  |  |  | 241.1962 |  |
|  |  | Δ7 |  |  |  |  | 189.1649 |  | **201.1649** |  |
|  |  | Δ10 | 181.087 |  |  |  |  |  | 161.1336 |  |
|  |  | Δ13 | 221.1183 | **205.1234** |  |  |  |  |  |  |
|  |  | Δ16 | 261.1496 | 245.1547 | **233.1547** |  |  |  |  |  |
|  |  | Δ19 |  | 285.186 |  |  |  |  |  |  |
| FA 24:2 | ω-6 (15, 18) | Δ15 | 255.1966 |  |  |  |  |  |  |  |
|  |  | Δ18 |  | 279.223 |  |  |  |  |  |  |
|  | ω-9 (12, 15) | Δ12 | 213.1496 |  |  |  |  |  |  |  |
|  |  | Δ15 | **253.1809** | 237.186 |  |  |  |  |  |  |
| FA 24:6 | ω-3 (9, 12, 15, 18, 21) | Δ6 |  |  |  |  | 229.10962 |  | **243.2118** |  |
|  |  | Δ9 | 169.087 |  |  |  |  |  |  |  |
|  |  | Δ12 | 209.1183 | **193.1234** |  |  |  |  |  |  |
|  |  | Δ15 | **249.1496** | 233.1547 |  |  |  |  |  |  |
|  |  | Δ18 | 289.1809 | 273.186 | 261.186 |  |  |  |  |  |
|  |  | Δ21 |  | 313.2173 |  |  |  |  |  |  |
